# Relying on the relationship with known disease-causing variants in homologous proteins to predict pathogenicity of *SORL1* variants in Alzheimer’s disease

**DOI:** 10.1101/2023.02.27.524103

**Authors:** Olav M. Andersen, Giulia Monti, Anne Mette G. Jensen, Matthijs de Waal, Marc Hulsman, Johan G. Olsen, Henne Holstege

**Affiliations:** Department of Biomedicine, Aarhus University, Høegh-Guldbergs Gade 10, DK8000 AarhusC, Denmark; Alzheimer Center Amsterdam, Department of Neurology, Amsterdam Neuroscience, Vrije Universiteit Amsterdam, Amsterdam UMC, Amsterdam, The Netherlands; Department of Biology, University of Copenhagen, Ole Maaløes Vej 5, DK2200, Copenhagen, Denmark

**Keywords:** SORLA, SORL1-associated Alzheimer’s disease, modular receptor, pathogenic variants, disease-mutations domain-mappping

## Abstract

*SORL1* encodes the retromer-associated receptor SORLA that functions in endosomal recycling. Rare variants in *SORL1* have been associated with Alzheimer’s disease (AD) and rare pathogenic variants are estimated to occur in up to 2.75% of early onset AD patients and in 1.5% of unrelated late onset AD patients. While truncation mutations are observed almost exclusively in AD patients, it is currently unknown which among the hundreds of rare missense variants identified in *SORL1*, are pathogenic. Here we address this question by relying on SORLA’s distinct molecular architecture. First, we completed a structure-guided sequence alignment for all the protein domains. Next, we identified proteins that contain domains homologous to those of SORLA, which include pathogenic variants for monogenic diseases. We identified the analogous domain positions of these variants in the SORLA protein sequence and showed that variants in these positions similarly impair *SORL1*, and lead to AD. Together, our findings represent a comprehensive compendium on SORLA protein variation and functional effects, which allowed us to prioritize *SORL1* genetic variants into high or moderate priority mutations. We envision that this compendium will be used by clinical geneticists for assessing variants they identify in patients, allowing further development of diagnostic procedures and patient counseling strategies. Utimately, this compendium will inform investigations into the molecular mechanisms of endosomal recycling which will support the development of therapeutic treatment strategies for *SORL1* variant-carrying patients.

## INTRODUCTION

SORLA, the sortilin-related receptor with type-A repeats, is also known as LR11 or by its gene name *SORL1*. Since 2007, a multitude of studies associated both common and rare variants in the *SORL1* gene with Alzheimer’s disease (AD) ^1–4^, and GWAS studies have recurrently shown that a non-coding single nucleotide polymorphism (SNP) near the *SORL1* gene (rs11218343) is associated with significant modified risk for the common, late onset form of AD (LOAD) ^5–7^ while a second SNP (rs74685827) was recently identified to independently associate with increased risk of AD ^8^. Furthermore, several exome sequencing studies identified a large number of potentially deleterious *SORL1* missense variants ^9–11^. In fact, a recent exome sequencing study that compared rare variants between AD cases and controls indicated that of all genes in the human genome, the *SORL1* gene encompassed the most pathogenic variants ^12^. GnomAD currently lists almost 3,500 previously identified *SORL1* variants ^13^, and only a subset of these are risk-increasing or possibly causative of AD. An estimated 2.75% of non-related early onset AD (EOAD) patients and 1.5% of non-related LOAD cases carry such a pathogenic variant, while an even larger fraction of AD patients carries a rare *SORL1* variant with lower predicted pathogenicity ^12^. This is a much larger fraction than the <1% of EOAD cases affected by *PSEN1, PSEN2* or *APP* carriers combined ^14^. Rare loss-of-function variants in *SORL1* (i.e. truncating nonsense, frameshift or splice variants) are observed almost exclusively in AD cases suggesting that *SORL1* is haploinsufficient ^9,10,15^. Carrying a loss-of-function variant was shown to lead to a >40-fold increased risk of EOAD and to a >10-fold increased risk for LOAD ^12^.

Next to the evidence presented by genetic studies, additional evidence implicating the role of SORLA in AD-associated mechanisms comes from studies showing that loss of SORLA function triggers hallmark pathologies of AD. SORLA is a type-1 transmembrane receptor that has long been known to function with the retromer endosomal trafficking complex ^16,17^. Recent studies have clarified how SORLA binds with retromer, forming a scaffold that stabilizes the highly dynamic tubules through which cargo is transported out of endosomes ^18,19^. SORLA deficiency in human and nonhuman cell lines and in different animal models, which serves as a model for *SORL1* haploinsufficiency, as observed in individuals who carry a truncating *SORL1* variant ^9^ is shown to impair endosomal recycling and to trigger hallmark features of AD’s amyloid, tau, and synaptic pathologies ^20–22^.

While the aggregate effect on AD risk of rare loss-of-function variants is well described ^12,23^, >90% of the observed *SORL1* variants are missense variants and it is currently unknown which among these hundreds of variants are pathogenic. However, the effect on AD of each missense variant is unique ^10^: while the common missense variant E270K does not associate with AD in GWAS ^8^, a recent study suggests that the rare D1545V missense variant leads to an autosomal dominant inheritance pattern of AD in an Icelandic family ^24^. However, for most carriers of *SORL1* variants, pedigrees are not available. In cases for which pedigrees were available, they included at most four generations of affected family members, with a variable age at onset per affected family member. This complicates the evaluation of variant penetrance such that alternative approaches to assess pathogenicity are warranted.

Here, we address this outstanding question by relying on the distinct molecular architecture of SORLA (**Fig. 1**). SORLA is a mosaic protein, which comprises functional domains of both the low-density lipoprotein receptor (LDLR) and the vacuolar protein sorting-10 protein (VPS10p) families, but almost one third of the SORLA protein cannot be assigned to a specific protein family (**Fig. 2**). The full SORLA sequence spans 2,214 residues, and after post-translational removal of a 28 amino acid signal peptide (SP), the processed human receptor contains 2,186 amino acids encoded by 48 exons: a pro-peptide (ProP), a VPS10p-domain with an adjacent 10CC region, a YWTD-repeated domain, an epidermal growth factor (EGF)-domain, clusters of complement-type repeat (CR)- domains and fibronectin-type III (3Fn) domains, a transmembrane (TM) domain, and a cytoplasmic tail (**Fig. 1**) ^25^. Whereas the CR-cluster and VPS10p-domain have been shown to interact directly with multiple ligands including APP and Amyloid-β, respectively, the binding partners and functions of the YWTD- and 3Fn-domains are largely unknown ^26^.

**Figure 1.**
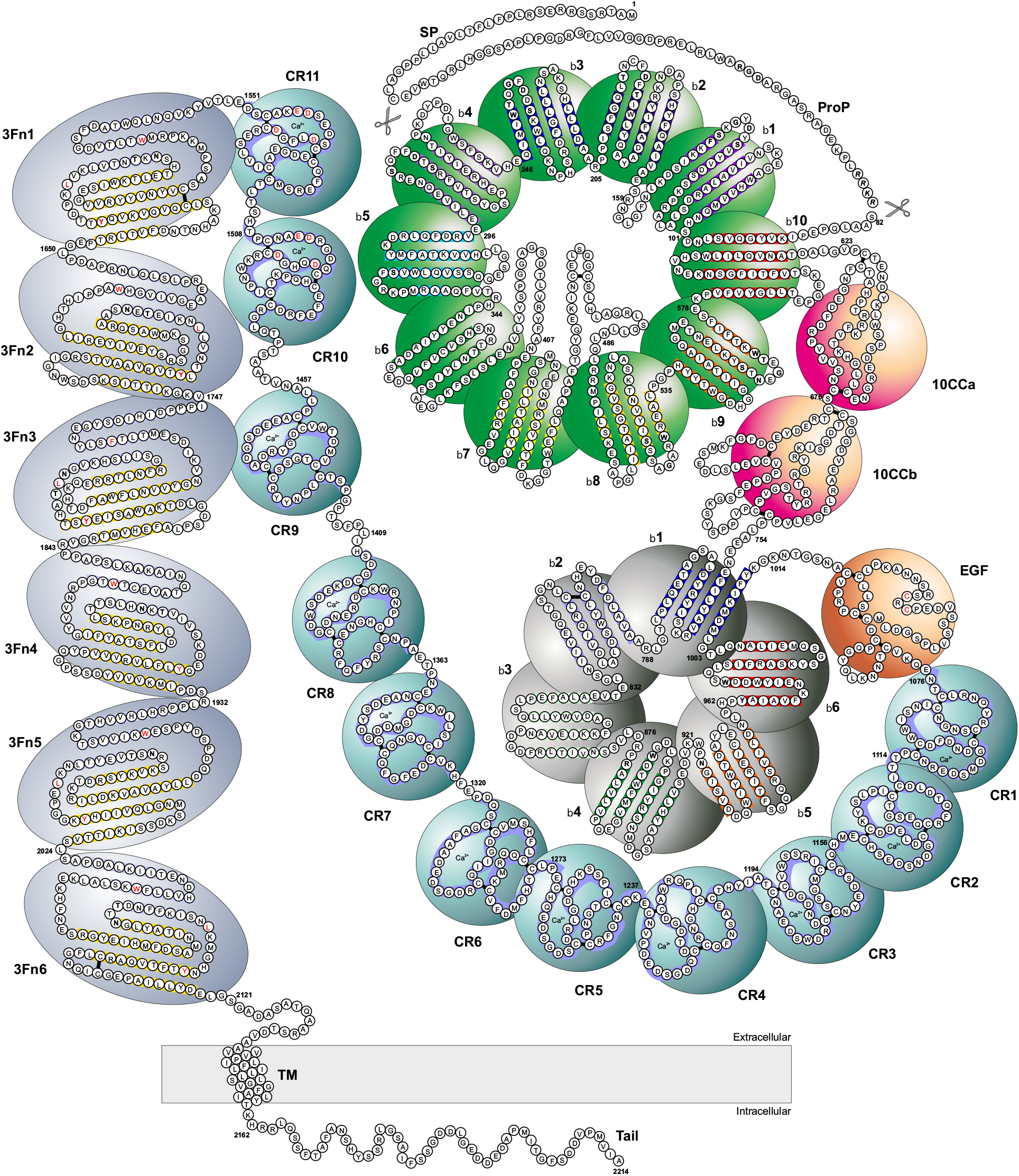
Diagram with 2,214 amino acids of human SORLA. Diagram representing the entire 2,214 amino acid sequence as ball presentations. The foldings of Individual domains are represented with the color codes as in the figures that display these domains in the Supplemental Information. Scissors at positions 28 and 81 indicate positions for cleavage by signal peptidase and Furin just after the signal- and the pro-peptides, respectively. A similar diagram will be made available at the alzforum.org as an interactive resource to provide detailed information on all known *SORL1* variants.

**Figure 2.**
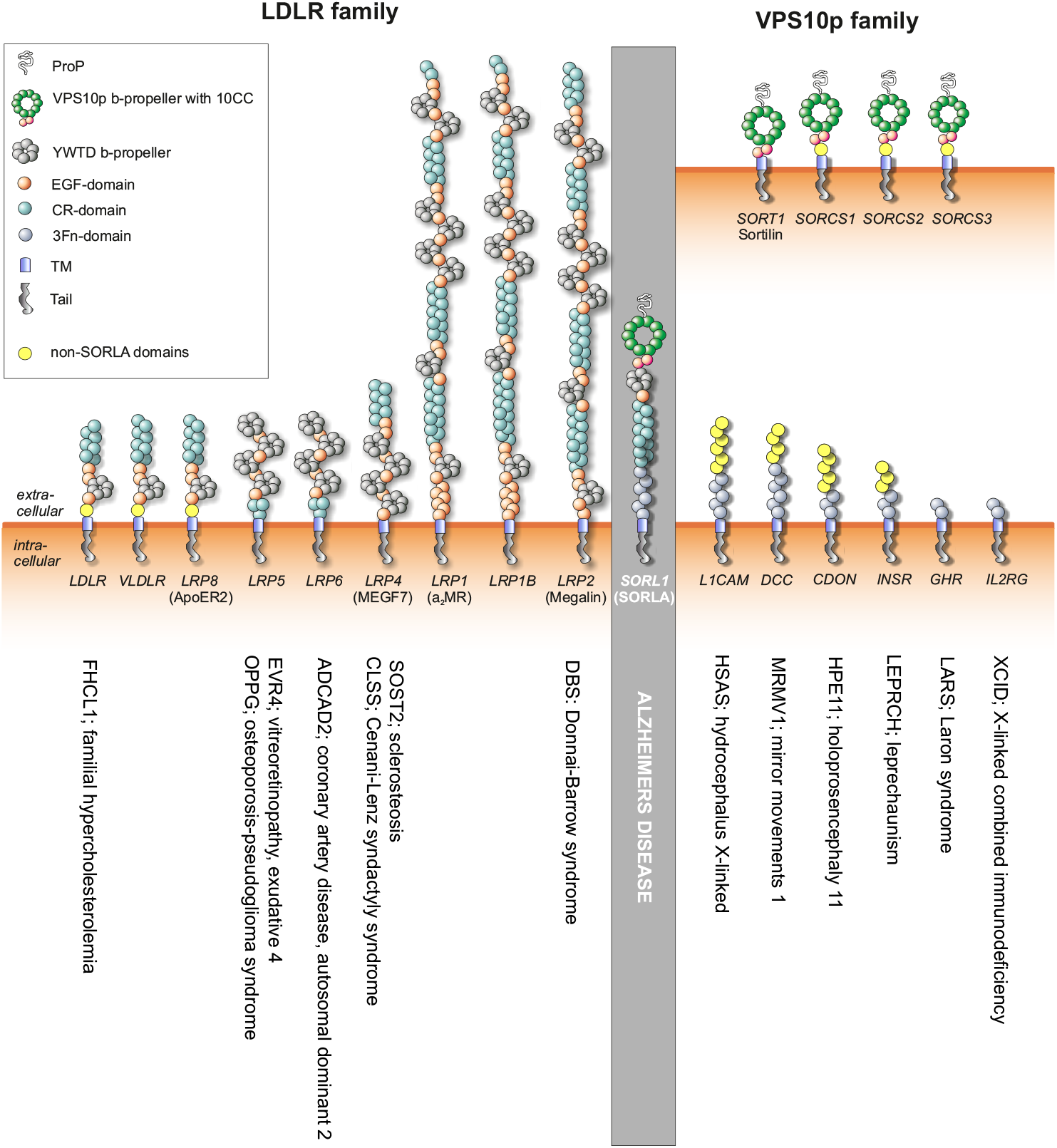
The SORLA modular receptor and homologous proteins. Schematic representation of the structural elements of SORLA and members of the mammalian LDLR- and VPS10p receptor families. Clustered copies of 3Fn-domains close to the membrane is present in a large number of unrelated proteins with diverse function (only a small subset included), thus not enabling assignment to any class of unique proteins like the other two receptor families. Some of the diseases the homologous proteins can cause when hit by pathogenic variants are listed below individual proteins.

Guided by its modular structure, we first performed a sequence alignment of SORLA’s many clustered domains or repeated sequences within the larger domains. We then rely on mutations in other monogenic diseases whose protein domains are homologous to those of SORLA to identify domain positions predicted to be pathogenic sites. Finally, we interrogated a set of *SORL1* variants previously identified in the ADES/ADSP exome sequencing study of unrelated AD cases and cognitively healthy controls ^12^, and tested to what extent variants in SORLA that reside at the disease-associated domain positions occurred in these individuals.

## METHODS

### SORLA DOMAIN SEQUENCE ALIGNMENTS

Manual alignment of the SORLA sequence was performed under the listed considerations:

#### VPS10p

Alignment of the 10 internally repeated sequences is based on β-blade boundaries and identified β-strand sequences as defined by the solved crystal structure in its uncomplexed form ^27^. There is only a limited conservation across the 10 sequences, so alignment is primarily done focusing on positions containing amino acid residues with hydrophobic side chains.

#### YWTD

Alignment of the 6 internally repeated sequences is based on β-blade boundaries of domains from the homologous LDLR domain ^28^ and guided by its β-strand sequences and conservation of a number of hydrophobic residues. Gaps in the alignment are preferentially assigned to positions in loop regions between β-strands. The sequences of YWTD-domains from SORLA and 14 other domains from proteins from the LDLR family are presented in **Supplemental Information 2d**.

#### EGF

The single eight-cysteines SORLA domain of the EGF-type is aligned either with 15 EGF-domain sequences from the LDLR family (selected from domains located C-terminal to YWTD-domains) or with 8 EGF-domain sequences from the integrins (**Supplemental Information 3d**).

#### CR

Alignment of the 11 individual CR-domain sequences from SORLA is done according to domain boundaries as defined by the genomic exon sequences. These domains sequences were also aligned with an additional 32 sequences from proteins containing pathogenic variants (**Supplemental Information 4d**)

#### 3Fn

Alignment of the 6 individual 3Fn-domain sequences is done based on a secondary structure prediction to identify the suggested position of 42 β-strands together with the presence of 4 highly conserved residues at domain positions 25 (W, strand B), 41 (Y, strand C), 77 (L, EF-loop), and 83 (Y, strand F) and allowing loops to accommodate most gaps. The SORLA domains were aligned with 3Fn-sequences from proteins containing disease-mutations in their 3Fn-domains, allowing the alignment to take into account also the partial conservation at positions 6, 7, 11, 13, 72, 74 and 94 (**Supplemental Information 5d**).

#### TM/CD

This part of the human SORLA sequence was aligned with 14 SORLA sequences from mammalian and less related species.

### SORLA-SPECIFIC DISEASE-MUTATION DOMAIN-MAPPING

The domain-mapping of disease-mutations (DMDM) approach displays an aggregate view of human pathogenic mutations by its position in a protein domain ^29^. This tool requires the ability to accurately assign the correct position of an observed variant within a domain sequence, and thus requires highly accurate alignments as here performed for the SORLA domains.

The VPS10- and EGF-domain sequences as well as the transmembrane and tail sequences were not included in the analysis: the VPS10p family is under-investigated, and there are no disease-associated variants firmly established for the other four family members (sortilin, SorCS1, SorCS2, SorCS3) (**Fig. 2**). However, the VPS10p-domain is the only domain in SORLA for which the structure has been determined ^27^, such that *SORL1* variants with unknown significance in the VPS10p-domain may be assessed based on the crystal structure to determine their impact on conformation folding/stability/energy. The EGF-domain in SORLA is longer than the typical ~40 amino acid EGF-domain, and thus alignments to other domains is not accurate and a disease mapping approach could therefore not be applied for this SORLA domain. The transmembrane and tail sequences are considered SORLA specific and has no direct comparable sequences in other proteins.

In contrast, we were able to apply the disease-mutation domain-mapping tool to SORLA domains having strong sequence consensus with homologous domains in other proteins such as the YWTD-CR-, and 3Fn-domains (comprising almost 2/3 of the entire receptor). To limit variants to a manageable number, we included sequences from **human** proteins in Uniprot.org, and manually curated a comprehensive overview of disease-associated variants using information provided by “*Natural variants*” involved in diseases (i.e., listed under “*Pathology & Biotech*” in sequences. **YWTD:** The sequence corresponding to the YWTD-domain is termed “LDL-receptor class B” by Uniprot and annotated as PS51120 (LDLRB) in PROSITE. In this database 14 of the 67 proteins with this domain type are human, of which 6 proteins included a total of 102 unique disease-associated variants that could be mapped (in red) onto the YWTD-domain sequence (**Supplemental Information 2d,e**). **CR**: The CR-domains are annotated as “LDL-receptor class A” in Uniprot and by PS50068 (LDLRA) in PROSITE. In the Uniprot database, 44 of the 168 proteins with CR-domains are human, of which 8 proteins contained at least one disease-associated variant within their CR-domain(s). This enabled us to map (in red) 63 different variants to the CR-domain sequences (**Supplemental Information 3d,e**). **3Fn**: These domains are reported by Uniprot as “Fibronectin type-III” and correspond to PS50853 (FN3) in PROSITE that list as many as 804 known Eukaryotic proteins of which 194 are human. In 35 of these, we identified naturally occurring disease-associated variants enabling us to map 222 unique disease-associated variants (in red) across the 3Fn-domain sequence (**Supplemental Information 5d,e**). Due to limited sequence conservation and variable number of amino acids between positions 50-71 of many 3Fn-domain sequences, variants that mapped to this part of 3Fn-domains were not included in the DMDM analysis.

A few additional variants that are not yet included in the Uniprot database at the time of the analysis were added for pathogenic variants identified from literature.

Finally, the number of disease-variants identified were plotted for each position within these three domain sequences, allowing the unambiguous identification of domain positions most likely to contain pathogenic variants when mutated in SORLA (**Supplemental Figures S2d, S4d, and S5e**). From the domain sequence alignments that established residues important for domain folding/stability (based on requirement of sequence conservation) in combination with the DMDM analysis informing on the prevalence of disease-associated mutations at given domain positions, we next generated a list/filter for the entire SORLA protein highlighting positions that we believe is of ‘*high priority*’ or ‘*moderate priority*’ risk for developing AD. This filter can be applied for interrogations of larger case/control dataset (Holstege *et al*., in preparation) or in a simpler manner by predicting whether single variants correspond to a dangerous position (**Supplemental Information 7**).

## RESULTS

Figure 1 represents a model that summarizes the domain affiliation of all the 2,214 amino acids of full-length human SORLA, which are explained in detail below and in the Supplemental Information 1-6. This diagram will form the basis for presentation of information for all identified *SORL1* variants online (www.alzforum.org/mutations).

### The VPS10p-and 10CC-domains (residues 1-753)

#### Propetide and signal peptide (residues 1-81)

After mRNA translation, the SORLA protein includes a Signalpeptide (SP: 1-28) and a Propeptide (ProP: 29-81) that precedes the VPS10p-domain of SORLA. These peptides are removed from the mature protein by a signal peptidase upon entry to the ER and by Furin in late Golgi/TGN compartments, respectively ^30^.

#### VPS10p (residues 82-617)

The VPS10p-domain itself is folded into a ten-bladed β-propeller structure with each of the ten blades composed of four antiparallel β-strands arranged around a central conical tunnel. Two loop regions are known to be important for binding activities of the VPS10p-domain extending the sequences of the sixths and sevenths blade ^27^.

The four repeated β-strands in each blade served as guide-sequence for the alignment of the ten blade sequences (**Fig. 3**). The domain structure depends on hydrophobic residues at several positions in the different β-strands (positions 5, 6, 19, 20, 21, 39, 40, and 41) and based on their contribution to domain stabilization (**Supplemental Information 1**), we predict that substitution of these residues to non-hydrophobic residues likely leads to a moderately increased risk for AD, whereas we predict that conservative substitutions at these positions are less harmful.

**Figure 3.**
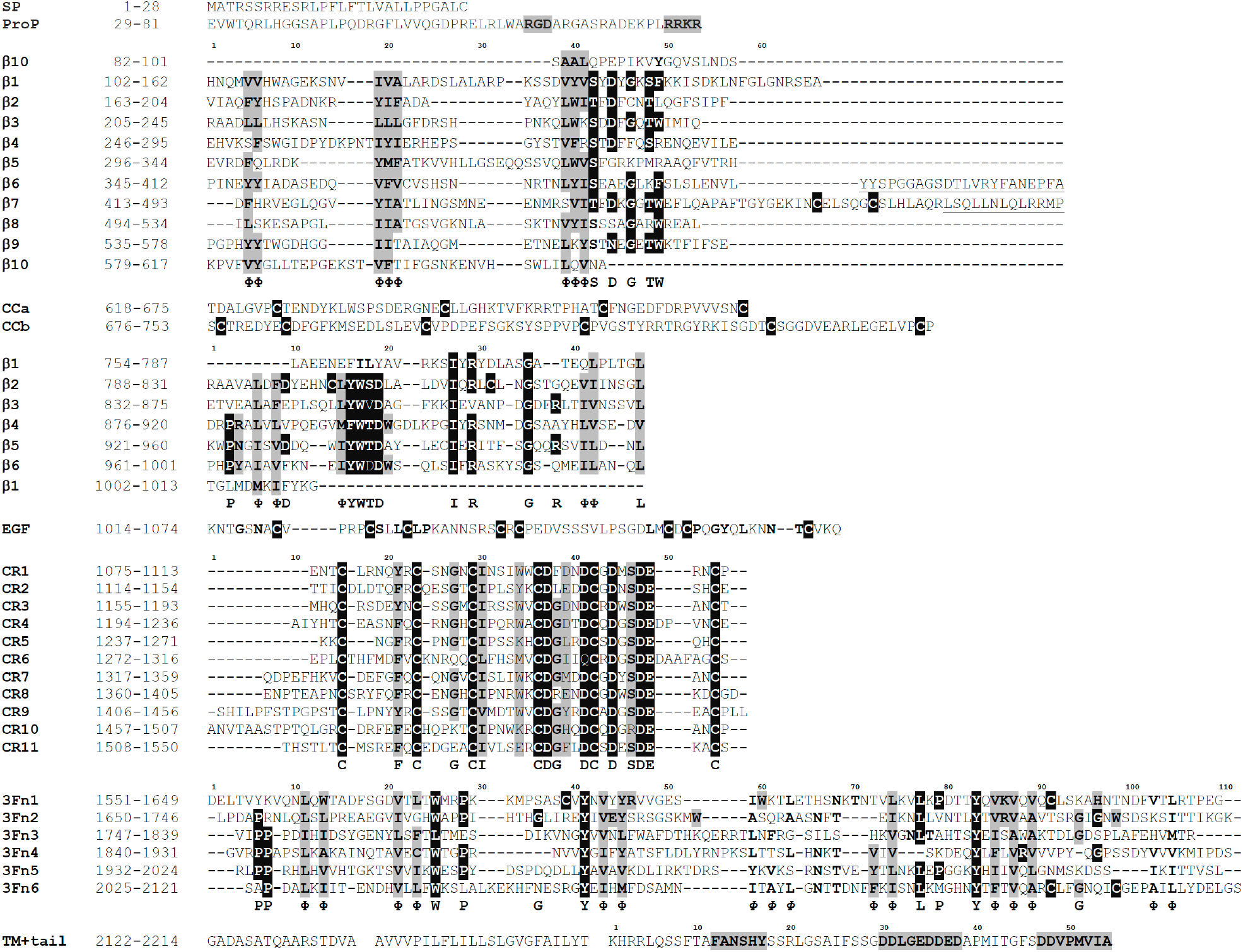
SORLA sequence alignments. Alignments of the different SORLA domains are presented in more details with indications of β-strand secondary structure in the respective Supplemental Information sections. Amino acid positions that are conserved and/or identified to frequently contain disease-causing mutations in proteins containing homologous domains and thus likely to provide a *high* risk of disease when mutated is shown in white letters on a black background. Positions suggested to provide a *moderate* increased risk for development of AD on a grey background.

#### VPS10p Asp-box and loops

Several residues in each blade are (partly) conserved. An Asp-box motif is completely or partially present in all β-blade sequences (**Supplemental Figure S1**): the exact function of the motif is not clarified, but based on speculations how these residues contribute to folding stability ^31^, we suggest that mutations that affect this motif might confer a high risk for AD. The sequence folding into the β-propeller domain includes only few Cys residues, such that an introduction of an additional Cys may not be pathogenic if in the domain core. However, a disulfide in the loop 2 that connects the seventh and eight blade, suggest that a mutation to a cysteine in either of the loop regions may be harmful. We previously identified variant p.Y391C in the sequence of loop 1, which was observed in 12 unrelated AD cases (mean age at onset 67.4 years) and in zero controls in the ADES/ADSP dataset ^12^, which supports our expectations that such variants are pathogenic.

The overall low sequence conservation between the ten blades complicates pathogenicity prediction. Therefore, we suggest to inspect the VPS10p-domain crystal structure for further insight for a given amino acid (RCSB PDB 3WSZ) ^27^. Second, the conservation of the residue in SORLA from other species can often inform whether a residue was conserved during evolution, but also whether the introduced ‘new’ residue by genetic mutation of human SORLA is tolerated in other SORLA proteins, which would suggest that pathogenicity is less likely. An alignment of 40 selected SORLA sequences is provided to identify positions that has not been conserved during evolution (**Supplemental Information 9**).

#### Cys residues in the 10CC domain (residues 618-753)

The 10CC region, which directly follows the β-propeller domain, is split into two shorter 10CCa and 10CCb subdomains, which stabilize the β-propeller fold of the VPS10p-domain. The domain has ten highly conserved Cys residues, and we predict that loss of any of these Cys residues, or a gain of an additional Cys in the 10CC sequence, leads to SORLA protein misfolding and an increased risk of AD. Therefore, such variants are among the *high* prioritized variants (**Supplemental Information 7**), and the ADES/ADSP dataset finds two such variants (p.C716W and p.Y722C) in 3 cases (mean age at onset 63.7 years) and absent from controls.

#### YWTD-repeated β-propeller (residues 754-1013)

Immediately following the VPS10p- and 10CC-domains, SORLA contains a region spanning 260 amino acids containing five incomplete copies of a characteristic YWTD-tetrapeptide (**Fig. 3**). YWTD-repeat regions generally fold into a compact 6-bladed β-propeller, each blade containing four antiparallel β-strands organized around a central pseudo symmetrical axis, forming an internal tunnel ^28,32^. The six repeated sequences that each form one blade, served as guide for the alignment of the six blade sequences (**Fig. 3; Supplemental Figure S2**).

#### YWTD positions 16, 17, 18, 19

We identified conserved positions in the YWTD motif itself (positions 16-19), and at positions 27 (Ile), 29 (Arg), and 35 (Gly), and where the side chains from these residues assist to stabilize domain folding, and we speculate that genetic variants that leads to changes of these residue, will likely associate with a highly increased AD-risk. Indeed, we found that position 19 (identifying 7 different disease variants of which six variants were substitution of an Asp) is the position most frequently affected by disease-mutations in the homologous YWTD-domains from LDLR family members (**Supplemental Information 2f**). In aggregate, we observed variants at positions 16-19 in the SORLA protein for a total of 8 AD cases (mean age at onset 59.0 years) and none for controls ^12^. The variants at YWTD-positions 16-19 in *SORL1* may thus be risk-increasing or even causative for AD, and should be regarded as high priority positions when variants are observed in a patient-carrier.

#### YWTD positions 6, 8, 15, 41, 42, and 47

Further, the fold of the six-bladed propeller depends on hydrophobic core interactions, similar to the VPS10p-domain, such that we predict that substitutions of hydrophobic residues at positions 6, 8, 15, 41, 42, and 47 into non-hydrophobic residues will provide a moderately increased risk of AD.

#### YWTD position 38

YWTD-repeated β-propellers are found in all core members of the LDLR family (**Fig. 2**) ^32^, and mutations in these proteins that lead to monogenic disease indicated that substitution of certain positions in the SORLA domain might increase the risk of AD. For example, when Arg residues at position 38 in the YWTD-domain of LDLR, LRP4 or LRP5 are substituted by other residues, this leads to autosomal dominant inherited forms of diseases (**Supplemental Information 2e**). Since the Arg at position 38 is not conserved across blades (proteins), prediction of pathogenicity based on sequence alignments alone is impossible. SORLA includes two blades with an Arg residue at position 38 (R866 in β3 and R953 in β5). The ADES-ADSP dataset includes three unrelated AD cases, with age at onset ranging between 46 and 58 years, that carry p.R953H; this variant was not observed in controls ^12^. This supports (but given the low numbers, does not prove) that rare *SORL1* variants at YWTD position 38 are pathogenic.

### EGF-domain (residues 1014-1074)

The SORLA protein contains only one copy of a 61 amino acid EGF-domain such that internal sequence-alignments are not available to guide residue pathogenicity prediction **(Fig. 3)**. Since eight Cys residues in this domain are all expected to participate in disulfide bonding, we predict that substitution of/with Cys residues (both Cys removal or introduction) will likely lead to a highly increased AD-risk (**Supplemental Figure S3**). The EGF-domains from the LDLR family members are only ~40 amino acid long, making it impossible to identify disease-associated residue substitutions in SORLA based on disease-associated substitutions in homologous EGF-domains from these proteins (**Supplemental Information 3d**). Deletion of the entire SORLA-EGF-domain, as identified in a family with AD ^33^, is likely to impair receptor activity. However, it is likely that, apart for substitutions involving cysteines, pathogenicity prediction of mutations in the EGF-domain will rely on in-silico pathogenicity prediction models in combination with functional studies as described elsewhere ^34^.

### CR-cluster (residues 1075-1550)

SORLA contains eleven CR-domains, each containing ~40 amino acids with several strictly conserved residues, which, when substituted by other residues are likely to profoundly increase the risk of AD. This applies to (i) 6 Cys residues at positions 15, 23, 29, 36, 42, 55, which form three invariable disulfide bridges ^35,36^ (**Fig. 3)**. (ii) 4 conserved acidic residues (at positions 37, 41, 47, 48) involved in the coordination of a Calcium ion (Ca^2+^) for each domain (**Fig. 3)**. (iii) A pair of conserved Asp and Ser residues at positions 44 and 46, that forms what is known as an Asx-turn structure. Furthermore, the CR-domains in SORLA also contain a pair of Gly residues (positions 27 and 38) that is conserved in eight of the eleven CR-domains (**Fig. 3**), together with a pair of conserved hydrophobic residues at positions 21 (Phe) and 30 (Ile) in the more N-terminal part of the sequence (**Supplemental Figure S4**). But as our domain mapping approach to our surprise did not find these positions enriched for disease-mutations we suggest that variants that target these CR-domain positions in SORLA lead to a moderately increased risk for AD.

#### Odd Numbered Cysteines (ONC), domain positions 15, 23, 29, 36, 42, 55 and random Cys introductions

Numerous disease-associated variants involved replacement or introduction of Cys in LDLR family members, incl. LDLR, LRP4, and LRP5, leading to an ‘*odd number of cysteines*’ (ONC). This was also observed for CR-domain-containing proteins outside the LDLR family, e.g. the transmembrane proteinase TMPRSS6 and the COP9 protein involved in the complement cascade. Together, these observations suggest that introduction of a Cys, or removal of one of the six Cys is a disease-causing event across CR-domain bearing proteins, and it is likely also true for SORLA (**Supplemental Information 4e**). Indeed, in a Swedish family a variant leading to p.R1303C in the sixth CR-domain was identified to segregate with AD in a small pedigree ^33^, and in a Saudi Arabian family, a variant leading to p.R1084C in the first CR-domain segregated with AD ^37^. Both variants resulted in CR-domains with 7 Cys residues. Also, a 59-year-old AD patient who carried a variant leading to p.C1192Y was reported ^38^, this variant results in the third CR-domain having 5 Cys residues. Furthermore, variants leading to p.C1344R was reported to associate with a possible family history in a Finnish family ^33^, as well as the p.C1453S and p.C1249S variants were reported for AD patients only ^9^. Also, ONC variants in *SORL1* identified in aggregate for the ADES-ADSP dataset were observed predominantly in AD cases (n=36; mean age at onset 67.8 years) compared to controls (n=8) ^12^. Hence, ONC substitutions in that dataset associate with a >6-fold increased risk of AD (OR = 6.31 95% CI: 2.45 - 16.24, p=5.1E-6; Fisher Exact test). Finally, we would like to highlight the similarity between ONC variants in *SORL1* associated with AD and variants in *NOTCH3* causal of Cerebral Autosomal Dominant Arteriopathy with Subcortical Infarcts and Leukoencephalopathy (CADASIL), where it has been established how stereotypic causal variants also result in an odd-number of cysteines in EGF-domains of NOTCH3 carrying 32-34 copies of this domain type ^39^.

#### Calcium Cage (CaCa), domain positions 37, 41, 47, and 48

In proteins with CR-domains, residues at positions 37 and 41 and 47 are all Asp (in SORLA, there is a single exception; Gln^1301^ at pos 41 in CR6) and positions 48 are all Glu (**Supplemental Information 4d**). The side chains of residues at these positions coordinate Ca^2+^ establishing an octahedral ‘*Calcium cage*’ (CaCa) (**Supplemental Figure S4**) which is critical for domain folding ^40–42^. As a consequence, substitutions of these variants may be strongly associated with disease. In LDLR, substitutions of CaCa residues lead to Familial hypercholesterolemia (FH) ^43,44^. We also identified disease-causing variants for CaCa positions in other proteins, e.g. LRP2, LRP5 and TMPRSS3 and TMPRSS6. Of interest, we also note two disease-associated substitutions of Asp with Glu at position 47 (**Supplemental Information 4e**), which is generally considered a conservative – and often non-pathogenic – substitution. However, in CR-domains there is not enough space in the Calcium cage to accommodate the larger Glu side chain at position 47 ^41^. Each SORLA protein includes 44 CaCa positions that can be mutated (4 possible positions in 11 domains), such that compared to other protein-features, disruption of the CaCa feature is relatively common. Carriers of CaCa disrupting genetic variants in SORL1 were previously reported only for AD patients: p.D1545E of CR11 (position 47; CADD score 15.9) ^45^, p.D1182N (position 41, CR3), and p.D1267E (position 47, CR5) ^9^ and p.D1389V (position 37, CR8) ^45–47^. Notably, a p.D1545V (position 47, CR11) variant is likely a dominant negative variant and was shown to segregate with AD in an Icelandic family, providing the first evidence for *SORL1* as an autosomal dominant Alzheimer gene ^24^. Furthermore, the ADES-ADSP dataset includes 11 calcium cages-affecting genetic variants, exclusively in AD cases (OR = INF): p.D1108N, p.D1219G, p.D1261G, p.D1267N, p.D1345N, p.D1389V, p.D1502G, p.D1535N, p.D1545N, p.D1545G, p.D1545E. These occurred in 13 unrelated AD cases with a relatively early age at onset (median 60 years, ranging from 47-73 years). Together, CaCa-related variants are associated with a highly increased risk of AD, and possibly causal for AD.

#### Asx-turn, domain positions 44, 46

The side chain of the conserved Asp at position 44 forms a structure known as an “*Asx-turn*” making a hydrogen bond with the backbone amides of two residues: one residue upstream at position 43, and a conserved Ser amino acid downstream at position 46 ^48^. Based on the sequence conservation and a number of disease-associated variants for other proteins, variants in *SORL1* that locate to these two conserved positions should also be considered as likely to increase risk of AD (**Supplemental Information 4f**). Indeed, SORL1 variants p.D1105H and p.D1146N are identified in 2 AD cases (mean age at onset 56.2 years) in the ADES/ADSP data and absent from the control cohort; while more evidence is necessary, these findings suggest that these variants are pathogenic.

We predict that only functionally important Asp residues in the CR-domains are pathogenic: for example, variant p.D1309N affects an Asp at domain position 50 which does not appear to have a functional consequence; this variant was not identified in AD patients in the ADES/ADSP dataset, although it was observed in 2 controls.

### 3Fn-cassette (residues 1551-2121)

3Fn-domains are typically composed of a sequence with 90-100 residues, arranged in seven β-strands (named A, B, C, C’, E, F, and G) forming two anti-parallel β-sheets (strands: A-B-E and strands: C-C’-F-G, respectively) (**Fig. 3; Supplemental Figure S5**). It is remarkable that despite high similarity in tertiary structure, sequence identity across 3Fn-domains is conspicuously low, typically less than 20% between domains ^49^, which complicates alignment of 3Fn-domain sequences.

However, the presence of a few highly conserved amino acids enables unambiguous identification of strands B, C, and F as described below (and explained in more detail in **Supplemental information 5**). Strand B is characterized by a Trp (position 25) preceded by two hydrophobic residues at positions 21 and 23; strand C contains a Tyr (position 41) followed by two hydrophobic residues at positions 43 and 45, with the latter position very often occupied by an additional Tyr; strand F begins with a Tyr (position 83) followed by three additional hydrophobic residues at positions 85, 87, and 89, with the latter position often being Ala. As the hydrophobic residues alternate within a β-strand secondary structure, their side chains point towards the same side of their respective strand, such that they form a large hydrophobic domain-core – sometimes described as ‘*the glue*’ between the two β-blades of the sandwich fold (**Supplemental Figure S5**). In contrast to the VPS10p-domain and the YWTD/EGF- and CR-domains that are representative of two distinct receptor families, a similar clear affiliation for 3Fn-domain containing proteins is not possible (**Fig. 2**). However, more than 2,100 domains are listed in PFAM as being 3Fn-domains ^50^, thus allowing us to perform a comprehensive disease-mutation domain-mapping approach also for this domain, relying on a large number of different proteins associated with a variety of different diseases (**Supplemental Information 5e**).

#### 3Fn-domain positions 25, 41, and 83

Based on the sequence conservation alone, we predict that any substitution of amino acids at the three key positions (25, 41, and 83) are *highly* likely to associate with increased risk for disease. Our disease-mutation domain-mapping analysis indicated that variants affecting positions 25 and 83 are among the most frequently mutated residues, with respectively 7 and 6 different disease mutations identified (**Supplemental Information 5f**). The proteins with disease-mutations at these positions are Usherin (USH2A), L1CAM, Fibronectin (FN1), Anosmin (ANOS1), MYBPC3 and the Growth-hormone receptor (for the Trp at position 25) and L1CAM, Insulin receptor (INSR), Fibronectin, Growth-hormone receptor and Tie2 (TEK) (for Tyr replacement mutations at position 83), in strong support of our prediction, which suggests that variants at these two positions are often pathogenic. It has previously been reported that the *SORL1* variant p.Y1816C (corresponding to position 83 in the third 3Fn-domain of SORLA) was found in AD patients but not controls ^9^, and also the ADES/ADSP dataset confirmed that this mutation was exclusively identified in 6 unrelated cases of AD (median age at onset 60.2 years) and not in controls ^12^. This is a high number taking into account that these are all non-related individuals, in strong support that this variant is risk increasing and possibly causal for AD. Indeed, for three of these cases a clear family history for AD and the *SORL1* p.Y1816C can be established (Jensen *et al*., in preparation).

#### 3Fn-domain positions 21, 23, 43, 45, 85, 87, and 89

The hydrophobic side chains of amino acids at positions 21, 23, 43, 45, 85, 87, and 89 all contribute to the ‘hydrophobic glue’ of the folded 3Fn-domain sandwich, and we therefore suggest that substitutions of these residues in SORLA to residues that are not hydrophobic will moderately increase risk of AD (**Fig. 3**).

Notably, conservative substitutions at these positions are less likely to increase AD risk, as exemplified by the common variant p.V2097I (position 87, the sixth 3Fn-domain of SORLA) which occurs in 33 controls and 48 AD cases of the ADES/ADSP dataset (OR=1.48; P= 8.2E-02), suggesting that such variants do not significantly associate with increased risk of AD ^12^.

#### 3Fn-domain positions 6, 7, 11, 13

The four remaining β-strands do not contain highly conserved residues and thus much more difficult to unambiguously identify without solved structures (**Supplemental Figure S5**). However, pairs of alternating hydrophobic residues may contribute to a hydrophobic core: positions 11 and 13 in strand A are often two hydrophobic amino acids approximately 5-8 amino acids upstream of strand B. Similar as for the hydrophobic residues at the other strands, we predict that subsitutions at positions 11 and 13 will have a moderate effect on disease-risk. On the other hand, depending on the subdomain, one or two Pro residues at positions 6 and 7 of the 3Fn-domain sequence are partly conserved. Their location indicate a functional role to form the turn of the protein backbone prior to the following β-strand and start of the 3Fn-domain (**Supplemental Information 5d**), such that we predict that substitution at these positions are likely to associate with high risk of disease. We find 3 cases of AD with substitutions at these Pro-positions, mean age at onset 76 years.

The other three strands (C’, E, and G) have very little sequence conservation complicating the identification of their locations based on amino acid sequence analysis. Therefore, we could not trustfully align amino acids at 3Fn-domain positions 50-71 and 99-111 with homologous proteins for the identification of disease-associated substitutions (**Supplemental Figure S5**, **Supplemental Information 5f**).

#### 3Fn-domain positions 77, 79, 80 (and 83): the tyrosine corner

The loop connecting strands E and F (EF-loop) that crosses from one sheet to the other, contains a Leu (position 77) located six positions upstream of the conserved Tyr at position 83 in the beginning of strand F. In many 3Fn-domains, including two of the SORLA domains, this loop also contains a conserved Pro (position 79) frequently accompanied by a Gly (position 80) (**Supplemental Figure S5**). This structural motif is known as “*the tyrosine corner*”, which contributes strongly to the stability of 3Fn-domains: the side chain of Leu-77 packs next to the Tyr-83 ring ^51^. Moreover, the phenol group of the Tyr-83 engages in hydrongen bonding with the backbone of the residue five residues upstream (ie. position 78), naming the 3Fn-domain in SORLA as the Δ5 subtype of tyrosine corners ^52^. While all other loops in the 3Fn-domain can elongate without significant loss of conformational stability, the length of the EF-loop is critical to maintain a stable domain fold ^53^. As a result of the high sequence conservation and functional importance we suggest that variants corresponding to positions 77 and 79 should be considered as highly risk increasing when affecting Leu or Pro, respectively (**Fig. 3**).

#### 3Fn-domain loop positions 27, 28, 36, 94 and 96

The loops at the top of the 3Fn-domain contain several conserved positions: the BC-loop often begins with one or two Pro at positions 27 and/or 28 and a Gly at position 36, and the FG-loop preferentially contains a Gly at position 94 – often in combination with a Gly at position 96 (**Supplemental Information 5d**). Accordingly, we assign a high priority for these risk positions (**Fig. 3**).

We identified seven disease-associated variants at position 96 of the 3Fn-domain according to our domain-mapping analysis (**Supplemental Information 5f**). According to the 3Fn-domain consensus sequence, the Gly at position 96 was not conserved, whereas the Gly at position 94 was partly conserved (**Supplemental Figure 5d**). Nevertheless, in five of the seven identified disease variants for position 96, a Gly was substituted. Furthermore, four of these variants were observed when the Gly two residues upstream (at position 94) in the sequence was also present, and we speculate that co-occurrence of the two Gly residues could have functional relevance relating to their localization in the FG-loop region, preferring to accommodate residues with small side chains. In SORLA, only the second 3Fn-domain contains this double Gly at positions 94 and 96 (amino acids 1730 and 1732) (**Fig. 3**). Interestingly, variant p.G1732A corresponding to position 96 is reported to segregate with AD in a Swedish family ^33^, in support of this variant being pathogenic.

#### 3Fn-domain ‘hotspot’ position 88

For position 88 of the 3Fn-domain, eight variants were identified in different proteins and associated with different diseases, suggesting that position 88 may be a mutational hotspot (**Supplemental Information 5f**). Notably, each of these disease-causing variants correspond to the substitution of an Arg, suggesting that an Arg at this position serves an indispensable function. For SORLA, only R1910 at position 88 in the fourth 3Fn-domain contains an Arg (**Fig. 3**), and thus far no variants have been reported for this position in SORLA. However, we predict that any mutation affecting position 1910 will highly increase risk of AD, and we are curiously awaiting to see if our prediction holds true.

### Transmembrane and cytoplasmic domains (residues 2122-2214)

Immediately following the 3Fn-domains is a 16 amino acid long stalk region (residues 2122-2137) suggested to lift the ectodomain a short distance from the plasma membrane, enabling TACE-dependent cleavage, leading to ectodomain shedding when mature SORLA is at the cell surface ^54,55^. After the stalk region, SORLA contains a 23 amino acid single-pass transmembrane (TM) domain (residues 2138-2160), presumably alpha-helical, and a cytoplasmic domain (CD; often referred to as the ‘*tail*’) including 54 amino acids (residues 2161-2214) (**Fig. 3**). These regions of SORLA are not amenable for sequence alignment with other proteins, but for completion of this compendium and to guide understanding the impact of potential disease-causing variants in *SORL1* that target this region, a detailed description of these regions and their contribution to SORLA activity is included in **Supplemental Information 6**.

#### Tail motif residues 2172-2177, 2190-2198, and 2207-2214

Binding of cytosolic proteins that assist in the intracellular trafficking of SORLA has been shown to mainly rely on the three motives: ‘FANSHY’ (residues 2172-2177), ‘acidic’ (residue 2190-2198), and ‘gga’ (2207-2214). Variants that lead to changes of these amino acids may provide a moderate risk-increase for AD.

## DISCUSSION

Here, we show that variants in the *SORL1* gene that affect specific SORLA protein functions may severaly increase AD risk, or some may even be causative for AD, often with a very early age at onset. One of these variants has been observed in an Icelandic family with an autosomal dominant inheritance pattern of AD ^24^, supporting that *SORL1* can be considered the fourth autosomal dominant Alzheimer gene ^10^, next to *APP, PSEN1*, and *PSEN2*. These genes biologically converge on inducing endosomal traffic jams ^56,57^ and all induce amyloid secretion from neurons. However, apart from very few variants, pedigrees of *SORL1* variant carriers are either not available, or uninformative for analysis of penetrance, complicating determining the variant pathogenicity.

Here we addressed this issue by relying on structural information and known pathogenic variants in proteins containing homologous domains as those present in SORLA. Using this diseasemapping approach we identified a number of positions in the receptor sequence that are likely to contain variants that increase the risk of developing AD, or that might even be causal for AD. The presentation of potentially protein impairing variants within the full protein sequence can be considered a compendium that can be exploited by a range of investigators. Most notably, we identified CaCa substitutions and ONC changes in the CR-domains, that was either entirely restricted to AD patients [OR = INF] or associated with a highly increased risk [OR =6.3] of developing AD, respectively. But also other findings were in strong support of our approach: for example that variants that lead to substituions of the YWTD-motif occurred also exclusively in patients with AD.

This comprehensive compendium, describing the many aspects of the SORLA protein function and the genetic variation in the *SORL1* gene, will inform a wide array of researchers. For structural biologists and biochemists, interested on the effect of variation on protein fold and SORLA function, the compendium provides deeper insight to SORLA structure-function relationships. For clinical geneticists, interested in the potential effect of genetic variants on disease risk, this compendium guides a focused, hypothesis-driven annotation of newly identified genetic variants for potential pathogenicity. We envision that this compendium will support investments on clinical counseling of AD patients beyond the pathogenic mutations in *APP, PSEN1* and *PSEN2*.

Among the many millions of AD patients in the US alone, a number that is exponentially rising with time ^58^, identifying those who are potentially affected by *SORL1* -associated AD is essential. Not only for the development of appropriate diagnostics and counseling procedures, but also for the development of personalized therapeutics for *SORL1* variant carriers.

## Acknowledgement

H.H. and O.M.A. are part of the EU Joint Programme-Neurodegenerative Disease Research (JPND) Working Group SORLA-FIX under the 2019 ‘‘Personalized Medicine’’ call (JPND2019-466-197, ZonMW 733051110, Danish Innovation Foundation and the Velux Foundation Denmark). H.H., is a recipient of ABOARD, a public-private partnership receiving funding from ZonMW (#73305095007) and Health~Holland, Topsector Life Sciences & Health (PPP-allowance; #LSHM20106). H.H. was supported by the Hans und Ilse Breuer Stiftung (2020) and the HorstingStuit Foundation (2018). O.M.A is supported by Novo Nordisk Foundation (#NNF20OC0064162), the Alzheimer’s Association (ADSF-21-831378-C), and the Danish Alzheimer’s Research Foundation (recipient of the 2022 Basic Research Science Award).

We would like to thank Gregory Petsko and Scott Small for their support and critical reading of our manuscript, and we are immensely grateful to Kathy Zahs and Elizabeth Wu for their collaborations to generate the online platform available at alzforum.org.

