## Supplemental Information for "Relying on the relationship with known disease-causing variants in homologous proteins to predict pathogenicity of *SORL1* variants in Alzheimer’s disease"

#### TABLE OF CONTENT

1 THE VPS10P-DOMAIN (RESIDUES 1-753) 3
1.A SEQUENCE DETAILS 3
1.B SUPPLEMENTAL FIGURE S1. VPS10P-DOMAIN 7
1.C *SORL1* VARIANTS IN VPS10P-DOMAIN 8
2 THE YWTD-REPEATED Β-PROPELLER (RESIDUES 754-1013) 9
2.A SEQUENCE DETAILS 9
2.B SUPPLEMENTAL FIGURE S2. YWTD-DOMAIN 13
2.C *SORL1* VARIANTS IN YWTD-DOMAIN 14
2.D YWTD-DOMAIN ALIGNMENT 17
2.E IDENTITY OF MAPPED YWTD VARIANTS: DISEASE PROTEINS AND DISEASE VARIANTS 20
2.F YWTD DISEASE VARIANTS LISTED ACCORDING TO DOMAIN POSITIONS 22
3 THE EGF-DOMAIN (RESIDUES 1014-1074) 24
3.A SEQUENCE DETAILS 24
3.B SUPPLEMENTAL FIGURE S3. EGF-DOMAIN 27
3.C *SORL1* VARIANTS IN EGF-DOMAIN 28
3.D EGF-DOMAIN ALIGNMENT 29
4 THE CR-CLUSTER (RESIDUES 1075-1550) 31
4.A SEQUENCE DETAILS 31
4.B SUPPLEMENTAL FIGURE S4. CR-DOMAINS 35
4.C *SORL1* VARIANTS IN CR-DOMAINS 36
4.D CR-DOMAIN ALIGNMENT 40
4.E IDENTITY OF MAPPED CR VARIANTS: DISEASE PROTEINS AND DISEASE VARIANTS 41
4.F CR DISEASE VARIANTS LISTED ACCORDING TO DOMAIN POSITIONS 43
5 THE 3FN-CASSETTE (RESIDUES 1551-2121) 45
5.A SEQUENCE DETAILS 45
5.B SUPPLEMENTAL FIGURE S5. 3FN-DOMAINS 50
5.C *SORL1* VARIANTS IN 3FN-DOMAINS 51
5.D 3FN-DOMAIN ALIGNMENT 53
5.E IDENTITY OF MAPPED 3FN VARIANTS: DISEASE PROTEINS AND DISEASE VARIANTS 59
5.F 3FN DISEASE VARIANTS LISTED ACCORDING TO DOMAIN POSITIONS 62
6 TRANSMEMBRANE AND CYTOPLASMIC DOMAINS (RESIDUES 2122-2214) 66
6.A SEQUENCE DETAILS 66
6.B SUPPLEMENTAL FIGURE S6. TRANSMEMBRANE AND CYTOPLASMIC DOMAINS 70
6.C *SORL1* VARIANTS IN TRANSMEMBRANE AND TAIL-DOMAINS 71
7 SUMMARY OF DOMAIN POSITION PRIORITIZATION FOR AD RISK 73
8 SORL1 SEQUENCES FOR ALIGNMENT: SPECIES CONSERVATION 78
9 SORL1 SEQUENCE ALIGNMENT 79
10 PHYLOGENETIC TREE FOR SORLA 80
10.A SUPPLEMENTAL FIGURE S7. PHYLOGENETIC TREE 81
11 SUPPLEMENTAL METHODS 82
12 PUTTING THE PIECES TOGETHER - THE CONFORMATIONAL SPACE 83
12.A SUPPLEMENTAL FIGURE S8. THE SORLA CONFORMATIONAL SPACE 88
13 SUPPLEMENTAL REFERENCES 89

### 1 The VPS10p-domain (residues 1-753)

#### 1.a Sequence details

*Signal peptide (residues 1-28)*

The most N-terminal part encoded by the *SORL1* gene is a signal peptide (SP) (residues 1-28) that directs the polypeptide into the endoplasmatic reticulum (ER), and is lost by signal peptidase cleavage upon translocation into the ER, similar to other transmembrane proteins. Accordingly, functional SORLA protein in cells does not contain this initial part of the polypeptide.

*Propeptide (residues 29-81)*

Residues 29-81 of SORLA is a propeptide (ProP), that can be removed from the remaining part of the receptor by enzymatic cleavage by the prohormone convertase furin that is active in secretory vesicles in the late Golgi/TGN [1]. The $^{78}$RRKR$^{81}$ tetrapeptide serves as a recognition site for furin binding and cleavage between residues 81-82 [2]. The ProP also contains a $^{63}$RGD$^{65}$ tripeptide motif, which serves as an interaction site for adhesive proteins, including integrins in other proteins. This suggests that SORLA proteins that escape cleavage by furin still include a ProP and may have a unique function in cell adhesion and integrin binding. The presence of ProP is also suggested to block binding of small ligands to the VPS10p-domain (see next section), which is speculated to prevent binding of certain ligands to the VPS10p-domain in the ER of cells where receptor and ligand are co-expressed [2].

*10-bladed β-propeller (residues 82-617)*

After the ProP follows the VPS10p-domain: with its 536 residues, this domain is one of the largest protein domains known. There is only modest sequence conservation between domains across the five members of the VPS10p family (SORLA, sortilin, SorCS1, SorCS2, SorCS3) [3]. The crystal structure of the VPS10p-domain of sortilin was solved in 2009 [4], and in 2015 for the SORLA domain [5], making it the only full SORLA-domain for which the three-dimensional conformation is currently determined. Both sortilin and SORLA VPS10p-domain structures are organized in a ten-bladed β-propeller, each blade composed of four antiparallel β-strands (A-D) arranged around a central conical tunnel. C-terminal to both VPS10p-domains is a small domain, corresponding to 136 amino acids of SORLA including ten conserved cysteine residues with a stringent spacing, known as the 10CC region. This domain (residues 618-753 in SORLA) interacts extensively with the β-propeller (see below) (**Supplemental Figure S1a**). The overall dimensions of the β-propeller resemble a 94 Å x 72 Å x 56 Å ellipsoid, and for SORLA the central cavity narrows toward the bottom face with a maximum width of

~25 Å [5]. The β-propeller is further characterized by a defined and almost flat bottom and top face, demarcated by the loops between strands A-B and C-D, and strands B-C and D-A, respectively (**Supplemental Figure S1b,c**). Interestingly, the solved domain structures of both sortilin and SORLA were determined in the presence of complexed ligands, which indicated that small (peptide) ligands can bind inside the tunnel.

Alignment of the ten sequences that define each blade, β1-β10 (residues 82-617) (**Supplemental Figure S1c**), revealed several conserved and functional motifs. The presence of two stretches of hydrophobic amino acids in the inner strands (A: positions 5, 6, and B: positions 19, 20, 21) of each blade is critical for domain stability. Similarly, 2-3 hydrophobic residues in strand C (positions 39, 40, 41) establish hydrophobic interactions among the blades that enable the formation of the large central tunnel. Additionally, they partake in the generation of a hydrophobic surface that allows the interaction of small lipophilic ligands with the propeller cavity, as demonstrated for the binding of the Amyloid-β peptide to SORLA [5].

*Asp-boxes (part of β-propeller)*

The motif with consensus sequence S(or T)-X-D(or N)-X-G-X-T(or S)-W(or F/Y) (spanning positions 42-49 of each blade) is known as the Asp-box [6], which folds as a conserved β-hairpin [7]. The SORLA VPS10p-domain contains Asp-boxes in the loops located (at the bottom of the domain) between the third (C) and fourth (D) strands of the blades (**Supplemental Figure S1b,c**). The exact function of this motif has not been clarified. However, the Asp-boxes of the SORLA domain are unlikely to be directly involved in ligand recognition, since only the top face of propellers appears to be used for this purpose [8]. Rather, the conserved amino acids of the Asp-box motif likely stabilize the propeller, forming blade-to-blade interactions, contacts with preceding loops, or with the nearby 10CC-domains.

*L1-L2 Loops (part of the β-propeller)*

The two sequences connecting β6 with β7 and β7 with β8 are longer than the sequences connecting other blades. Part of these two longer protrusions (residues Y391-F411 and A457-P493, respectively) include the two "*loop structures*" termed L1 (residues Y391-A412) and L2 (residues L481-P493), respectively [5] (**Supplemental Figure S1c**; loop residues underligned). L2 is located close to the upper entrance of the tunnel, and seems to push bound ligand against the tunnel wall at neutral pH. The L1 segment occupies a more central position, blocking part of the pore. Interestingly, L1 is not present in VPS10p-domains in other proteins, which suggests that the VPS10p-domain in SORLA has a unique ligand binding profile. Indeed, both L1 and L2 appear to be essential for peptide binding since removal of either of these two protrusions completely abolishes binding activity [5]. Both loops are flexible and undergo conformational changes at different pH, assuming a more stable conformation at pH 6.5 than at pH 4.5. The disordering of the loops at low pH was suggested as important for ligand release at acidic pH, providing a molecular mechanism for dissociating ligands from this SORLA domain in lysosomes, when SORLA encounters the low pH in this organelle.

*10CC region (residues 618-753)*

The VPS10p β-propeller fold is stabilized by the neighboring 10CC region, which is split into two shorter domains named 10CCa (residues 618-675) and 10CCb (residues 676-753). While rather similar to each other, these domains show neither sequence nor structural resemblance to other known domains. The ten conserved cysteines form five intrachain disulfide bridges with connectivity between cysteines as $\text{Cys}^{I}$-$\text{Cys}^{III}$ and $\text{Cys}^{II}$-$\text{Cys}^{IV}$ (in 10CCa), and $\text{Cys}^{V}$-$\text{Cys}^{IX}$, $\text{Cys}^{VI}$-$\text{Cys}^{VII}$, and $\text{Cys}^{VIII}$-$\text{Cys}^{X}$ (in 10CCb) [5,9]. Structurally, the two domains wrap around the bottom face of the propeller, and form strong contacts with the propeller through numerous hydrogen and ionic bonds. Attempts to express either the propeller or the 10CC regions alone were not successful, which suggests that these interactions between propeller and the 10CC region ensure a compact structure and provide stability to the entire unit. However, the 10CC region is mobile and undergoes a pH-dependent conformational change as evidenced for both sortilin and SORLA [4,5]. For SORLA the 10CCb-domain exhibits the largest rearrangement, with a “lever-like” motion when the pH increases from acidic conditions with the 10CCb-domain is tightly associated with the propeller, to more neutral conditions with the 10CCb-domain forming almost no contacts. Keeping in mind that full-length SORLA is a modular multi-domain protein, such a movement may affect the overall receptor conformation and is likely relevant for ligand binding activity and possibly receptor dimerization as it traffics between cellular compartments with different pH conditions.

Andersen
Supplemental Figure S1 - VPS10p-domain

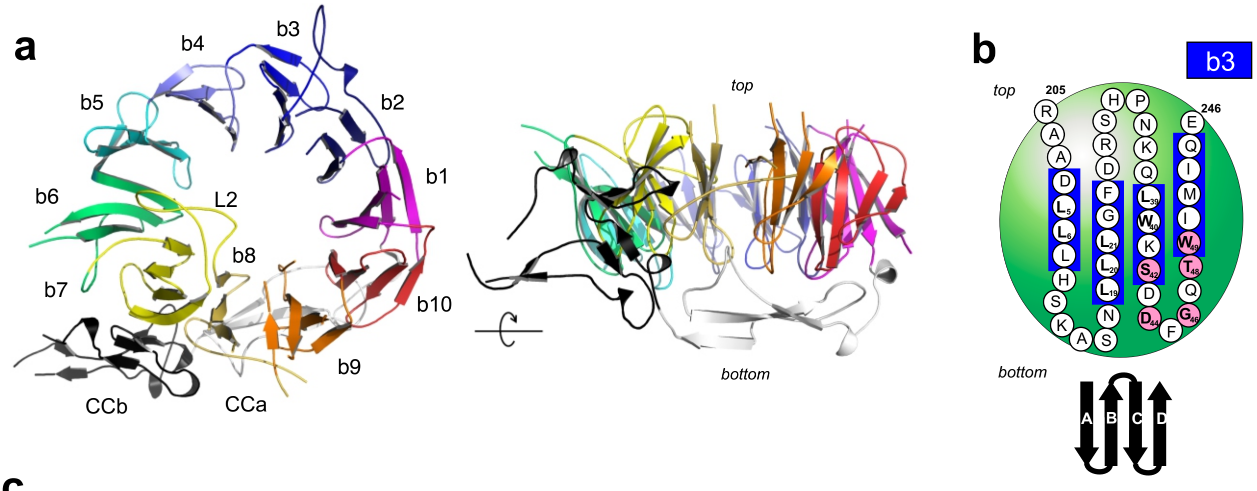

**c**

```
SP      1-28     MATRSSRRESRLPFLFTLVALLPPGALC
ProP   29-81     EVWTQRLHGGSAPLPQDRGFLVVQGDPRELRLWARGDARGASRADEKPLRRKR

                   A         bottom     B         top          C       bottom  D    top

                 1        10        20        30        40        50        60
β10   82-101     -------------------------------------SAALQPEPIKVYGQVSLNDS-----------------------------------
β1   102-162     HNQMVVHWAGEKSNV---IVALARDSLALARP--KSSDVYVSYDYGKSFKKISDKLNFGLGNRSEA--------------------------
β2   163-204     VIAQFYHSPADNKR----YIFADA----------YAQYLWITFDFCNTLQGFSIPF------------------------------------
β3   205-245     RAADLLLHSKASN-----LLLGFDRSH-------PNKQLWKSDDFGQTWIMIQ---------------------------------------
β4   246-295     EHVKSFSWGIDPYDKPNTIYIERHEPS-------GYSTVFRSTDFFQSRENQEVILE-----------------------------------
β5   296-344     EVRDFQLRDK--------YMFATKVVHLLGSEQQSSVQLWVSFGRKPMRAAQFVTRH-----------------------------------
β6   345-412     PINEYYIADASEDQ----VFVCVSHSN-------NRTNLYISEAEGLKFSLSLENVL-------------YYSPGGAGSDTLVRYFANEPFA (L1)
β7   413-493     ---DFHRVEGLQGV----YIATLINGSMNE----ENMRSVITFDKGGTWEFLQAPAFTGYGEKINCELSQGCSLHLAQRLSQLLNLQLRRMP (L2)
β8   494-534     ---ILSKESAPGL-----IIATGSVGKNLA----SKTNVYISSSAGARWREAL---------------------------------------
β9   535-578     PGPHYYTWGDHGG-----IITAIAQGM-------ETNELKYSTNEGETWKTFIFSE------------------------------------
β10  579-617     KPVFVYGLLTEPGEKST-VFTIFGSNKENVH---SWLILQVNA-------------------------------------------------
                     ΦΦ            ΦΦΦ                 ΦΦΦS D G TW

                        Cys^I-Cys^III                     Cys^II-Cys^IV
CCa  618-675     TDALGVPCTENDYKLWSPSDERGNECLLGHKTVFKRRTPHATCFNGEDFDRPVVVSNC
CCb  676-753     SCTREDYECDFGFKMSEDLSLEVCVPDPEFSGKSYSPPVPCPVGSTYRRTRGYRKISGDTCSGGDVEARLEGELVPCP
                        Cys^VI-Cys^VII          Cys^V-Cys^IX                     Cys^VIII-Cys^X
```

#### 1.b Supplemental Figure S1. VPS10p-domain

**a**. The VPS10p-domain folds into a 10-bladed β-propeller (PDB: 3WSX [5]) presented from the top (left) and from the side (right), with the 10CC-domains located at the bottom of the folded propeller. Each blade is colored according to the code from ref [5].

**b**. Amino acids 205-246 corresponding to β3 are presented by their one-letter-code. The blue rectangular background indicates the four A-D anti-parallel strands. Asp-box residues are presented on a pink background towards the bottom of the folded domain in the C-D loop. Sheet topology is indicated below.

**c.** Residues 1-753 in SORLA comprise a signal peptide (SP; residues 1-28), a pro-peptide (ProP; residues 29-81), 10 repeats forming the VPS10p-domain (residues 82-617), and two parts with conserved cysteines (CCa/b; residues 618-753). Numbering of the sequence follows Uniprot entry Q92673 from as originally described by ref [10]. The 10 repeats are aligned based on the identified four β-strands within each repeat (grey; A-D), with positions of conserved hydrophobic amino acids in strands A-C as well as amino acids of the Asp-box motifs (**S**X**D**X**G**X**TW**) shown in bold. Horizontal lines indicate the locations of the five disulfides that bridge the 10 cysteines in CCa and CCb.

#### 1.c *SORL1* variants in VPS10p-domain

The VPS10p-domain is found only in the 5 proteins of the VPS10p-receptor family, with too little pathological information such that there are no orthologous proteins suitable for evaluation of disease-associated variants. In many cases pathogenicity may be inferred from interrogation of the available crystal structure [5]. This has been done for the p.G511R variant (predicted as "*likely pathogenic*"), which was previously reported to segregate with AD across 2 generations [11]. While the residue is located outside the L1/L2 regions, it maps to a position that fixes one end of the L2 loop that is directly involved in Aβ binding. This suggests that conversion of the small glycine to a large amino acid (i.e. like arginine) could cause a severe disturbance in the conformation (or stability) of L2 with possibly the loss of Aβ-binding ability [5]. In line with this reasoning, functional studies suggested that this mutation impairs binding of Aβ to the VPS10p-domain, leading to decreased lysosomal delivery of Aβ and a consequential increase of secreted Aβ [12]. Taken together, this variant may, at least in part, explain the observed AD in the family [11].

Despite the very small (A528T) or absence (E270K) of a variant effect on AD-risk observed in GWAS studies, functional assays using cells transfected with these variants indicated an impaired ability of mutant SORLA to decrease APP processing [13]. Inspection of the VPS10p crystal structure and our sequence alignment (**Fig. 3, Supplemental Figure S1c**) is in agreement with neither of these variants being located at very dangerous positions.

### 2 The YWTD-repeated β-propeller (residues 754-1013)

#### 2.a Sequence details

Immediately following the VPS10p- and 10CC-domains, SORLA contains a region spanning 260 amino acids containing five incomplete copies of a characteristic YWTD-tetrapeptide (**Supplemental Figure S2c, strand B**). In general, a YWTD-repeat region folds into a compact 6-bladed β-propeller, and each blade contains four antiparallel β-strands that are organized around a central pseudo symmetrical axis, forming an internal tunnel. The dimensions of this β-propeller type mimic a 68 Å x 58 Å x 40 Å ellipsoid. Accordingly, this domain is significantly smaller than the 10-bladed VPS10p β-propeller, and there are no reports describing ligands being able to bind inside these narrow tunnels, that often are so tight they appear closed. YWTD-repeated β-propellers are found in all core members of the LDLR family (**Supplemental Figure S2a**) as well as in the physiologically unrelated proteins Nidogen, Osteonidogen, and the precursor of EGF [14].

Crystal structures of homologous domains from LDLR [15], LRP4 [16], LRP6 [17-20], ApoER2 [21] and Nidogen [22] have all been solved, showing how only 5 Å separates the N- and C-terminal residues of the domain in space although they are separated by 260 amino acids in the primary structure (**Supplemental Figure S2**).

*Alignment – unique residues*

As in homologous YWTD-propellers, the YWTD-motif is absent from the β1 blade where the tetrapeptide is represented by the $^{760}$FILY$^{763}$ sequence (**Supplemental Figure S2c**). For blades β2-β6, the YWTD-motifs are located in the second (B) strand (**Supplemental Figure S2c**). The conserved aromatic residues at the Tyr and Trp positions (16 and 17 of each blade) are embedded inside the domain and contribute to an apparent hydrophobic core. The aspartate residues at position 19 participate in extensive hydrogen bonding with residues in strand A and C of the same blade and strand A of the adjacent blade, thereby maintaining the structural integrity of the β-propeller fold [15] (**Supplemental Figure S2b**). Hydrophobic residues within strand A (positions 6 and 8), B (position 15), and D (positions 41 and 42) are present in blades β2-β6 and add further to a hydrophobic core (**Supplemental Figure S2b,c**). A conserved isoleucine (position 27) also contributes to stabilization of the hydrophobic contacts. An arginine is frequently located at position 29 in strand C, where it also functions in domain stabilization through interactions with the Tyr of the YWTD-motif. Three of the six β-propeller blades include a conserved Pro residue at position 3, which supports the loop-structure between blades. Furthermore, a Gly (position 35) and Leu (position 47) represent conserved positions in most YWTD-domain sequences including that of SORLA (**Fig. 3, Supplemental Figure S2**). Together, the conserved residues in the YWTD-domain of SORLA point towards an essential role in maintaining the rigidity of the propeller.

*A SBiN-type YWTD-domain in SORLA*

To understand how YWTD-domains interact with binding partners, several studies have analyzed the structures of ligand-bounded YWTD-domains [22]. The ligand often contains an Asn-Ile pair (the NXI-pair, with the N and I residues separated by a variable residue) that binds a motif called the *Shutter Binding NXI* (SBiN)-motif, found in several YWTD-domain sequences [16 23]. The Asn side chain of the ligand NXI-pair interacts with a Trp (pos 20), a Phe (pos 4) and an Asn (pos 4) of the β-propeller, whereas the Ile of the ligand NXI-pair engages with an additional paired Trp (pos 20) - Arg (pos 4) of the YWTD-domain [22]. In aggregate, the residues at these five positions all locate to positions 4 or 20 of the β-blade sequence alignment and make up the SBiN-motif [16,23]. The SORLA YWTD-propeller contains an SBiN-motif composed of residues W895 (β4, pos 20), N924 (β5, pos 4), Y964 (β6, pos 4), and the R879-W978 (β4 pos 4-β6 pos 6) pair (**Supplemental Figure S2c – residues highlighted, black background**). All five residues are brought together at the top of the folded domain in line with their position in the sequence either before strand A or at the end of strand B.

The SORLA sequence contains five *internal* NXI-pairs, which suggests that other SORLA domains may also serve as ligand to the β-propeller to form intramolecular (*intrinsic*) contacts. The five internal NXI-pairs are located in blades β1 and β4 of the VPS10p-domain ($^{115}$NVI$^{117}$ and $^{262}$NTI$^{264)}$, in the first CR-domain ($^{1089}$NCI$^{1091}$ and $^{1092}$NSI$^{1094}$), and in the sixth 3Fn-domain ($^{2105}$NQI$^{2107}$). As the two NXI sequences within the VPS10p-domain are situated at the bottom of the large β-propeller (in agreement with their position in the primary sequence between strands A and B), it is unlikely they would be in contact with the YWTD-domain SBiN motif. The recently determined model of the full SORLA protein ectodomain by AlphaFold [24] shows no indication that internal NXI-pairs bind to the YWTD-domain.

*Takes two to tangle*

Our phylogenetic analysis indicates that in all known SORLA receptors the single YWTD-propeller coexists with the preceding VPS10p β-propeller, suggesting these two domains form a single rigid unit (**Supplemental Figure S7**). This is in agreement with the crystal structure of the extracellular domain of LRP6, which contains four YWTD-propellers that form pairs of two rigid structural blocks, with a short intervening hinge that restrains their relative orientation [18]. This pairing is observed in several proteins with multiple YWTD-domains (i.e. LRP4 and LRP6): it enables interactions with large ligands including co-receptors in multimeric complexes, or large soluble ligands requiring two adjacent β-propellers for efficient binding [18] [16]. It is tempting to speculate that both the VPS10p and YWTD β-propellers of SORLA exclusively bind ligands to their top faces. The SBiN residues in the YWTD domain locate to the top face (**Supplemental Figure S2c**), while an EGF-domain, located to the C-terminal end of the β-propeller, forms intimate domain-domain interactions with the bottom face, making it unavailable for ligand-interactions [25]. Similarly, the bottom face of the VPS10p-domain is occupied by the 10CC-domains, such that this side is also unavailable for ligand-interactions.

Andersen
Supplemental Figure S2 - YWTD-domain

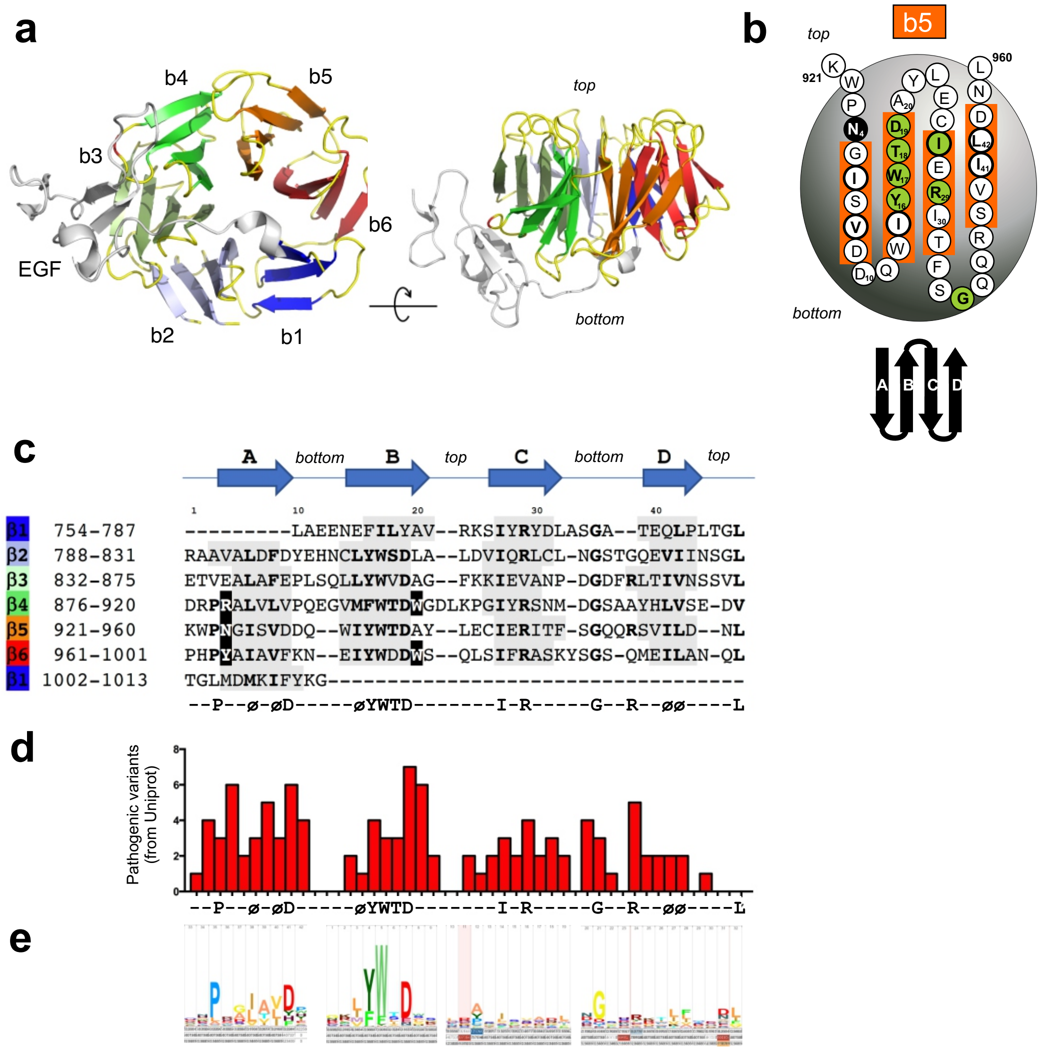

#### 2.b Supplemental Figure S2. YWTD-domain

**a**. The YWTD-domain of LDLR folds into a 6-bladed β-propeller (PDB: 1IJQ [15]), likely reflecting a similar structure for the YWTD-domain of SORLA. The propeller is shown from the top (left) and from the side (right). The EGF-domain of LDLR is shown in light grey at the bottom of the propeller.

**b**. Schematics of the fifth β-sheet (β5) of the β-propeller (gray circle), depicting residues 921-960 by their one-letter-code with indication of the four A-D anti-parallel strands in orange. The YWTD-motif in strand B together with some of the few additionally conserved residues of each repeat are presented with green circles and bold letters. Blade topology is shown below.

**c**. Alignment of the SORLA sequence spanning residues 754-1013 and corresponding to an YWTD-domain was done based on the YWTD-motif and the presence of the predicted four β-strands of each repeat (grey background). Lower case letters or capital letters indicate positions occupied by similar or identical residues in each repeat, respectively. Residues $R^{879}$, $W^{905}$, $N^{924}$, $Y^{964}$, $W^{979}$ forming the SBiN-motif are shown in white letters on a black background.

**d.** Histograms showing the number of pathogenic variants that occur for each YWTD-domain position as listed in **Supplemental Information 2f**.

**e.** Logo representation of the domain sequence conservation: the larger the letter the higher the conservation across YWTD-domain sequences.

#### 2.c *SORL1* variants in YWTD-domain

*YWTD-motif at positions 16-19:*

Not surprisingly, all four positions (16-19) of the tetrameric YWTD-sequence are potentially deleterious of protein/domain function (**Supplemental Figure S2d**). **Tyr (pos 16),** was identified to be mutated in LDLR1, LRP2, and twice in LRP5 for different patient/diseases. Intriguingly, for each disease-variant the Tyr is replaced by a His: p.Y442H$^{LDLR}$ (identified in patients with FHCL1; [26]), p.Y2522H$^{LRP2}$ (considered causal of Donnai-Barrow syndrome (DBS); [27]), p.Y733H$^{LRP5}$ (identified in a patient with OPPG; [28]), and p.Y1168H$^{LRP5}$ (identified in woman with total retinal detachment and retinoschisis (EVR4); [29]) (**Supplemental Information 2e**). This suggests that introduction of a His is particularly damaging for position 16 of the propeller-domain. **Trp (pos 17)**, was frequently mutated in patients with FH (W577S$^{LDLR}$ [30], W577G$^{LDLR}$ [31], W577R$^{LDLR}$ [26], and functional characterization of mutant proteins with either Gly substitution [31] or Ser substitution [32] showed that substitutions completely abolish receptor membrane expression and LDL uptake. **Thr (pos 18)** in β-propellers of LRP5 (p.T390K$^{LRP5}$) associates with increased risk for osteoporosis-pseudoglioma syndrome (OPPG) [28] or is considered causal (p.T253I$^{LRP5}$) of Osteopetrosis, Autosomal Dominant 1 (OPTA1) in two related families on Fyn in Denmark [33], respectively. **Asp** (**pos 19),** mutations of this residue in LDLR are linked to familial hypercholesterolaemia (FH) (p.D492N$^{LDLR}$ [34], p.D579N$^{LDLR}$ [30,35,36] (for this variant less than 2% of receptor activity is reported) and p.D579Y$^{LDLR}$ [37]), while mutations in LRP5 or LRP4 are found in patients with OPPG (p.D434N$^{LRP5}$ [28]) or considered causal of Cenani-Lenz syndactyly syndrome (CLSS) (p.D529N$^{LRP4}$ [38], p.D1403H$^{LRP4}$ [39]), respectively (**Supplemental Information 2e**). Taken together, this suggests that variants at these positions in *SORL1* may be risk-increasing or even causative for AD, and should alert the clinical geneticist as variants with high priority when observed in a patient-carrier. In line with this p.D806N occurring at position 19 was observed so far only in AD patients and not controls [40].

*SBiN-motif at positions 4 and 20:*

In proteins that contain a YWTD-repeated β-propeller with SBiN-motifs, three of the six blades include residues of the ligand-binding SBiN motifs (i.e. β4-β6) at positions 4 and 20 (**Supplemental Figure S2d**) [23]. Variants that map to these positions in blades β4-β6 may be more pathogenic than if they map to β1-β3. This is supported by the pathogenicity of variants listed in Uniprot in LDLR, LRP4, and LRP5 that affect the SBiN-residues: p.N564S$^{LDLR}$ [41] and p.N564H$^{LDLR}$ [30,34,42-44] (position 4 of β5) in patients with FH. Functional analysis of p.N564H$^{LDLR}$ found 64-73% reduced uptake and degradation of LDL in fibroblasts from heterozogous and compound carriers [43,44]. Variant p.W1186S$^{LRP4}$ was identified in patient with Sclerosteosis (SOST2) [45] and show impaired Wnt-suppressing activity of the mutant receptor [46], and p.W478R$^{LRP5}$ (position 20 both in β4) in a family with OPPG where the variant segregate with affected subjects [47], respectively. Interestingly, *SORL1* variant p.N924S (conservation of Asn: 40/40 if **Supplemental Information 9**) affects the Asn residue in the SBiN-motif (position 4 in β5) has been reported in patients with AD [40], but there is no functional studies nor genetic statistical support to yet claim this variant to be pathogenic.

*Arg at position 29:*

In SORLA, 5 of 6 YWTD blades contain an Arg at position 29. Uniprot lists 4 pathogenic variants for this position, corresponding to substitution of an Arg: p.R570W$^{LRP5}$ (patients with OPPG [28,48]), p.R570Q$^{LRP5}$ (patients with EVR [28,49]), p.R1277H$^{LRP4}$ (patients with CMS17 [45]), and p.R473Q$^{LRP6}$ (segregate with disease in a family with metabolic syndrome; ADCAD2 [50]). This suggests that removal of Arg at position 29 may increase risk of AD. The preference for Arg at this position is not obvious from the logo-web consensus (**Supplemental Figure S2e**), but is noticeable in our alignments (**Supplemental Information 2d**).

*Asp at position 9:*

Position 9 is the top position for mutations in YWTD-propellers (**Supplemental Figure S2d**) with 6 disease-associated variants listed in Uniprot: p.D482H$^{LDLR}$ (in FHCL1 patients, listed as causal genetic variant [51,52]), p.D203N$^{LRP5}$ (in OPPG patients [28]), p.D381N$^{LRP5}$ (identified in a small family with familial EVR1 and functional analysis showed mutations lead to complete receptor inactivity [53]), p.D511A$^{LRP5}$ (identified in a small family with familial EVR4 [54]), p.D683N$^{LRP5}$ (identified in patients with OPPG [28]), and p.T852M$^{LRP5}$ (identified in a family with EVR4 and determined as pathogenic because of 95% reduction in LRP5 activity [55]). Notably, there is also preference of Asp at position 9 in the consensus sequence (**Supplemental Information 2d**), and the above listed mutations strongly suggest that substitutions that replace an Asp is very likely disease-associated. In the SORLA sequence only two of the six β-blades have an Asp at position 9; D794 (β2) and D929 (β5) (**Fig. 3**). The SORL1 variant p.D929Y has been identified in both a control and an AD case leaving it an open question still whether this is a dangerous position for the SORLA domain [40].

We would like to provide an example how to apply our species conservation tool, trying to decide whether the variant leading to p.V884M substitution is dangerous. Based on the identification of this variant in 4 AD cases and not yet in any controls [40], it could be speculated that this variant is pathogenic. But when using the alignment of multiple SORLA sequences from 40 different species (**Supplemental Information 9**), it is evident that there is no specific requirement for a valine at this position of SORLA as other hydrophobic residues as Leu and Ile are present at this position for other species, and several species even carry a Met at this position (e.g. SORLA from horse, salmon, pike, piranha, and zebrafish). This strongly suggests that the p.V884M is a benign mutation.

*Arg at position 38:*

Our DMDM analysis based on pathogenic variants listed in Uniprot identified 4 variants for position 38 that associate with four diseases: one in LDLR: p.R595W$^{LDLR}$ (in FHCL1 patients [34]), and three in LRP5: p.R494Q$^{LRP5}$ (in patients with OPPG [28,48]), p.R752G$^{LRP5}$ (in a family with EVR4 [49]), and p.R1188W$^{LRP5}$ (segregate with disease in large family pedigrees with PCLD4 [56]). Although no strong preference for an Arg in the SORLA sequence (**Fig. 3**), but a slight enrichment in the bigger sequence alignment for the β-blade at this position (**Supplemental Information 2d**), it is interesting that each of the four disease-variants involve the replacement of this positively charged amino acid. In SORLA two of the YWTD blades have an Arg at position 38: R866 (β3) and R953 (β5) (**Fig. 3**). We suggest that variants that affect these two amino acids may be deleterious of SORLA function and associated with AD. The ADES-ADSP dataset includes p.R953H in three cases with very early ages at onset ranging between 46 and 58 years, and not in controls [40].

#### 2.d YWTD-domain alignment

Mapping of naturally occurring variants found in human YWTD-domain containing proteins, and being listed as associated with pathology at [www.uniprot.org](http://www.uniprot.org). The alignment follows the SORLA alignment shown on top, and pathogenic variants are highlighted on a red background.

```
         1           10          20          30          40
                  LAEENEFILYAV--RKSIYRYDLASGA--TEQLPLTGL
        RAAVALDFDYEHNCLYWSDLA--LDVIQRLCL-NGSTGQEVIINSGL
        ETVEALAFEPLSQLLYWVDAG--FKKIEVANP-DGDFRLTIVNSSVL
        DRPRALVLVPQEGVMFWTDWGDLKPGIYRSNM-DGSAAYHLVSE-DV
        KWPNGISVDDQ--WIYWTDAY--LECIERITF-SGQQRSVILD--NL
        PHPYAIAVFKN--EIYWDDWS--QLSIFRASKYSGS-QMEILAN-QL
        TGLMDMKIFYKG-----------------------------------
          p  Φ Φd     ΦYWTD     I R    dG  r  ΦΦ    L
LDLR:
                 KAVGSIAYLFFTN--RHEVRKMTL-DRSEYTSLIPNL
        RNVVALDTEVASNRIYWSDLS--QRMICSTQLDRAHGVSSYDTVISRDI
        QAPDGLAVDWIHSNIYWTDSV--LGTVSVADT-KGVKRKTLFRENG
        SKPRAIVVDPVHGFMYWTDWG-TPAKIKKGGL-NGVDIYSLVTENI
        QWPNGITLDLLSGRLYWVDSK--LHSISSIDVNGGN-RKTILEDEKRL
        AHPFSLAVFED--KVFWTDII--NEAIFSANRLTGS-DVNLLAENL
        LSPEDMVLFHN
LRP2_6:
                  AISTENFLIFALSNSLRSLHLDPENHSPPFQTINVE
        RTVMSLDYDSVSDRIYFTQNLASGVGQISYATLSSGIHTPTVIASGI
        GTADGIAFDWITRRIYYSDYL--NQMINSMAE-DGSNRTVIARV
        PKPRAIVLDPCQGYLYWADWD-THAKIERATL-GGNFRVPIVNSSL
        VMPSGLTLDYEEDLLYWVDAS--LQRIERSTL-TGVDREVIVNAA
        VHAFGLTLYGQ--YIYWTDLY--TQRIYRANKYDGSGQIAMTTNLL
        SQPRGINTVVKNQKQQ
LRP5_1:
                PAPAAASPLLLFAN---RRDVRLVDAGGVKLESTIVVSGL
        EDAAAVDFQFSKGAVYWTDVS-EEAIKQTYLNQTGAAVQNVVISGL
        VSPDGLACDWVGKKLYWTDSE--TNRIEVANL-NGTSRKVLFWQDL
        DQPRAIALDPAHGYMYWTDWG-ETPRIERAGM-DGSTRKIIVDSDI
        YWPNGLTIDLEEQKLYWADAK--LSFIHRANL-DGSFRQKVVEGSL
        THPFALTLSGD--TLYWTDWQ--TRSIHACNKRTGGKRKEILSAL
        YSPMDIQVLSQERQPFFHTR
LRP5_2:
                  KAGAEEVLLLAR--RTDLRRISL-DTPDFTDIVLQVDDI
        RHAIAIDYDPLEGYVYWTDDE--VRAIRRAYL-DGSGAQTLVNTEI
        NDPDGIAVDWVARNLYWTDTG--TDRIEVTRL-NGTSRKILVSEDL
        DEPRAIALHPVMGLMYWTDWG-ENPKIECANL-DGQERRVLVNASL
        GWPNGLALDLQEGKLYWGDAK--TDKIEVINV-DGTKRRTLLED
        KLPHIFGFTLLGDFIYWTDWQ--RRSIERVHK-VKASRDVIID
        QLPDLMGLKAVNVAKVVGTNP
LRP5_3:
                   IVPEAFLVFTS--RAAIHRISL-ETNNNDVAIPLTGV
        KEASALDFDVSNNHIYWTDVS--LKTISRAFM-NGSSVEHVVEFGL
        DYPEGMAVDWMGKNLYWADTG--TNRIEVARL-DGQFRQVLVWRDL
        DNPRSLALDPTKGYIYWTEWG-GKPRIVRAFM-DGTNCMTLVDKV
        GRANDLTIDYADQRLYWTDLD--TNMIESSNM-LGQER-VVIAD
        DLPHPFGLTQYSDYIYWTDWN--LHSIERADKTSGRNR-TLIQGHL
        DFVMDILVFHSSRQDGLND
LRP5_4:
                   SPPTTFLLFSQ--KSAISRMIPDDQHSPDLILPLHGL
        RNVKAIDYDPLDKFIYWVDGR---QNIKRAKD-DGTQPFVLTSLSQGQNPD
        RQPHDLSIDIYSRTLFWTCEA--TNTINVHRL-SGEAMGVVLRGDR
        DKPRAIVVNAERGYLYFTNMQDRAAKIERAAL-DGTEREVLFTTGL
        IRPVALVVDNTLGKLFWVDAD--LKRIESCDL-SGANRLTLEDANI
        VQPLGLTILGK--HLYWIDRQ--QQMIERVEKTTGDKRTRIQGRVAHLTGI
```

HAVEEEVSLEEFSAHP

**LRP4_1:**

KALGPEPVLLFAN--RIDIRQVLP-HRSEYTLLLNNL
ENAIALDFHHRRELVFWSDVT--LDRILRANL-NGSNVEEVVSTGL
ESPGGLAVDWVHDKLYWTDSG--TSRIEVANL-DGAHRKVLLWQNL
EKPRAIALHPMEGTIYWTDWG-NTPRIEASSM-DGSGRRIIADTHL
FWPNGLTIDYAGRRMYWVDAK--HHVIERANL-DGSHRKAVISQGL
PHPFAITVFED--SLYWTDWH--TKSINSANKFTGKNQ-EIIRNKL
HFPMDIHTLHPQRQPAGKN

**LRP4_2:**

ISSHACAQSLDKFLLFAR--RMDIRRISF-DTEDLSDDVIPLADV
RSAVALDWDSRDDHVYWTDVS--TDTISRAKW-DGTGQEVVVDTSL
ESPAGLAIDWVTNKLYWTDAG--TDRIEVANT-DGSMRTVLIWENL
DRPRDIVVEPMGGYMYWTDWG-ASPKIERAGM-DASGRQVIISSNL
TWPNGLAIDYGSQRLYWADAG--MKTIEFAGL-DGSKRKVLIGSQL
PHPFGLTLYGE--RIYWTDWQ--TKSIQSADRLTGLDRETLQENLENLMDIHVFHRRRP
PVSTPCAMEN

**LRP4_3:**

PTGINLLSDGKTCSPGMNSFLIFAR--RIDIRMVSL-DIPYFADVVVPINITM
KNTIAIGVDPQEGKVYWSDST--LHRISRANL-DGSQHEDIITTGL
QTTDGLAVDAIGRKVYWTDTG--TNRIEVGNL-DGSMRKVLVWQNL
DSPRAIVLYHEMGFMYWTDWG-ENAKLERSGM-DGSDRAVLINNNL
GWPNGLTVDKASSQLLWADAH--TERIEAADL-NGANRHTLVSPV
QHPYGLTLLDS--YIYWTDWQ--TRSIHRADK--GTGSNVILVRS
NLPGLMDMQAVDRAQPLGF

**LRP4_4:**

DPSPETYLLFSS--RGSIRRISL-DTSDHTDVHVPVPEL
NNVISLDYDSVDGKVYYTDVF--LDVIRRADL-NGSNMETVIGRGL
KTTDGLAVDWVARNLYWTDTG--RNTIEASRL-DGSCRKVLINNSL
DEPRAIAVFPRKGYLFWTDWG-HIAKIERANL-DGSERKVLINTDL
GWPNGLTLDYDTRRIYWVDAH--LDRIESADL-NGKLRQVLVSHV
SHPFALTQQDR--WIYWTDWQ--TKSIQRVDKYSGRNKETVLANVEGL
MDIIVVSPQRQTGTNA

**LRP6_1:**

VLLRAAPLLLYAN--RRDLRLVDATNGKENATIVVGGL
EDAAAVDFVFSHGLIYWSDVS--EEAIKRTEFNKTESVQNVVVSGL
LSPDGLACDWLGEKLYWTDSE--TNRIEVSNL-DGSLRKVLFWQEL
DQPRAIALDPSSGFMYWTDWG-EVPKIERAGM-DGSSRFIIINSEI
YWPNGLTLDYEEQKLYWADAK--LNFIHKSNL-DGTNRQAVVKGSL
PHPFALTLFED--ILYWTDWS--THSILACNKYTGEGLREIHSDI
FSPMDIHAFSQQRQPNATNP

**LRP6_2:**

KDGATELLLLAR--RTDLRRISL-DTPDFTDIVLQLEDI
RHAIAIDYDPVEGYIYWTDDE--VRAIRRSFI-DGSGSQFVVTAQI
AHPDGIAVDWVARNLYWTDTG--TDRIEVTRL-NGTMRKILISEDL
EEPRAIVLDPMVGYMYWTDWG-EIPKIERAAL-DGSDRVVLVNTSL
GWPNGLALDYDEGKIYWGDAK--TDKIEVMNT-DGTGRRVLVED
KIPHIFGFTLLGDYVYWTDWQ--RRSIERVHK-RSAEREVIIDQLPDLMGLKATNVHRVIG
SNPMGAVLRSLLA

**LRP6_3:**

IVPEAFLLFSR--RADIRRISL-ETNNNNVAIPLTGV
KEASALDFDVTDNRIYWTDIS--LKTISRAFM-NGSALEHVVEFGL
DYPEGMAVDWLGKNLYWADTG--TNRIEVSKL-DGQHRQVLVWKDL
DSPRALALDPAEGFMYWTEWG-GKPKIDRAAM-DGSERTTLVPNV
GRANGLTIDYAKRRLYWTDLD--TNLIESSNM-LGLNR-EVIAD
DLPHPFGLTQYQDYIYWTDWS--RRSIERANKTSGQNR-TIIQGHL
DYVMDILVFHSSRQSGWNE

**LRP6_4:**

SAPTTFLLFSQ--KSAINRMVI-DEQQSPDIILPIHSL
RNVRAIDYDPLDKQLYWIDSR--QNMIRKAQE-DGSQGFTVVVSSVPSQNLE
IQPYDLSIDIYSRYIYWTCEA--TNVINVTRL-DGRSVGVVLKGEQ
DRPRAVVVNPEKGYMYFTNLQERSPKIERAAL-DGTEREVLFFSGL
SKPIALALDSRLGKLFWADSD--LRRIESSDL-SGANR-IVLEDSNI
LQPVGLTVFEN--WLYWIDKQ--QQMIEKIDM-TGREGRTKVQARI
AQLSDIHAVKELNLQEYRQHP

#### 2.e Identity of mapped YWTD variants: disease proteins and disease variants

Summary of included proteins containing naturally occurring variants associated with diseases in domains shared with SORLA.

| Gene | Protein | Uniprot | #dom | Associated Diseases/Syndromes (CODE) | Variants mapped onto big alignments (from Uniprot entries except where a specific ref is provided) | #variants | #positions | Citations for variants discussed in main text |
|---|---|---|---|---|---|---|---|---|
| LDLR | Low-density lipoprotein receptor | P01130 | 1x6 | Familial hypercholesterolemia (FHCL1) | FHCL1: A399D, L401V, F403L, T404P, R406W, E408K, L414R, D415G, R416W, I423T, V429M, A431T, L432V, D433H, T434K, **Y442H**, I451T, T454N, L479P, D482H, W483R, **D492N**, V523M, P526S, G549D, N564H/N564S, L568V, R574C/R574H, **W577G/W577S/W577R**, **D579N/D579Y**, I585T, G592E, **R595W**, D601H, P608S, R633C, V639D, P649L | 43 | 38 | **Y422H** identified in two patients with FHCL1 [26]<br>**D482H** as causal of FH [51,52]<br>**R595W** in FH pts [34]<br>**D492N** in FH pts [34]<br>**N564S** in Polish familial FH [41]<br>**N564H** in Brazilian, German, French and Danish familial FH [30,34,42-44]<br>**W577G** in FH pts, and functional test show no membrane expression and zeron LDL uptake, ie pathogenic [31]<br>**W577R** in FH pts [26]<br>**W577S** in FH pts [30] and functional test showed no LDLR maturation nor surface expression [32]<br>**D579N** has less than 2% receptor activity [30,35,36]<br>**D579Y** in FH pts [37] |
| LRP4 | Low-density lipoprotein receptor-related protein 4 | O75096 | 4x6 | Cenani-Lenz syndactyly syndrom (CLSS)<br>Sclerosteosis 2 (SOST2)<br>Myasthenic syndrome, congenital, 17 (CMS17) | CLSS: D449N, T461P, L473F, **D529N**, I450V [39], L953P [39],<br>Syndactyly: **D1403H** [57], Q1564K [57]<br>SOST2: R1170W, **W1186S**<br>CMS1: E1233K, **R1277H** | 12 | 12 | **W1168S** in SOST2 pts and with impaired Wnt-suppressing activity [45,46]<br>**R1277H** in patients [45] |
| LRP5 | Low-density lipoprotein receptor-related protein 5 | O75197 | 4x6 | Vitreoretinopathy, exudative 1 (EVR1)<br>Vitreoretinopathy, exudative 4 (EVR4)<br>Osteoporosis-pseudoglioma syndrome (OPPG)<br>High bone mass trait (HBM)<br>Endosteal hyperostosis, Worth type (WENHY)<br>Osteopetrosis, autosomal dominant 1 (OPTA1) | EVR1: **D381N**, R348W<br>EVR4: L145F, T173M, A422T, E441K, R444C, **D511A**, A522T, T535M, L540P, G550R, **R570Q**, **R752G**, T798A, R805W, **T852M**, N1182D, **Y1168H**<br>OPPG: **D203N**, T244M, R348W, R353Q, S356L, **T390K**, A400E, G404R, T409A, **D434N**, E460K, | 49 | 47 | D203N, T390K, D434N, G520V, Y733H identified in OPPG patients [28]<br>**W478R** in a family with OPPG and segregates with disease [47] |

| | | | | | | | | |
|---|---|---|---|---|---|---|---|---|
| | | | | Polycystic liver disease 4 with or without kidney cysts (PCLD4) | **W478R**, R494Q, W504C, **G520V**, N531I, **R570W**, D683N, **Y733H**, D1099Y, R1113C<br>**HBM**: R154M, M282V<br>**WENHY**: A214T/A214V, A242T<br>**OPTA1**: D111Y, G171R, A242T, **T253I**,<br>**PCLD4**: V454M, V684A, | | | **D381N** in a family with EVR. Functional test show complete mutant receptor inactivation [53]<br>**D511A** in a family with EVR [54]<br>**R570W** in pts with OPPG [28,48]<br>**R570Q** in pts with EVR [28,49]<br>**R752G** in a family with EVR [49]<br>**Y1168H** identified in patient with EVR [29]. Further evidence needed to establish if causal<br>**T852M** in a family with EVR and pathogenic based on 95% reduced mutant activity [55]<br>**T253I** most likely disease causing and segregate in two (related) families on Fyn/Denmark [33] |
| LRP6 | Low-density lipoprotein receptor-related protein 6 | O75581 | 4x6 | Coronary artery disease, autosomal dominant, 2 (ADCAD2) | **ADCAD2**: R360H, N433S, **R473Q** | 3 | 3 | **R473Q** segregate with disease in a family with metabolic syndrome [50] |
| LRP2 | Low-density lipoprotein receptor-related protein 2 | P98164 | 8x6 | Donnai-Barrow Syndrom (DBS) | **DBS**: **Y2522H** [27] | 1 | 1 | **Y2522H** causal of DBS [27] |
| | | | | | | **108** | 101 | |

#### 2.f YWTD disease variants listed according to domain positions

Disease-mutations domain-mapping analysis with identification of pathogenic variants in other proteins with YWTD-domains (as listed in **Supplemental Information 2e**). Here variants are mapped onto domain positions following alignment of internally repeated sequences in the SORLA domain sequences. The number of hits for each position depicted in the bar diagram of **Supplemental Figure 2d**.

| **YWTD-domain positions** | Number of hits | Identified variants | Domain conservation | Priority |
|---|---|---|---|---|
| 1 | 1 | p.R360H(LRP6) | | |
| 2 | 4 | p.E460K(LRP5), p.W504C(LRP5), p.R805W(LRP5), p.R1113C(LRP5) | | |
| 3; p | 3 | p.V429M(LDLR), p.P608S(LDLR), p.P649L(LDLR) | Loss of Pro | high |
| 4(sbin) | 6 | p.N564S(LDLR), p.N564H(LDLR), p.D111Y(LRP5), p.R154M(LRP5), p.M282V(LRP5), p.R154M(LRP5) | Loss of SBIN | moderate |
| 5 | 2 | p.A431T(LDLR), p.A242T(LRP5) | | |
| 6; ø | 3 | p.L432V(LDLR), p.L479P(LDLR), p.L473F(LRP4) | Loss of Leu? | moderate |
| 7 | 5 | p.D433H(LDLR), p.V523M(LDLR), p.T244(LRP5), p.A(LRP5), p.G(LRP5) | | |
| 8; ø | 4 | p.T434K(LDLR), p.L568V(LDLR), p.L(LRP4), p.Q(LRP4) | | |
| 9; d | 6 | p.D482H(LDLR), p.D203N(LRP5), p.D381N(LRP5), p.D511A(LRP5), p.D683N(LRP5), p.T852M(LRP5) | Loss of Asp | high |
| 10 | 4 | p.W483R(LDLR), p.P526S(LDLR), p.V684A(LRP5), p.N1121D(LRP5) | | |
| 11 | 0 | | | |
| 12 | 0 | | | |
| 13 | 0 | | | |
| 14 | 2 | p.R574C(LDLR), p.R574H(LDLR) | | |
| 15; ø | 1 | p.A399D(LDLR) | | moderate |
| 16; **Y** | 4 | p.Y442H(LDLR), p.Y733H(LRP5), p.Y1168H(LRP5) p.Y2522H(LRP2)[27] | Loss of Tyr | high |
| 17; **W** | 3 | p.L401V(LDLR), p.W577G(LDLR), p.W577S(LDLR) | Loss of Trp | high |
| 18; **T** | 3 | p.T253I(LRP5), p.T390K(LRP5), p.G520V(LRP5) | Loss of Thr | high |
| 19; **D** | 7 | p.F403L(LDLR), p.D492H(LDLR), p.D579N(LDLR), p.D579Y(LDLR), p.D434N(LRP5), p.D529N(LRP4)[39], p.D1403H(LRP4)[39] | Loss of Asp | high |
| 20(sbin) | 6 | p.T404P(LDLR), p.A214T(LRP5), p.A214V(LRP5), p.W478R(LRP5), p.A522T(LRP5), p.W1186S(LRP4) | | moderate |
| 21 | 2 | p.G171R(LRP5), p.R348W(LRP5) | | |
| 22 | 0 | | | |
| 23 | 0 | | | |
| 24 | 2 | p.R406W(LDLR), p.T173M(LRP5) | | |
| 25 | 1 | p.E1233K(LRP4) | | |
| 26 | 2 | p.E408K(LDLR), pD449N(LRP4) | | |
| 27; **I** | 3 | p.I451T(LDLR), p.I585T(LDLR), p.I450V(LRP4)[39] | Loss of Ile | high |
| 28 | 2 | p.R353Q(LRP5), p.E441K(LRP5) | | |
| 29; **R** | 4 | p.R570Q(LRP5), p.R570W(LRP5), p.R1277H(LRP4), p.R473Q(LRP6) | Loss of Arg | high |
| 30 | 2 | p.T454N(LDLR), p.A400E(LRP5) | | |
| 31 | 3 | p.S356L(LRP5), p.R444C(LRP5), p.N531I(LRP5) | | |
| 32 | 2 | p.L414R(LDLR)), p.R633C(LDLR) | | |
| 33 | 0 | | | |
| 34; d/n | 4 | p.D415G(LDLR), p.G592E(LDLR), p.D1099Y(LRP5), p.N433S(LRP6) | | |
| 35; **G** | 3 | p.R416W(LDLR), p.G549D(LDLR), p.G404R(LRP5) | | high |
| 36 | 1 | p.T535M(LRP5) | | |
| 37 | 0 | | | |
| 38; R | 5 | p.R595W(LDLR), p.R494Q(LRP5), p.R752G(LRP5), p.R1188W(LRP5), p.R632H(LRP4)[58] | Loss of Arg | high |
| 39 | 2 | p.V639D(LDLR), p.T461P(LRP4) | | |
| 40 | 2 | p.T798A(LRP5), p.T409A(LRP5) | | |
| 41; ø | 2 | p.L145F(LRP5), p.L540P(LRP5) | | moderate |

| | | | | |
|---|---|---|---|---|
| 42; ø | 2 | p.I423T(LDLR), p.V454M(LRP5) | | moderate |
| 43 | 0 | | | |
| 44 | 1 | p.D601H(LDLR) | | |
| 45 | 0 | | | |
| 46 | 0 | | | |
| 47; **L** | 0 | | | moderate |
| | **109** | | | |

### 3 The EGF-domain (residues 1014-1074)

#### 3.a Sequence details

EGF-domains are widely present in proteins with diverse biological functions [59]. This domain typically has ~40 amino acids with several short β-strands containing conserved cysteines invariably forming intradomain disulfide bridges [59]. In the mammalian proteome, EGF-domains are commonly divided into two subgroups: (1) those containing eight cysteines, which often occur in proteins of the extracellular space, (i.e. Laminin, Fibrillin, and the β-subunit of integrins) and (2) those with six cysteines commonly found in LDLRs. Indeed, the YWTD β-propellers in LDLR, LRP4, LRP6, and ApoER2 are most often flanked with 1 or 2 *six*-cysteine-EGF-domains before, and always by at least one *six*-cysteine EGF-domain at their C-terminal end [15,60] (**Fig. 1**). While SORLA has historically been acknowledged as a member of the LDLR family [10,61,62] it includes only one *eight*-cysteine EGF-domain C-terminal to the β-propeller and none before (**Fig. 1, Fig. 3**). Moreover, the SORLA EGF-domain contains 61 amino acids (residues 1014-1074; encoded by a single exon 22) such that it is substantially larger than EGF-domains in other LDLR family members. (**Supplemental Information 3d**). Only a single amino acid separates the fourth and fifth cysteines and there is also only a single residue between the sixth and the seventh cysteines of the SORLA domain [63]. The disulfide connectivity for integrin-type domains is $Cys^{I}$-$Cys^{V}$, $Cys^{II}$-$Cys^{IV}$, $Cys^{III}$-$Cys^{VI}$, and $Cys^{VII}$-$Cys^{VIII}$. While, based on the number of the cysteine residues and their spacing in the sequence, the SORLA EGF-domain might resemble the "integrin-like" type, there is little similarity between the SORLA EGF-domain and the EGF-domains from integrin β-subunits (**Supplemental Information 3d**). In contrast, several residues from the SORLA-EGF domain can be aligned with the sequence of EGF-domains positioned C-terminal to YWTD β-propellers from the LDLR family with the highest sequence similarity between the two last cysteine residues (**Supplemental Information 3d**). This suggests that the SORLA EGF-domain is evolutionary related to the LDLR family: the connectivity of the outer six cysteines follows the stereotypic pattern of LDLR EGF-domains, and the two additional central cysteines ($Cys^{IV}$ and $Cys^{V}$) could allow the formation of a loop onto the regular EGF-domain fold (**Supplemental Figure S3b**).

The solved structure of the YWTD-EGF domain pair from LDLR indicated that the C-terminal EGF-domain stabilizes the 6-bladed YWTD β-propeller. The EGF-domain forms contacts with the propeller through a series of tight hydrophobic interactions with the linker region and the YWTD-domain that assist to fold the combined two-domain structure, analogous to the association between 10CC- and VPS10p-domain pairs. Indeed, functional studies demonstrated that it was impossible to isolate the propeller without the EGF-domain of LDLR [15]. In LDLR, the EGF-A domain forms a protein-protein interaction with the catalytic domain of the protein PCSK9, which assists in the endocytosis and subsequent lysosomal degradation of LDLR in the liver [64].

Andersen
Supplemental Figure S3 - EGF-domain

**a**

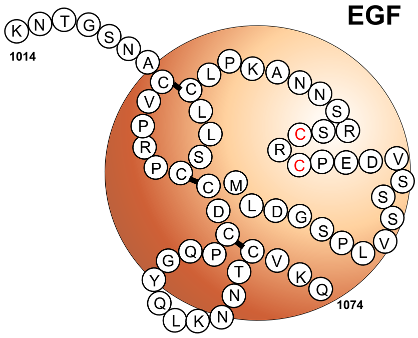

**b**

disulfides as in integrin-type EGF-domains

```
EGF  1014-1074  KNTGSNACV-----PRPCSLLCLPKANNSRSCRCPEDVSSSVLPSGDLMCDCPQGYQLKNN--TCVKQ   SORLA

                   GVNWCERTTLSNGGCQYLCLPAPQI-------------NPHSPKFTCACPDGMLLARDMRSCL   LDLR
                   -PDACELSVQPNGGCEYLCLPAPQI-------------SSHSPKYTCACPDTMWLGPDMKRCY   APOER2
                   GKNWCEED-MENGGCEYLCLPAPQI-------------NDHSPKYTCSCPSGYNVEENGRDCQ   VLDLR
                   ATNPCGID---NGGCSHLCLMSPVK-----------------PFYQCACPTGVKLLENGKTCK   LRP6_1
                   GSNPCAEE---NGGCSHLCLYRPQG-------------------LRCACPIGFELISDMKTCI   LRP6_2
                   GWNECASS---NGHCSHLCLAVPVG------------------GFVCGCPAHYSLNADNRTCS   LRP6_3
                   RQHPCAQD---NGGCSHICLVKGDG------------------TTRCSCPMHLVLLQDELSCG   LRP6_4
                   GKNRCGDN---NGGCTHLCLPSGQN-------------------YTCACPTGFR-KISSHACA   LRP4_1
                   VSTPCAME---NGGCSHLCLRSPNP-----------------SGFSCTCPTGINLLSDGKTCS   LRP4_2
                   GFNKCGSR---NGGCSHLCLPRPSG-------------------FSCACPTGIQLKGDGKTCD   LRP4_3
                   GTNACGVN---NGGCTHLCFARASD-------------------FVCACPD----EPDSRPCS   LRP4_4
                   FHTRCEED---NGGCSHLCLLSPSE-----------------PFYTCACPTGVQLQDNGRTCK   LRP5_1
                   GTNPCADR---NGGCSHLCFFTPHA-------------------TRCGCPIGLELLSDMKTCI   LRP5_2
                   GLNDCMHN---NGQCGQLCLAIPGG-------------------HRCGCASHYTLDPSSRNCS   LRP5_3
                   SAHPCARD---NGGCSHICIAKGDG------------------TPRCSCPVHLVLLQNLLTCG   LRP5_4
                   G-N-C--D---NGGCS-LCLP-P---------------------Φ-C-CP-GΦ-L--D--TC-
```

disulfides as in LDLR EGF-domains

#### 3.b Supplemental Figure S3. EGF-domain

**a**. Schematics of the EGF-domain of SORLA (orange circle), depicting residues 1014-1074 by their one-letter-code with indication of the three speculated disulfides, and the possible long loop including the extra pair of cysteines (the fourth and fifth Cys out of the eight, in red).

**b**. Alignment of SORLA residues 1014-1074 with sequences of EGF-domains located at the C-terminal of β-propellers of members from the LDLR family (as presented in **Fig. 2**). All cysteines are shown in red, and residues at positions with strong (capital letters) or weak (lower case letters) conservation are shown below the alignment.

On the top of the SORL1 sequence is indicated the Cys connectivity for the eight Cys residues located within integrin-type EGF-domains (as shown in **Supplemental Information 3d**).

#### 3.c *SORL1* variants in EGF-domain

The importance of the 61-residue EGF-domain in SORLA as a whole is underscored by a pedigree for a Swedish AD-family carrying variant 11:121437647, which translates to c.3050-2A>G that leads to loss of adjacent splice acceptor site and exclusion of exon 22 – and thus deletion of the entire EGF-domain (p.Gly1017-Glu1074del) [65]. We speculate that lack of the EGF-domain is likely not compatible with folding of a functional receptor, which may also explain the observed co-occurrence of the YWTD-domain with a neighboring EGF-domain at its C-terminal end in LDLRs. Due to the limited sequence similarity between the SORLA EGF-domain and the EGF-domains in the LDLR proteins it was not possible to accurately assess pathogenic variants in Uniprot that map to the SORLA EGF-domain.

#### 3.d EGF-domain alignment

We prepared alignments of the SORLA EGF-domain sequence with homologous domains from LDLR or from Integrin beta2 and beta4. The SORLA EGF-domain share overall more similarly to YWTD- than integrin-like EGF-domains, despite the presence of 8 Cys residues in the SORLA domain and ony 6 Cys in the LDLR-type EGF-domains. The color code indicates partial amino acid conservation across the domain sequence.

**Alignment between EGF-domain sequences located C-terminal to YWTD-propellers (see Fig 2)**

```
LRP6_1:    ATNPCGID---NGGCSHLCLMSPVK-----------------PFYQCACPTGVKLLENGKTCK
LRP6_2:    GSNPCAEE---NGGCSHLCLYRPQG-------------------LRCACPIGFELISDMKTCI
LRP6_3:    GWNECASS---NGHCSHLCLAVPVG------------------GFVCGCPAHYSLNADNRTCS
LRP6_4:    RQHPCAQD---NGGCSHICLVKGDG------------------TTRCSCPMHLVLLQDELSCG
LRP4_1:    GKNRCGDN---NGGCTHLCLPSGQN-------------------YTCACPTGFR-KISSHACA
LRP4_2:    VSTPCAME---NGGCSHLCLRSPNP-----------------SGFSCTCPTGINLLSDGKTCS
LRP4_3:    GFNKCGSR---NGGCSHLCLPRPSG-------------------FSCACPTGIQLKGDGKTCD
LRP4_4:    GTNACGVN---NGGCTHLCFARASD-------------------FVCACPD----EPDSRPCS
LRP5_1:    FHTRCEED---NGGCSHLCLLSPSE-----------------PFYTCACPTGVQLQDNGRTCK
LRP5_2:    GTNPCADR---NGGCSHLCFFTPHA-------------------TRCGCPIGLELLSDMKTCI
LRP5_3:    GLNDCMHN---NGQCGQLCLAIPGG-------------------HRCGCASHYTLDPSSRNCS
LRP5_4:    SAHPCARD---NGGCSHICIAKGDG------------------TPRCSCPVHLVLLQNLLTCG
LDLR:      GVNWCERTTLSNGGCQYLCLPAPQI-------------NPHSPKFTCACPDGMLLARDMRSCL
APOER:      PDACELSVQPNGGCEYLCLPAPQI-------------SSHSPKYTCACPDTMWLGPDMKRCY
VLDLR:     GKNWCEED-MENGGCEYLCLPAPQI-------------NDHSPKYTCSCPSGYNVEENGRDCQ
SORLA:  KNTGSNACV-----PRPCSLLCLPKANNSRSCRCPEDVSSSVLPSGDLMCDCPQGYQLKNNT--CVKQ
```

**Alignment with EGF-domains from integrin beta2 and beta4 following a published alignment of these domains in refs** [63,66]

```
Beta2_1:      CR---------DQSR----DRSLCHG----KGFLEC-------------GICRCDT----GY--IGKNCE
Beta2_2:      CQTQGRSSQELEGSCRKDNNSIICSG----LGDCVC-------------GQCLCHTSDVPGKLIYGQYCE
Beta2_3:      CD---------TINCERY-NGQVCGG--PGRGLCFC-------------GKCRCHP----GF--EGSACQ
Beta2_4:      CER-------TTEGCLNP-RRVECSG----RGRCRC-------------NVCECHS----GY--QLPLCQ
Beta4_1:      CE--------LQKEV----RSARCSF----NGDFVC-------------GQCVCSE----GW--SGQTCN
Beta4_2:      CST-G--SLSDIQPCLREGEDKPCSG----RGECQC-------------GHCVCYGE---GR-YEGQFCE
```

```
Beta4_3:       Y---------DNFQCPRT-SGFLCND----RGRCSM-------------GQCVCEP----GW--TGPSCD
Beta4_4:       CPL-------SNATCIDS-NGGICNG----RGHCEC-------------GRCHCHQQ----SLYTDTICEINYS
SORLA: KNTGSNACV---------PRPCS-----LLCLPKANNSRSCRCPEDVSSSVLPSGDLMCDCPQ----GYQLKNNTCVKQ
```

### 4 The CR-cluster (residues 1075-1550)

#### 4.a Sequence details

More than half of all the ligands identified to bind the SORLA receptor, including APP, interact with the CR-cluster [67-69]. In fact, SORLA that lacks all its eleven CR-domains fails to bind APP [70], making it unlikely that APP binds to regions outside the CR-cluster.

CR-clusters of LDLR family members all contain 2 to 12 CR-domains (**Fig. 2**), while proteins that belong to the complement system include single CR-domains (i.e. C6, C7, C8α, C8β, C9, factor 1). This has led to two different definitions of the same module: "LDLR-type A repeats" and "Complement-type repeat (CR)-domains". One CR-domain sequence contains approximately 40 amino acids with several highly conserved residues, including six cysteines and five conserved acidic residues. The 6 cysteines (positions 15, 23, 29, 36, 42, 55) form three invariable disulfide bridges with the connectivity $Cys^{I}$-$Cys^{III}$, $Cys^{II}$-$Cys^{V}$, and $Cys^{IV}$-$Cys^{VI}$ [71,72] (**Supplemental Figure S4)**, whereas the 5 conserved acidic residues were originally thought to engage in binding of ligands containing exposed basic residues in their receptor-binding site. In addition, a serine (position 46), together with a pair of hydrophobic residues at positions 21 (phenylalanine) and position 30 (isoleucine) in the more N-terminal part of the sequence are also highly conserved across CR-domains (**Supplemental Information 4d**). Furthermore, the CR-domains in SORLA also contain a pair of glycines (positions 27 and 38) that is conserved in eight of the eleven CR-domains (**Fig. 3**). With this high sequence conservation for 16 out of 40 positions across CR-domains from several proteins, it is not surprising that they all show a very similar folding (including domains from LDLR, ApoER2, LRP1, and LRP2) [73-81]. Structure determination by NMR indicates a very compact folding consisting of a β-hairpin structure with two short strands, followed by a series of β-turns (**Supplemental Figure S4**). The glycine at position 27 is at the center of the β-hairpin-turn, which requires a small side chain or none at all. The conserved phenylalanine (position 21) and isoleucine (position 30) pack against each other in a small hydrophobic core of the domain interior, preventing their side chains from engaging in ligand interactions (**Supplemental Figure S4**).

In members of the LDLR family, the combination of CR-domains plays a key role in ligand binding [82], outlining a functional consequence of exon skipping in some CR-clusters. This is best exemplified by the human ApoER2 gene (aka LRP8), which can encode a receptor with eight different CR-domains [83]. Most of the translated ApoER2 molecules *lack* three central CR-domains CR4-CR6, producing a receptor that binds efficiently to Reelin but not $\alpha_2$-

macroglobulin. However, when CR4-CR6 is included this longer ApoER2 isoform can also bind $\alpha_2$-macroglobulin [84]. Another ApoER2 isoform, with an unknown effect on ligand binding, lacks the most C-terminal CR-domain due to a second alternative splice event. Similarly, exon 4 skipping of human very low-density lipoprotein receptor *VLDLR* leads to a receptor isoform that lacks the third CR-domain, which increases VLDLR-affinity for ApoE-lipoproteins compared to VLDLR containing all its eight CR-domains [85]. Accordingly, alternative splicing of exons encoding CR-domains presents a mechanism to generate receptor variants with unique patho/physiological ligand binding properties.

*A conserved Calcium cage - and the minimal motif*

Ligand-binding to members of the LDLR family is critically dependent on calcium ions, which are coordinated by four of the conserved acidic residues in each CR-domain (at positions 37, 41, 47, and 48) (**Supplemental Figure S4a, red arrows in panel c**). Their acidic side chains form an octahedral calcium cage [75], which also stabilizes the folding of the C-terminal part of the domain [86]. The side chain of a fifth conserved acidic residue (aspartate at position 44) forms a structure known as an “Asx-turn”: it makes a hydrogen bond with the backbone amides of two residues: one residue upstream and the conserved serine two residues downstream (position 46) [87]. Two additional amino acids (positions 34 and 39) contribute to the calcium coordination with their backbone carbonyl groups (**Supplemental Figure S4a, and black arrows in panel c**). This geometry makes the side chains of residues at these two positions (most often a Trp-Asp pair in LDLR family proteins) ideally positioned at the domain’s molecular surface to engage in calcium-dependent ligand interactions [88-90]. These two amino acids, at positions 34 and 39, have therefore been named “CR-domain fingerprint residues” [23]. APP binding to SORLA Is also strictly dependent on the presence of calcium [67].

The side chains of the two fingerprint residues interact most often with a lysine residue of the ligand (often positioned on a helical structure [91]) and a residue containing a hydrophobic side chain. It is intriguing how such a simple motif, called the “the minimal motif” [23,81,92], allows for discrimination of interaction partners [81,93-96]. The fingerprint of the SORLA CR-cluster is strongly conserved among all the 11 CR-domains across evolution, highlighting the importance of the CR-cluster function (**Supplemental Information 9**). Substitutions at positions of fingerprint residues can impair binding of specific ligands, but do not affect overall folding and stability of CR-domains [88-90,97].

The CR-domain fingerprint residues of five of the eleven CR-modules in SORLA (CR1, CR2, CR3, CR4, and CR8) are represented by the canonical Trp-Asp pair of amino acids common to the receptor:ligand complexes for the LDLR family (**Supplemental Figure S4a**). The remaining six pairs of fingerprint residues have a hydrophobic residue instead of the aspartate, and three CR-domains (CR5, CR10 and CR11) have a charged residue (Glu or Lys) instead of the Trp, suggesting that the SORLA CR-cluster may also bind ligands with motifs different from the common lysine-based motif for ligands of LDLR family members. The SORLA ligand profile of these CR-domains may be more similar to SCO-spondin which contains 10 CR-domains, none of which has a Trp-Asp fingerprint pair [98]. Alternatively, the binding partner for these CR-domains may be another part of the SORLA receptor, maybe depending on its overall folded conformation.

*The necklace model and linker length*

CR-domains fold independently from neighboring CR-domains, and substitutions that cause local misfolding of one CR-domain (e.g., substitutions of residues in the Calcium cage) still allow for correct folding of its adjacent CR-domain *in vitro* [99-101]. This agrees with the observed negligible interdomain interactions [21,102,103], suggesting that CR-clusters are like a necklace with the individual CR-domains behaving as “pearls-on-a-string” [102]. This modular organization allows for a high degree of flexibility that seems primarily determined by the length (and eventually the composition) of the interdomain sequence of amino acids; ie.e., the so-called linkers. This flexibility enables different CR-domains to wrap around larger ligands and engage in minimal motif interactions with multiple sites of the ligand. Such avidity, including two or more receptor CR-domains, leads to high-affinity ligand binding [88,104]. The linker sequences of CR-clusters are commonly 3 to 4 residues, with 12 amino acids as the longest connective string in LDLR. Many linker sequences in the SORLA CR-cluster are longer, and the linkers towards the most C-terminal part of the cluster are extremely long. The sequences connecting CR8 with CR9 and CR9 with CR10 contain 15 and 17 amino acids, respectively, suggesting a unique flexibility of the SORLA CR-cluster. However, as these linker-sequences are also highly conserved among species (**Supplemental Information 9**), we hypothesize that they fulfill an important role in the physiological function of SORLA. Interestingly, some of the SORLA linkers become modified by *O*-linked glycans, including residues T1198 and T1508 [105,106], but how that relates to receptor activity has not yet been determined.

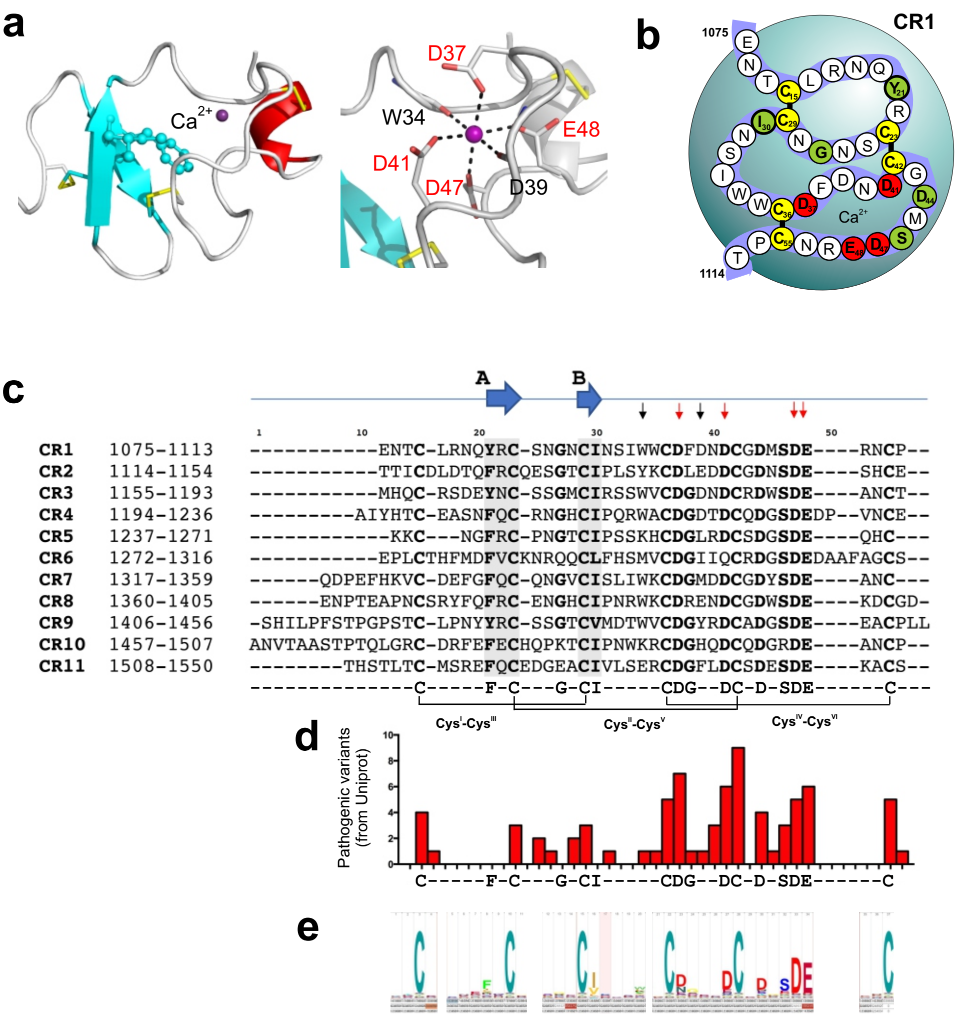
a
Ca2+
W34
D37
E48
D41
D47
D39
b
CR1
1075
1114
Ca2+
c
A
B
CR1 1075-1113 -----------ENTC-LRNQYRC-SNGNCINSIWWCDFDNDCGDMSDE----RNCP--
CR2 1114-1154 -----------TTICDLDTQFRCQESGTCIPLSYKCDLEDDCGDNSDE----SHCE--
CR3 1155-1193 -----------MHQC-RSDEYNC-SSGMCIRSSWVCDGDNDCRDWSDE----ANCT--
CR4 1194-1236 ---------AIYHTC-EASNFQC-RNGHCIPQRWACDGDTDCQDGSDEDP--VNCE--
CR5 1237-1271 -------------KKC---NGFRC-PNGTCIPSSKHCDGLRDCSDGSDE----QHC---
CR6 1272-1316 -----------EPLCTHFMDFVCKNRQQCLFHSMVCDGIIQCRDGSDEDAAFAGCS--
CR7 1317-1359 ------QDPEFHKVC-DEFGFQC-QNGVCISLIWKCDGMDDCGDYSDE----ANC---
CR8 1360-1405 ------ENPTEAPNCSRYFQFRC-ENGHCIPNRWKCDRENDCGDWSDE----KDCGD-
CR9 1406-1456 -SHILPFSTPGPSTC-LPNYYRC-SSGTCVMDTWVCDGYRDCADGSDE----EACPLL
CR10 1457-1507 ANVTAASTPTQLGRC-DRFEFECHQPKTCIPNWKRCDGHQDCQDGRDE----ANCP--
CR11 1508-1550 --------THSTLTC-MSREFQCEDGEACIVLSERCDGFLDCSDESDE----KACS--
--------------C-----F-C---G-CI-----CDG--DC-D-SDE------C---
$Cys^{I}$-$Cys^{III}$
$Cys^{II}$-$Cys^{V}$
$Cys^{IV}$-$Cys^{VI}$
d
Pathogenic variants (from Uniprot)
C-----F-C---G-CI-----CDG--DC-D-SDE------C
e

#### 4.b Supplemental Figure S4. CR-domains

**a**. The structure of CR7 from LRP1 (PDB: 1J8E) [78] with two short strands (light blue) and a short $\alpha$-helix (pink). Close-up of the octahedral coordination of a $Ca^{2+}$ ion (purple) by side chain carboxylates of D37, D41, D47, and E48 as well as backbone carbonyls from residues at position 34 and 39 (numbering according to sequence position in panel **c**).

**b**. Schematic of the residues 1075-1114 of CR1 of SORLA with conserved residues on a colored background: cysteines (yellow), $Ca^{2+}$ coordination via side chain carboxylates (red), and other highly conserved positions (green).

**c**. Alignments of the eleven CR-domains of SORLA (residues 1075-1550). Conserved residues including six cysteines, two hydrophobic amino acids at positions 21 and 30, and 5 acidic residues and a Ser are all indicated below the alignment. Red and black arrows on top of the alignment indicate positions involved in calcium chelation via side chain or backbone carbonyls, respectively. Horizontal lines indicate the locations of the three invariable disulfides.

**d.** Histograms showing the number of pathogenic variants that occur for each CR-domain position as listed in **Supplemental Information 4f**.

**e.** Logo representation of the domain sequence conservation: the larger the letter the higher the conservation across CR-domain sequences.

#### 4.c *SORL1* variants in CR-domains

*Odd-numbered cysteines (ONC, positions 15, 23, 29, 36, 42, and 55 + all):*

Either removal of one of the six conserved cysteines or introduction of a cysteine residue at another position of the domain sequence may disrupt the disulfide connectivity of CR-domains. There is strong evidence from our DMDM analysis that variants leading to an odd number (5 or 7) of cysteines (ONC) result in dysfunctional folding for other proteins with CR-clusters (**Supplemental Information 4f**). Most prominently, we found that 21 out of 51 positions in LDLR linked to FH involve the replacement of a cysteine residue within the CR-cluster (**Supplemental Information 4e**). Also the introduction of an extra cysteine, p.R78C$^{LDLR}$ is considered pathogenic [51]. The DMDM analysis also identified disease associated variants that involve replacement of cysteines in other LDLR family members: p.C160Y$^{LRP4}$ causal for CLSS [38] and p.C1361G$^{LRP5}$ in patients with EVR4 [28,29]. Such ONC mutations are also pathogenic in proteins outside the LDLR family, e.g. the transmembrane proteinase TMPRSS6, (p.C510S$^{TMPRSS6}$ [107,108] and p.C510R$^{TMPRSS6}$ [107] in patients with IRIDA) or the COP9 protein (p.C119G$^{C9}$ [109] in patients with C9D).

Two different ONC variants in *SORL1* has lately been reported to segregate with AD in small pedigrees that indicate that such mutations are truly pathogenic. First, a Swedish family (PED.25) with segregating variant p.R1303C in the sixth CR-domain was identified [65], and later a family in Saudi Arabia with variant p.R1084C in the first CR-domain was reported [110], both resulting in CR-domains with 7 cysteines. Also, a 59-year-old AD patient with variant p.C1192Y that results in having 5 cysteines in the third CR-domain has been identified [111], supporting that ONC variants in *SORL1* is associated with high risk for AD. The variant p.C1344R was reported associated with a possible family history in Finland [65], and also variants p.C1453S and p.C1249S were identified exclusively in AD patients [112].

Additional analysis of ONC variants in *SORL1* from the ADES-ADSP dataset [40], show how they occur predominantly in AD cases (p.R1080C, p.W1095C, pC1112Y, p.R1124C, p.R1172C, p.C1177Y, p.C1196CY, p.R1243C, p.R1260C, p.C1275S, p.C1286C, p.R1303C, p.Y1371C, p.Y1424C, p.Y1441C, p.C1453F, p.C1478S, p.R1490C, p.C1521R, p.C1540S; n=31) compared to controls (p.Y1196C, p.R1490C, p.C1497Y; n=5). Hence, ONC substitutions associate with a very strong increased risk of AD (OR = 6.31 95% CI: 2.45 - 16.24, p=5.1E-6; Fisher Exact test).

We note the similarity between ONC variants in *SORL1* associated with AD and variants in *NOTCH3* causal of Cerebral Autosomal Dominant Arteriopathy with Subcortical Infarcts and

Leukoencephalopathy (CADASIL), where stereotypic causal variants also result in an odd-number of cysteines in EGF-domains of NOTCH3 carrying 32-34 copies of this domain type [113]. Or how 22 of 80 cysteines from the sequence of ten EGF-domains (of the eight-cysteine type) from the Usherin protein (encoded by *USH2A* are found mutated in patients with retinitis pigmentosa (*USH2A* LOVD mutation database, http://www.lovd.nl/USH2A). This demonstrates a general mechanism how variants in small cysteine-rich protein disulfide-containing domains that affects the number of cysteine residues may associate with a very high disease penetrance, and suggest that also ONC variants in *SORL1* should with high confidence be considered pathogenic and causal of AD.

*Calcium cage (CaCa, positions 37,41, 47, and 48):*

In proteins with CR-domains, residues at positions 37 and 41 and 47 are almost all Asp (in SORLA, there is a single exception; $Gln^{1301}$ at pos 41 in CR6) and positions 48 are all Glu (**Fig. 2e**). The side chains of these residues coordinate $Ca^{2+}$ establishing an octahedral $Ca^{2+}$ cage (CaCa) that is critical for domain folding [101,114]. As a consequence, substitutions of these variants may be strongly associated with disease.

Our DMDM analysis showed that in LDLR, substitutions of CaCa residues are associated with FH; **position 37**: p.D90G [51], p.D90N [51], p.D90Y [115], p.D168A [34], p.D168H [116], p.D168N [51,117], p.D168Y [118], p.D172N [30,117], p.D221G [26,34,51,52,119,120], p.D221N [42,52], p.D221Y [119,120]; **position 41**: p.D301G [34,117,121], p.D301A [122]; **position 47**: p.D139H, p.D227E [51,123], p.D266E [26]; and **position 48**: p.E101K [51,52], p.E140K [30,115,124]; p.E228K [34,125], p.E228Q [34]. Note the two disease-associated substitutions of Asp with Glu at position 47, which is generally considered a conservative – and often non-pathogenic – substitution. However, in CR-domains there is not enough space in the Calcium cage to accommodate the larger glutamate side chain at position 47 [75]. Uniprot also lists disease-associated variants for CaCa positions in other proteins: a variant in LRP2 at position 37 associates with intellectual disability ($p.D3779N^{LRP2}$) [126] and another LRP2 variant at position 47 causes Stickler syndrome ($p.D3828G^{LRP2}$) [127]. Uniprot further lists disease-associated variants in LRP5 ($p.E1367K^{LRP5}$ in patients with EVR [28,49]), TMRPSS3 ($p.D103G^{TMPRSS3}$ causal of deafness [128,129]) and TMRPSS6 (variants $p.D521G^{TMPRSS6}$ [107], $p.D521N^{TMPRSS6}$ [130,131] and $p.E522K^{TMPRSS6}$ [131] considered causative of IRIDA) (**Supplemental Information 4e**).

There is also evidence that CaCa variants in *SORL1* is associated with AD. Studies report carriers of variant p.D1389V (position 37, CR8) only in AD patients, and these have very early onset (<51 years) [132-134]. Other studies identified variant p.D1545E of CR11 (position 47; CADD score 15.9) [132], p.D1182N (position 41, CR3), and p.D1267E (position 47, CR5) [112] in

AD patients only. Interestingly, the p.D1267E and p.D1545E variants have very low CADD scores (~16) presumably because a substitution of an Asp with a Glu is generally considered benign, only not when present in a CR-domain Calcium cage. We also recently found that the CaCa p.D1545V (position 47, CR11) variant is acting as a dominant negative and causal variant of AD in an Icelandic family [135].

The ADES-ADSP dataset includes such variants (p.D1108N, p.D1219G, p.D1261G, p.D1267N, p.D1345N, p.D1389V, p.D1502G, p.D1535N, p.D1545N, p.D1545G, p.D1545E) at positions in Calcium cages exclusively in AD cases (n=13) with a relatively early age at onset (median 60 years, ranging from 47-73 years) such that they are associated with a strong increased risk of AD (OR = INF) and in practice should be considered as causative for AD as loss-of-function variants that is now accepted as causal for AD [136].

*Asx-turn: aspartate at position 44:*

In SORLA all eleven CR-domains contain an aspartate at position 44, which forms the Asx-turn [75]. Functional studies indicated that in LDLR, even the slightest modification of this residue results in FH, exemplified by the conservative aspartate to asparagine mutation in an LDLR CR-domain, because both carboxylate oxygens of the aspartate are necessary for hydrogen bonding [75]. The large alignment of CR-domain sequences confirms a strong preference for aspartate at position 44, and it may thus be considered a hotspot for pathogenic mutations (**Supplemental Information 4f**). Uniprot lists four disease-associated variants at this position: p.D175N$^{LDLR}$ (causal of FH in Afrikaners [123]), p.D175Y$^{LDLR}$ (in FH patients [137]), p.D224V$^{LDLR}$ (causal of elevated LDL cholesterol in FH patients [119]), and p.D137N$^{LRP4}$ (variant is considerd causal of CLSS [38]).

In the ADES-ADSP dataset, we observed variants p.D1105H and p. D1146N at this position in *SORL1*, in respectively a 64- and a 48-year-old AD patient and none in controls [40]. Functional tests in cell culture studies showed strongly reduced shedding of D1105H mutant SORLA, and found low sSORLA levels in CSF from two carriers of the variant, supporting these *SORL1* variants as pathogenic (Holstege and Andersen, unpublished).

*Hydrophobic “core”: phenylalanine and Isoleucine at positions 21 and 30*

These two positions in the CR-domain sequence are strongly conserved and stabilize the N-terminal part of CR-domains [73,74]. *In vitro* studies showed that mutation of residues at these two positions destabilized the CR-domain folding and impaired ligand-binding activity of LDLR [138]. Surprisingly, our DMDM analysis of disease-associated variants in Uniprot did not find any pathogenic variants for neither of these two positions in any CR-domain containing proteins, including LDLR (**Supplemental Information 4f**). This is surprising but suggests that despite being positions with strongly conserved amino acids, it may not be dangerous to substitute with other residues at these positions.

#### 4.d CR-domain alignment

Mapping of naturally occurring variants found in human CR-domain containing proteins, and being listed as associated with pathology at www.uniprot.org. The alignment follows the SORLA alignment shown on top, and pathogenic variants are highlighted on a red background.

```
                    1        10        20        30        40        50
CR1     1075-1113   -----------ENTC-LRNQYRC-SNGNCINSIWWCDFDNDCGDMSDE----RNCP--
CR2     1114-1154   -----------TTICDLDTQFRCQESGTCIPLSYKCDLEDDCGDNSDE----SHCE--
CR3     1155-1193   -----------MHQC-RSDEYNC-SSGMCIRSSWVCDGDNDCRDWSDE----ANCT--
CR4     1194-1236   ---------AIYHTC-EASNFQC-RNGHCIPQRWACDGDTDCQDGSDE--DPVNCE--
CR5     1237-1271   ------------KKC---NGFRC-PNGTCIPSSKHCDGLRDCSDGSDE----QHC---
CR6     1272-1316   -----------EPLCTHFMDFVCKNRQQCLFHSMVCDGIIQCRDGSDEDAAFAGCS--
CR7     1317-1359   ------QDPEFHKVC-DEFGFQC-QNGVCISLIWKCDGMDDCGDYSDE----ANC---
CR8     1360-1405   ------ENPTEAPNCSRYFQFRC-ENGHCIPNRWKCDRENDCGDWSDE----KDCGD-
CR9     1406-1456   -SHILPFSTPGPSTC-LPNYYRC-SSGTCVMDTWVCDGYRDCADGSDE----EACPLL
CR10    1457-1507   ANVTAASTPTQLGRC-DRFEFECHQPKTCIPNWKRCDGHQDCQDGRDE----ANCP--
CR11    1508-1550   --------THSTLTC-MSREFQCEDGEACIVLSERCDGFLDCSDESDE----KACS–
                                  C     F C   g CI      CDG  DC D SDE       C

LDLR:
                             VGDRC-ERNEFQC-QDGKCISYKWVCDGSAECQDGSDE--SQETCLS
Δ=GG                           VTC-KSGDFSCΔRVNRCIPQFWRCDGQVDCDNGSDE----QGCP
                              PKTC-SQDEFRC-HDGKCISRQFVCDSDRDCLDGSDE----ASCPV
                               LTC-GPASFQC-NSSTCIPQLWACDNDPDCEDGSDE--WPQRCRG
                       LYVFQGDSSPC-SAFEFHC-LSGECIHSSWRCDGGPDCKDKSDE----ENCA
                              VATC-RPDEFQC-SDGNCIHGSRQCDREYDCKDMSDE----VGCV
                             NVTLCEGPNKFKC-HSGECITLDKVCNMARDCRDWSDE--PIKECG
TMPRSS6:
                                PC--PGEFLCSVNGLCVPA---CDGVKDCPNGLDE----RNC
                                VC--RATFQCKEDSTCISLPKVCDGQPDCLNGSDE----EQCQ
Δ=PQ                            PC-GTFTFQC-EDRSCVKKPNΔCDGRPDCRDGSDE----EHCD
COP9:
                               DDC--GNDFQC-STGRCIKMRLRCNGDNDCGDFSDE----DDCES
LRP4:
Δ=ALGEC                         AC-GRSHFTCAVSΔTCIPAQWQCDGDNDCGDHSDE----DGCI
                              LPTC-SPLDFHC-DNGKCIRRSWVCDGDNDCEDDSDE----QDCP
                              PREC-EEDEFPC-QNGYCIRSLWHCDGDNDCGDNSDE-----QCD
                              MRKC-SDKEFRC-SDGSCIAEHWYCDGDTDCKDGSDE----ENCP
                          SAVPAPPC-NLEEFQC-AYGRCILDIYHCDGDDDCGDWSDE----SDCS
                             SHQPC-RSGEFMC-DSGLCINAGWRCDGDADCDDQSDE----RNCT
                              TSMC-TAEQFRC-HSGRCVRLSWRCDGEDDCADNSDE----ENCE
                           NTGSPQC-ALDQFLC-WNGRCIGQRKLCNGVNDCGDNSDE-SPQQNCRPR
LRP5:
Δ=AT                            TC-SPDQFACΔGEIDCIPGAWRCDGFPECDDQSDE----EGCP
                                VC-SAAQFPC-ARGQCVDLRLRCDGEADCQDRSDE----ADCD
                               AIC-LPNQFRC-ASGQCVLIKQQCDSFPDCIDGSDE----LMCEI
Corin:
                               LLCGRGENFLC-ASGICIPGKLQCNGYNDCDDWSDE----AHC
                                NC-SENLFHC-HTGKCLNYSLVCDGYDDCGDLSDE----QNC
                                DCNPTTEHRC-GDGRCIAMEWVCDGDHDCVDKSDE----VNC
                                SCHSQGLVEC-RNGQCIPSTFQCDGDEDCKDGSDE----ENCSV
                                EC-SPSHFKC-RSGQCVLASRRCDGQADCDDDSDE----ENC
                                GCKERDLWECPSNKQCLKHTVICDGFPDCPDYMDE----KNC
                               SFC-QDDELEC-ANHACVSRDLWCDGEADCSDSSDE----WDCVT
TMPRSS3:
                                DC--SGKYRCRSSFKCIELIARCDGVSDCKDGEDE----YRCV
CFAI:
                         VCYTQKADSPMDDFFQC-VNGKYISQMKACDGINDCGDQSDE----LCCK
                                AC-QGKGFHC-KSGVCIPSQYQCNGEVDCITGEDE----VGCA
```

#### 4.e Identity of mapped CR variants: disease proteins and disease variants

Summary of included proteins containing naturally occurring variants associated with diseases in domains shared with SORLA.

| Gene | Protein | Uniprot | #dom | Associated Diseases/Syndromes (CODE) | Variants mapped onto big alignments (from Uniprot entries except where a specific ref is provided) | #variants | #positions | Citations for variants discussed in main text |
|---|---|---|---|---|---|---|---|---|
| LDLR | Low-density lipoprotein receptor | P01130 | 7 | Familial hypercholesterolemia (FHCL1) | FHCL1: **C27W**, **C46S**, A50S/A50T, S56P, **R78C**, W87G, **C89Y**, **D90G/D90N/D90Y**, Q92E, **C95G**, **E101K**, **C116R**, **C134F/C134W**, **E140K**, **C143R**, **C148Y**, **C155Y**, **C160Y**, **D168A/D168H/D168N/D168Y**, **D172N**, **C173W**, **D175N/D175Y**, S177L, **C184W/C184Y**, **C197R**, H211L, **D221G/D221N/D221Y**, **C222Y**, **D224V**, **D227E**, **E228K/E228Q**, **C231G**, **C243R**, **C248Y**, Q254P, **C261F**, **D266E**, **C276R/C276W/C276Y**, E277K, H285Y, S286R, E288K, R300G, **D301G**, **C302W/C302Y**, S306L, **C313R**, G314R | 63 | 48 | **C27W** [51]<br>**C46S** [139]<br>**C89Y** [51,52]<br>**C95G** [120]<br>**C116R** [117,120]<br>**C134F** [37]<br>**C134W** [37]<br>**C143R** [140]<br>**C148Y** [140]<br>**C155Y** [31,121]<br>**C160Y** [34,51]<br>**C173W** [141]<br>**C184W** [140]<br>**C184Y** [34,142]<br>**C197R** [122]<br>**C222Y** [37]<br>**C231G** [143]<br>**C243R** [30]<br>**C248Y** [122]<br>**C261F** [144]<br>**C276R** [37]<br>**C276W** [34]<br>**C276Y** [34]<br>**C302W** [122]<br>**C302Y** [120]<br>**C313R** [51]<br>**D175N** causal in Afrikaners [123]<br>**D224V** causal of LDL cholesterol elevations [119] |
| LRP4 | Low-density lipoprotein receptor-related protein 4 | O75096 | 8 | Cenani-Lenz syndactyly syndrome (CLSS) | CLSS: **D137N**, **C160Y** | 2 | 2 | **D137N** and **C160Y** considered causal of CLSS [38] |
| LRP5 | Low-density lipoprotein receptor-related protein 5 | O75197 | 3 | Vitreoretinopathy, exudative 4 (EVR4) | EVR4: **C1361G**, **E1367K** | 2 | 2 | **C1361G** identified in patient with EVR4 [28,29]. |

| | | | | | | | | |
|---|---|---|---|---|---|---|---|---|
| | | | | | | | | Further evidence needed to establish if causal |
| TMPRSS6 | Transmembrane protease serine 6 | Q8IU80 | 3 | Iron-refractory iron deficiency anemia (IRIDA) | IRIDA: **C510R/C510S, D521G/D521N, E522K** | 5 | 3 | **C501R** [107]<br>**C501S** [107,108]<br>**D521G** [107]<br>**D521N** [130,131]<br>**E522K** [131] |
| TMPRSS3 | Transmembrane protease serine 3 | P57727 | 1 | Deafness, autosomal recessive, 8 (DFNB8) | **DFNB8**: **D103G** | 1 | 1 | **D103G** causal of deafness [128,129] |
| C9 | Complement component 9 | P02748 | 1 | Complement component 9 deficiency (C9D) | **C9D**: **C119G** | 1 | 1 | **C119G** [109] |
| CORIN | Atrial natriuretic peptide-converting enzyme | Q9Y5Q5 | 7 | Pre-eclampsia/eclampsia 5 (PEE5) | **PEE5**: K317E | 1 | 1 | |
| CFI | Complement factor I | P05156 | 2 | CFI deficiency<br>Hemolytic uremic syndrome atypical 3 (AHUS3) | **CFI deficiency**: G243D<br>**AHUS3**: G287R | 2 | 2 | |
| | | | | | | **77** | 60 | |

#### 4.f CR disease variants listed according to domain positions

Disease-mutations domain-mapping analysis with identification of pathogenic variants in other proteins with CR-domains (as listed in **Supplemental Information 4e**). Here variants are mapped onto domain positions following alignment of internally repeated sequences in the SORLA domain sequences. The number of hits for each position depicted in the bar diagram of **Supplemental Figure 4d**.

| **CR-domain positions** | Number of hits | Identified variants | | |
|---|---|---|---|---|
| 1 | | | | |
| 2 | | | | |
| 3 | | | | |
| 4 | | | | |
| 5 | | | | |
| 6 | | | | |
| 7 | | | | |
| 8 | | | | |
| 9 | | | | |
| 10 | | | | |
| 11 | | | | |
| 12 | | | | |
| 13 | | | | |
| 14 | | | | |
| 15; **C** | 4 | p.C27W(LDLR), p.C148Y(LDLR), p.C197Y(LDLR), p.C276Y(LDLR) | Loss of Cys | high |
| 16 | 1 | P:E277K(LDLR) | | |
| 17 | 0 | | | |
| 18 | 0 | | | |
| 19 | 0 | | | |
| 20 | 0 | | | |
| 21; **F** | 0 | | Loss of Phe/Tyr? | moderate |
| 22 | 0 | | | |
| 23; **C** | 3 | p.C155Y(LDLR), p.C116R(LDLR), p.C243R(LDLR) | Loss of Cys | high |
| 24 | 0 | | | |
| 25 | 2 | p.R78C(LDLR), p.H285Y(LDLR) | | |
| 26 | 1 | p.S286R(LDLR) | | |
| 27; G | 0 | | | |
| 28 | 2 | p.E288K(LDLR), p.K317E(CORIN) | | |
| 29; **C** | 3 | p.C160Y(LDLR), pC248Y(LDLR), p.C160Y(LRP4) | Loss of Cys | high |
| 30; **I** | 0 | | Loss of Ile? | moderate |
| 31 | 1 | p.H211L(LDLR) | | |
| 32 | 0 | | | |
| 33 | 0 | | | |
| 34; fp | 1 | p.W87G/LDLR) | | moderate |
| 35 | 1 | p.Q254P(LDLR) | | |
| 36; **C** | 5 | p.C46S(LDLR), p.C89Y(LDLR), p.C510R(TMPRSS6), p.C510S(TMPRSS6), p.C119G(C9) | Loss of Cys | high |
| 37; **D** | 7 | p.D90G(LDLR), p.D90N(LDLR), p.D90Y(LDLR), p.D168A(LDLR), p.D168H(LDLR), p.D168N(LDLR), p.D168Y(LDLR) | Loss of Cys | high |
| 38 | 1 | p.G243D(CFI) | | |
| 39; fp | 1 | p.Q92E(LDLR) | | moderate |
| 40 | 3 | p.A50S(LDLR), p.A50T(LDLR), p.R300G(LDLR) | | |
| 41; **D** | 6 | p.D172N(LDLR), p.D221G(LDLR), p.D221N(LDLR), p.D221Y(LDLR), p.D301A(LDLR), p.D301G(LDLR) | Loss of Asp | high |
| 42; **C** | 9 | p.C95G(LDLR), p.C134F(LDLR), p.C134W(LDLR), p.C173W(LDLR), p.C222Y(LDLR), p.C261F(LDLR), p.C302W(LDLR), p.C302Y(LDLR), p.C1361G(LRP5) | Loss of Cys | high |
| 43 | 0 | | | |

| | | | | |
|---|---|---|---|---|
| 44; **D** | 4 | p.D175N(LDLR), p.D175Y(LDLR), p.D224V(LDLR), p.D137N(LRP4) | Loss of Asp | high |
| 45 | 1 | p.G287R(CFI) | | |
| 46; **S** | 3 | p.S56P(LDLR), p.S177L(LDLR), p.S306L(LDLR) | Loss of Ser | moderate |
| 47; **D** | 5 | p.D227E(LDLR), p.D266E(LDLR), p.D521G(TMPRSS6), p.D521N(TMPRSS6), p.D103G(TMPRSS3) | Loss of Asp | high |
| 48; **E** | 6 | p.E101K(LDLR), p.E140K(LDLR), p.E228K(LDLR), p.E228Q(LDLR), p.E522K(TMPRSS6), p.E1367K(LRP5) | Loss of Glu | high |
| 49 | 0 | | | |
| 50 | 0 | | | |
| 51 | 0 | | | |
| 52 | 0 | | | |
| 53 | 0 | | | |
| 54 | 0 | | | |
| 55; **C** | 5 | p.C143R(LDLR), p.C184W(LDLR), p.C184Y(LDLR), p.C231G(LDLR), p.C313Y(LDLR) | Loss of Cys | high |
| 56 | 1 | p.G314R(LDLR) | | |
| | **76** | | | |

### 5 The 3Fn-cassette (residues 1551-2121)

#### 5.a Sequence details

Although the region containing the six 3Fn-domains corresponds to almost a third of the entire SORLA extracellular part (the ectodomain), surprisingly little is known about its function. Many mutations observed in AD patients locate within the 3Fn-domains of SORLA, which suggests that this receptor region represents an important structural and/or functional aspect of SORLA [112,145,146]. In other proteins, 3Fn-domains are commonly involved in ligand binding, but for the SORLA protein, no interaction partner has yet been identified to bind to this region [69]. It can therefore be speculated that this region is important for the structural integrity of SORLA, and may be involved in bending of the full modular receptor, thereby arranging binding surfaces in space. Interaction "partners" may therefore rather be another SORLA molecule to form SORLA-dimers [24], or other domains within the same SORLA protein instead of foreign ligands.

3Fn-domains were originally identified in the modular protein Fibronectin, hence the name of the domain [147]. Fibronectin, which includes 15 copies of the 3Fn-domains, plays myriad fundamental biological roles such as adhesion, cell migration, and hemostasis, and similar multifunctionality is also true for many other proteins containing 3Fn-domains. This domain type is present in a high number of animal and bacterial protein families including extracellular matrix proteins, cell surface receptors, kinases and phosphatases, muscle proteins, etc. [148]. Accordingly, in contrast to the VPS10p-domain and the YWTD/EGF- and CR-domains that are representative of two distinct receptor families, a similar clear affiliation for 3Fn-domain containing proteins is not possible. More than 2,100 domains are listed in PFAM as being 3Fn-domains [148]. The structure of the second SORLA 3Fn-domain is deposited in the protein databank (2DM4.pdb), but not yet described in any publication.

*Binding motif for SORLA 3Fn-domains with interacting partners not yet clear*

3Fn-domains may occur as single repeats in proteins, but they are more frequently clustered as 2-6 adjacent domains [148]. In membrane anchored receptor proteins, including SORLA, the clustered 3Fn-domains are most often located directly proximal to the plasma membrane, where they may engage in contact with other proteins. While ligand interactions are characterized by RGD- or GSWGS-motifs in some 3Fn-domain-containing receptors [149-153], a unifying motif for ligand binding has not (yet) been identified for 3Fn-domains in general nor for the six SORLA domains in particular.

*Structure and important amino acids*

A typical 3Fn-domain structure has an ellipsoid shape with approximate dimensions 38 Å x 20 Å x 25 Å [154]. The incoming and outgoing amino acid sequence ends at opposite sides of the folded domain (**Supplemental Figure S5**), which is in agreement with one of its main functions: to act as a spacer for proper positioning of protein structures [154]. This fold is topologically closely related to that of Immunoglobulin (IgG)-like domains: however, the 3Fn-domains lack the conserved disulfide bonds, and strands A and C' are interchanged between sheets relative to the IgG domains [154]. 3Fn-domains are typically composed of a sequence with 90-100 residues, arranged in seven β-strands (named A, B, C, C', E, F, and G) forming two anti-parallel β-sheets (strands: A-B-E and strands: C-C'-F-G, respectively) (**Supplemental Figure S5**). It is remarkable that despite high similarity in tertiary structure, sequence identity across 3Fn-domains is conspicuously low, typically less than 20% between domains in general [155], which complicates alignment of 3Fn-domain sequences. However, the presence of a few highly conserved amino acids enables unambiguous identification of strands B, C, and F (**Supplemental Figure S5**).

**Strand B** is characterized by a tryptophan (position 25) preceded by two hydrophobic residues at positions 21 and 23; **strand C** contains a tyrosine (position 41) followed by two hydrophobic residues at positions 43 and 45, with the latter position very often occupied by an additional tyrosine residue; **strand F** begins with a tyrosine (position 83) followed by three additional hydrophobic residues at positions 85, 87, and 89, with the latter position often being an alanine. As the hydrophobic residues alternate within a β-strand secondary structure, their side chains point towards the same side of their respective strand, such that they form a large hydrophobic domain-core – sometimes described as 'the glue' between the two β-blades (**Supplemental Figure S5**). The four remaining strands do not contain highly conserved residues, complicating their identification based on their primary structure. However, pairs of alternating hydrophobic residues are likely located in these β-strands as well, such that they can contribute to a hydrophobic core. This is often the case for strand A (positions 11 and 13) where the two hydrophobic amino acids are between 5 and 8 amino acids upstream of strand B, and either one or two prolines at the very beginning (positions 6 and 7) of the 3Fn-domain sequence (**Supplemental Information 5d**). The other three strands (C', E, and G) have very little sequence conservation and can't be identified based on their amino acid sequence analysis, and their location is better identified needs by secondary structure prediction tools.

Both the first and the sixth 3Fn-domain of SORLA include two cysteine residues, suggesting that these two domains contain an intradomain disulfide bond. This is supported by a model showing how their side chains, which we predict are in close proximity, can facilitate intradomain disulfide bond formation in the folded conformation (**Fig. 3**). A single, likely unpaired, cysteine is located in strand B of the fourth 3Fn-domain, but predicted to point its side chain into a hydrophobic core and not be surface exposed.

*Bottom Loops – including The Tyrosine Corner*

Following the conserved folding topology, the loop regions between strands can be grouped as part of the "*top*" or "*bottom*" of a 3Fn-domain (**Supplemental Figure S5**). The BC-, C'E- and FG-loops combined with the N-terminal incoming sequence form the *top* of each 3Fn-domain, whereas the AB-, CC'- and EF-loops form the *bottom* of the domain. Some of these loops also contain conserved residues informative for sequence alignment. Most prominently, in the loop connecting strands E and F (**EF-loop**) that crosses from one sheet to the other, a leucine (position 77) is located six residues upstream of the tyrosine (position 83) in the beginning of strand F. In many 3Fn-domains, including two of the SORLA domains, this loop also contains a conserved proline (position 79) frequently accompanied by a glycine (position 80) (**Supplemental Figure S5**). This structural motif is known as "***the tyrosine corner***", which contributes strongly to the stability of 3Fn-domains: the side chain of Leu-77 packs next to the Tyr-83 ring [156]. Moreover, the -OH group of the Tyr-83 engages in H-bonding with the backbone of the residue five residues upstream (i.e. position 78), naming the 3Fn-domain as the $\Delta5$ subtype of tyrosine corners [157]. While all other loops can elongate without significant loss of conformational stability, the length of the EF-loop is critical to maintain a stable domain fold [158].

*Top Loops – antigen binding homology and N-glycosylations*

Among the loops at the top, a few conserved positions can be noticed: the **BC-loop** often begins with at least one proline at positions 27 and/or 28 as well as a glycine at position 36, and the **FG-loop** preferentially contains a glycine at position 94 – often together with another glycine at position 96 (**Supplemental Information 5d**). This is not evident from the six SORLA sequences due to the low conservation at these positions, but is clearer when looking at our alignment across multiple Fn3-domain sequences (**Supplemental Information 5d**).

Within – or close to – each of the six **C'E-loops** of the SORLA 3Fn-domain is an NXT-motif, each of which represents a potential glycosylation acceptor site [159] (**Fig. 2g**). From our alignment of almost 300 3Fn-domain sequences, we noticed that an *N*-glycan acceptor site is frequently present in the C'E-loop. The function of these glycans probably varies for individual 3Fn-domains. A 3Fn-domain in the interleukin 21 receptor (IL21R) carries a large glycan at the C'E-loop, and this modification was first shown to be essential to keep the relative orientation between two neighboring domains in a fixed position, allowing interactions with residues from the adjacent 3Fn-domain [160]. Second, it was shown that this glycan is essential for the successful transport of IL21R to the cell surface [161]. We recently found that the composition of *N*-glycans in SORLA regulates the proteolytic cleavage by sheddase (TACE) of the membrane-bound SORLA protein at the cell surface, and thus shedding of the soluble SORLA ectodomain [162]. It is tempting to speculate that the regulatory glycan(s) locate in the proximity of the TACE cleavage site, i.e. somewhere in the 3Fn-domain region, and that the glycan(s) induce steric/conformational changes of SORLA, allowing the sheddase to access the cleavage site located just C-terminal to the sixth 3Fn-domain [162].

For IgG-domains, the *top* loops correspond to antigen binding regions (**Supplemental Figure S5**). Possibly, the top loops of the SORLA 3Fn-domains may bind extrinsic ligands. However, there are still many unclarities: many 3Fn-domains occur in tandem arrays of varying lengths, and the structure-function relationship of the entire region is highly dependent on the relative orientations between domain pairs, often measured as "*tilt*" and "*twist*" angles between domains [163] (see **Supplemental Information 12**). These measures are in part determined by the "linkers" between the domains but certainly also by the amino acid composition located in the loop regions.

*Trp-ladders*

Many 3Fn-domains share a motif that resembles a ladder with steps provided by alternating aromatic and charged amino acid side chains [164]. Since the aromatic "steps" are often tryptophan residues, this ladder is termed the "tryptophan-ladder". Generally, such ladders contain six or more "steps", but they may also be shorter. Upon inspection of the SORLA 3Fn-domain sequences, the presence of two short Trp-ladders was observed in the first and second domains most distal to the membrane, with steps made by (3Fn1: R1593, W1600, K1626, and H1636 and 3Fn2: E1690, W1698, R1722, and W1734). The function of these motifs is not completely understood. However, for some proteins it is suggested that they interact directly with sulphated glycoconjugates [165]. It has been demonstrated that some *N*-glycans in the VPS10p-domain of SORLA are terminally sulphated [166], raising the hypothesis that they may be intrinsic targets of the Trp-ladders in 3Fn-region, possibly regulating SORLA-dimer formation and/or self-binding of bent-conformations of SORLA (see **Supplemental Information 12**).

Andersen
Supplemental Figure S5 - 3Fn-domain

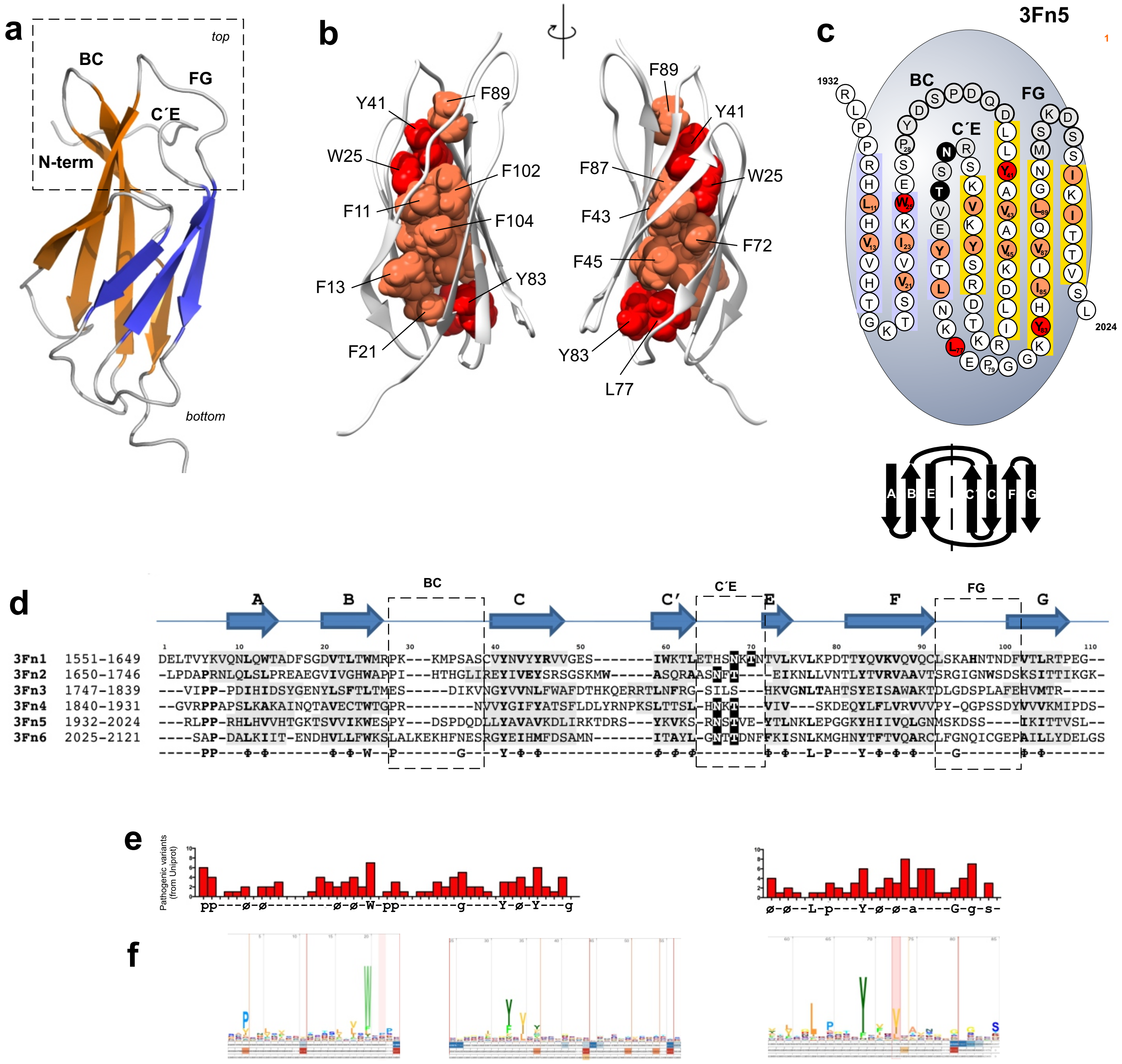

#### 5.b Supplemental Figure S5. 3Fn-domains

**a.** Structure of the second SORLA 3Fn-domain (PDB: 2DM4), showing the two-bladed sandwich conformation representative for 3Fn-domain structures in general. The BC-, C'E- and FG-loops together with the N-terminal residues (boxed) correspond to the antigen binding part of the similarly folded IgG-domains.
**b.** Structure of the 3Fn-domain showing how the alternating hydrophobic residues contribute to a compact core of the domain (orange colored side chains – identified in *panel c*), forming hydrophobic interactions between their side chains that keep the two sheets tightly together. The side chain of the four conserved residues at positions 25 (Trp), 41 (Tyr), 77 (Leu), and 83 (Tyr) are colored red. The two structures represent a 180° turn seeing into the sandwich from opposite sites.
**c**. Residues 1932-2024 of the fifth 3Fn-domain presented according to the characteristic anti-parallel strand topology A-B-E-C'-C-F-G. The two β-sheets are indicated by purple and yellow background for their strands, respectively. The alternating residues that contribute to the hydrophobic domain interior are indicated by orange circles, amino acids that are part of the BC-, C'E-, and FG-loops are on grey background, and the four most conserved residues $W^{25}$, $Y^{41}$, $L^{77}$, and $Y^{83}$ are presented on red circles. Sheet topology is indicated below.
**d**. The SORLA sequence of residues 1551-2121 represents six 3Fn-domains. Alignment of the 3Fn-domain sequences was done according to the β-strand secondary structure diagram shown in panel **c**. The limited number of positions with conserved amino acids and the identified alternating hydrophobic residues (Φ) are presented below the alignment. Residues in top loop regions are included in the broken line boxes. The alignment was specifically designed to allow the conserved consensus motif for *N*-glycosylation (NXT, black background) to be located in the C'E-loop when possible.

**e.** Histograms showing the number of pathogenic variants that occur for each 3Fn-domain position as listed in **Supplemental Information 5f**.

**f.** Logo representation of the domain sequence conservation: the larger the letter the higher the conservation across 3Fn-domain sequences.

#### 5.c *SORL1* variants in 3Fn-domains

*Trp at position 25 (strand B):*

The 3Fn-domain is vulnerable for mutation of the Trp at position 25, for which Uniprot lists seven disease-associated variants: p.W2744C$^{USH2A}$ (segregate with USH2A in a family [167]), p.W3521R$^{USH2A}$ (in patients with USH2A [168]), p.W1036L$^{L1CAM}$ (in patients with HSAS, and functional test of the mutant protein show defective cellular transport [169]), p.W1925R$^{FN1}$ (causal of GFND2 [170]), p.W571R$^{ANOS1}$ (in patients with Kallmann Syndrom/HH1 [171]), p.W792R$^{MYBPC3}$ (in patients with CM44 [172]), and p.W68R$^{GHR}$ (in patients with LARS [173]). In line with the the orthologue data, a report described identification of an AD patient carrying p.W1862C (position 25 in the fourth 3Fn-domain) [174], which suggests that variants at this position in SORLA domains may associate with an increased risk for AD.

*Tyr at position 83 (Tyr-corner):*

The tyrosine at position 83 at the beginning of strand F is conserved across all 3Fn-domains including those in SORLA (**Fig. 3**). Our DMDM analysis found that five of the six other proteins for which Uniprot lists disease-associated substitutions for this position, each time the Tyr was replaced by a Cys: in Tie2, p.Y611C$^{TEK}$ (reduced response to ligand and decreased ligand-induced phosphorylation GLC3E [175]), in the insulin receptor, p.Y818C$^{IR}$ (abolishes post-translational processing, LEPRC [176,177]), in Fibronectin causal for GFND2 p.Y973C$^{FN1}$ (GFND2 [170]), in L1CAM, p.Y784C$^{L1CAM}$ (HSAS [178]) and in the growth hormone receptor, p.Y226C$^{GHR}$ (causal of LARS [179]) (**Supplemental Information 5e**).

It has previously been reported that variant p.Y1816C that locate to the third 3Fn-domain of SORLA was found in AD patients and but not controls [112]. Interestingly, also in the case of SORLA, the tyrosine is substituted by a cysteine, such that a possible pathogenic effect may either be due to loss of the tyrosine or due to introduction of the cysteine, or both. We note that all the other proteins that harbor this type of variant form active dimers by their 3Fn-region, which suggests that such mutations, including p.Y1816C in SORLA, may act via impaired dimerization.

*Position 88 (strand F):*

For position 88, located between the alternating hydrophobic residues in strand F, there is no sequence conservation (**Fig. 2g, Supplemental Information 5d**). However, the side chain of the residue at this position will expose towards the domain surface. Intriguingly, Uniprot lists as many as 9 disease-associated variants at this position (**Supplemental Information 5e**), of which 8 correspond to arginine replacements: p.R926W$^{IR}$ (mutated IR with markedly impaired insulin binding and impaired post-translational processing in patients with LEPRCH [177,180]), p.R312P$^{CRLF1}$ (CISS1 [181]), p.R224W$^{IL2RG}$ (XSCID [182]), p.R201L$^{IL21R}$ (mutant receptor with defective trafficking, misfolding, and impaired processing IMD56 [160,161]), p.R213W$^{IL12RB1}$ (IMD30 [183]), p.R114C$^{IFNGR2}$ (misfolding and abnormal glycosylation, mistrafficking, reduced response to INFG, IMD28 [184,185]), p.R257L$^{MPL}$ (CAMT [186]), and p.R308C$^{CSF3R}$ (decreased localization to plasma membrane and decreased receptor signaling, SCN7 [187]). Detailed analysis of the variant in IL21R revealed how this arginine side chain is required for interaction with a glycan from a neighboring 3Fn-domain and the relative domain-domain orientation [160]. The SORLA sequence includes only an arginine at position 88 in the fourth 3Fn-domain, and no variants is yet identified for this residue.

*Positions 94 and 96 (glycines in FG-loop):*

From Uniprot we identified seven disease-associated variants at position 96 of the 3Fn-domain according to our DMDM analysis (**Supplemental Information 5e**). There is a slight preference for Gly at this position according to the 3Fn-domain consensus sequence, but it is not nearly as conserved as Gly at position 94. (**Supplemental Information 5d**). However, we noticed that in five of the seven identified disease variants for position 96, a Gly was substituted. Furthermore, four of these variants were observed when the Gly two residues upstream (at position 94) in the sequence was also present: p.G698R$^{L1CAM}$ (causal for hydrocephalus HSAS/MASA and mutation is inherited in a mendelian fashion segregating with disease [188,189]), p.G805E$^{DCC}$ (causal of isolated agenesis of the corpus callosum (MRMV1) in a family, and mutation disturb nestrin-1 binding to the FG-loop of the DCC 3Fn-domain [190]), p.G2757V$^{SPEG}$ (causal of Centronuclear Myopathy with Dilated Cardiomyopathy (CNM5) in a small family [191]), and p.G516R$^{EPHB4}$ (mutation segregate with the Capillary Malformation-Arteriovenous Malformation (CMAVM2)-phenotype in three small families [192]). We speculate that co-occurrence of both Gly residues could have functional relevance relating to their localization in the FG-loop region preferring to accommodate residues with small side chains. In SORLA, only the second 3Fn-domain contains this double Gly at positions 94 and 96 (amino acids 1730 and 1732) (**Fig. 3**). Very interestingly variant p.G1732A corresponding to position 96 is reported to segregate with AD in a Swedish family [65], in support of this variant being pathogenic.

#### 5.d 3Fn-domain alignment

Mapping of naturally occurring variants found in human 3Fn-domain containing proteins, and being listed as associated with pathology at [www.uniprot.org](http://www.uniprot.org). The alignment follows the SORLA alignment shown on top, and pathogenic variants are highlighted on a red background.

**SORLA:**

```
   DELTVYKVQNLQWTA-DFSGDVTLTWMRPK---KMPSASCVYNVYYRVVGES-------IWKTLETHSNKTN----------------TVLKVLKPDTTYQVKVQVQCLSKAHNTNDFVTLRTPEG--
   -LPDAPRNLQLSLPR-EAEGVIVGHWAPPI---HTHGLIREYIVEYSRSGSKMW-----ASQRAASNFT--------------------EIKNLLVNTLYTVRVAAVTSRGIGNWSDSKSITTIKGK-
   ---VIPP-PDIHIDS-YGENYLSFTLTMES-----DIKVNGYVVNLFWAFDTHKQERRTLNFRG-SILS-------------------HKVGNLTAHTSYEISAWAKTDLGDSPLAFEHVMTR-----
   --GVRPPAPSLKAKA-INQTAVECTWTGPR--------NVVYGIFYATSFLDLYRNPKSLTTSL-HNKT-------------------VIV---SKDEQYLFLVRVVVPY-QGPSSDYVVVKMIPDS-
   ---RLPP-RHLHVVH-TGKTSVVIKWESPY---DSPDQDLLYAVAVKDLIRKTDRS---YKVKS-RNSTVE-----------------YTLNKLEPGGKYHIIVQLGNMSKDSS----IKITTVSL-
   ----SAP-DALKIIT--ENDHVLLFWKSLALKEKHFNESRGYEIHMFDSAMN-------ITAYL-GNTTDNF----------------FKISNLKMGHNYTFTVQARCLFGNQICGEPAILLYDELGS
        pp   Φ Φ        Φ Φ W pp       g    Y Φ Φ   g         Φ Φ Φ                        Φ Φ  L p   Y Φ Φ a    g       Φ Φ
                       sø ø W PP       g ø  Y ø ø                                            ø gL p t Y ø ø a    G g s
```

**Usherin:**

```
   PFQQPPP--RGQVQS---SSAINLSWSPPD---SPNAHWLTYSLLRDGFEIYTTEDQYPYSIQY------------------------FLDTDLLPYTKYSYYIETTNVHGSTRSVAVTYKTKP
GVPEGNLTLS--YIIPIG---SDSVTLTWTTLS---NQSGPIEKYILSCAPLAGGQPCVSYEGHETS------------------------ATIWNLVPFAKYDFSVQACTSGGCLHSLPITVTTAQAPPQ
     RLSPP--KMQKIS---STELHVEWSPPA---ELNGIIIRYELYMRRLRSTKETTSEESRVFQSSGWLSPHSFVESANENALKPPQTMTTITGLEPYTKYEFRVLAVNMAGSVSSAWVSERTGESAPVF
      MIPP-SVFPLSS----YSLNISWEKPA-DNVTRGKVVGYDINMLSEQSPQQSIPMAFSQLLHTAKSQELS---------------YTVEGLKPYRIYEFTITLCNSVGCVTSASGAGQTLAAAPA
      RGAVVNLASVSSGAVRVNLDGCLSTDS-AVNCRGNDSILVYQGKEQS--------------------------------------VYEGGLQPFTEYLYRVIASHEGGSVYSDWSRGRTTGAAPQ
    SVPTPS--RVRSLN---GYSIEVTWDEPV----VRGVIEKYILKAYSEDSTRPPRMPSASAEFVNTSNLT-----------------GILTGLLPFKNYAVTLTACTLAGCTESSHALNISTPQE
  APQEVQPP--VAKSL----PSSLLLSWNPPK---KANGIITQYCLYMDGRLIYSGSEEN------------------------------YIVTDLAVFTPHQFLLSACTHVGCTNSSWVLLYTAQLPPE
     HVDSP--VLTVLD---SRTIHIQWKQPR---KISGILERYVLYMSNHTHDFTIWSVIYNSTELFQD--------------------HMLQYVLPGNKYLIKLGACTGGGCTVSEASEALTDED
  IPEGVPAP--KAHSYS---PDSFNVSWTEPE---YPNGVITSYGLYLDGILIHNSSELSYRAYGFAPWSLHSFRVQACT----------AKGCALGPLVENRTLEAPPE
       GTVNVFVKTQG---SRKAHVRWEAPF---RPNGLLTHSVLFTGIFYVDPVGNNYTLLNVTKVMYSGEETNLW------------VLIDGLVPFTNYTVQVNISNSQGSLITDPITIAMPPG
  APDGVLPP--RLSSAT---PTSLQVVWSTPA-RNNAPG-SPRYQLQMRSGDSTHGFLELFSNPSASLS---------------------YEVSDLQPYTEYMFRLVASNGFGSAHSSWIPFMTAED
  KPGPVVPP--ILLDVK---SRMMLVTWQHPR---KSNGVITHYNIYLHGRLYLRTPGNVTN----------------------------CTVMHLHPYTAYKFQVEACTSKGCSLSPESQTVWTLPG
  APEGIPSP--ELFSDT---PTSVIISWQPPT---HPNGLVENFTIERRVKGKEEVTTLVTLPRSHSMRFI------------------DKTSALSPWTKYEYRVLMSTLHGGTNSSAWVEVTTRPS
  RPAGVQPP--VVTVLE---PDAVQVTWKPPL---IQNGDILSYEIHMPDPHITLTNVTSAVLS--------------------------QKVTHLIPFTNYSVTIVACSGGNGYLGGCTESLPTYVTTHPT
   VPQNVGP-LSVIPLS---ESYVVISWQPPS---KPNGPNLRYELLRRKIQQPLASNPPEDLNRWHNIYSGTQWL--------------YEDKGLSRFTTYEYMLFVHNSVGFTPSREVTVTTLAG
  LPERGANL--TASVLN---HTAIDVRWAKPT-VQDLQGEVEYYTLFWSSATSNDSLKILPDVNS-------------------------HVIGHLKPNTEYWIFISVFNGVHSINSAGLHATTCDG
  EPQGMLPP--EVVIIN---STAVRVIWTSPS---NPNGVVTEYSIYVNNKLYKTGMNVPGS----------------------------FILRDLSPFTIYDIQVEVCTIYACVKSNGTQITTVEDTPSD
      IPTP--TIRGIT---SRSLQIDWVSPR---KPNGIILGYDLLWKTWYPCAKTQKLVQDQSDELCKAVRC----------------QKPESICGHICYSSEAKVCCNGVLYNPKPGHRCCEE
     CPASMEATEHCGRCDFNFTSHICTVIRG-SHNSTGKASIEEMCSSAEETIHTGSVNTYS---------------------------YTDVNLKPYMTYEYRISAWNSYGRGLSKAVRARTKED
  VPQGVSPP--TWTKID-NLEDTIVLNWRKPI---QSNGPIIYYILLRNGIERFRGTSLSF----------------------------SDKEGIQPFQEYSYQLKACTVAGCATSSKVVAATTQG
  VPESILPP--SITALS---AVALHLSWSVPE---KSNGVIKEYQIRQVGKGLIHTDTTDRRQ---------------------------HTVTGLQPYTNYSFTLTACTSAGCTSSEPFLGQTLQAAPE
    GVWVTP--RHIIIN---STTVELYWSLPE---KPNGLVSQYQLSRNGNLLFLGGSEEQN----------------------------FTDKNLEPNSRYTYKLEVKTGGGSSASDDYIVQTPMS
  TPEEIYPP-YNITVIG---PYSIFVAWIPPG--ILIPEIPVEYNVLLNDGSVTPLAFSVGHHQS-------------------------TLLENLTPFTQYEIRIQACQNGSCGVSSRMFVKTPEA
  APMDLNSP--VLKALG---SACIEIKWMPPE---KPNGIIINYFIYRRPAGIEEESVLFVWSEGALEFM--------------------DEGDTLRPFTLYEYRVRACNSKGSVESLWSLTQTLEA
  PPQDFPAP--WAQATS---AHSVLLNWTKPE---SPNGIISHYRVVYQERPDDPTFNSPTVHAFTVKGTSHQ----------------AHLYGLEPFTTYRIGVVAANHAGEILSPWTLIQTLES
      SPSGLRNFIVEQKENGRALLLQWSEPM---RTNGVIKTYNIFSDGFLEYSGLNRQ------------------------------FLFRRLDPFTLYTLTLEACTRAGCAHSAPQPLWTDEA
```

```
PPDSQLAP--TVHSVK---STSVELSWSEPV---NPNGKIIRYEVIRRCFEGKAWGNQTIQADEKIVFTEYNTERNTFM-----------YNDTGLQPWTQCEYKIYTWNSAGHTCSSWNVVRTLQA
PPEGLSPP--VISYVS-MNPQKLLISWIPPE---QSNGIIQSYRLQRNEMLYPFSFDPVTFN----------------------------YTDEELLPFSTYSYALQACTSGGCSTSKPTSITTLEAAPSE
VSPP--DLWAVS---ATQMNVCWSPPT---VQNGKITKYLVRYDNKESLAGQGLC-------------------------------LLVSHLQPYSQYNFSLVACTNGGCTASVSKSAWTMEALPE
NMDSP--TLQVTG---SESIEITWKPPR---NPNGQIRSYELRRDGTIVYTGLETR-------------------------------YRDFTLTPGVEYSYTVTASNSQGGILSPLVKDRTSPS
APSGMEPP--KLQARG---PQEILVNWDPPV---RTNGDIINYTLFIRELFERETKIIHINTTHNSFGMQS-------------------YIVNQLKPFHRYEIRIQACTTLGCASSDWTFIQTPEIAPLM
QPPPHLEVQMAPGGFQPTVSLLWTGPL---QPNGKVLYYELYRRQIATQPRKSNPVLIYNGSSTS---------------------FIDSELLPFTEYEYQVWAVNSAGKAPSSWTWCRTGPA
PPEGLRAP--TFHVIS---STQAVVNISAPG---KPNGIVSLYRLFSSSAHGAETVLSEGMATQ--------------------------QTLHGLQAFTNYSIGVEACTCFNCCSKGPTAELRTHPA
PPSGLSSP--QIGTLA---SRTASFRWSPPM---FPNGVIHSYELQFHVACPPDSALPCTPSQIETKYTGLGQK----------------ASLGGLQPYTTTYKLRVVAHNEVGSTASEWISFTTQKELP

L1CAM:
SPGPVPRLVLSDLHLLTQSQVRVSWSPA---EDHNAPIEKYDIEFEDKEMAPEKWYSLGKVPGNQT---------------------STTLKLSPYVHYTFRVTAINKYGPGEPSPVSETVVTPEA
APEKNP-VDVKGEG-NETTNMVITWKPL-RWMDWNAPQVQYRVQWRPQGTRGPWQEQIVSDPF------------------------LVVSNTSTFVPYEIKVQAVNSQGKGPEPQVTIGYSGED
YPQAIP-ELEGIEI-LNSSAVLVKWRPV-DLAQVKGHLRGYNVTYWREGSQRKHSKRHIHKDHVVVPANTTS---------------VILSGLRPYSSYHLEVQAFNGRGSGPASEFTFSTPEG
VPGHP-EALHLEC-QSNTSLLLRWQPP---LSHNGVLTGYVLSYHPLDEGGKGQLSFNLRDPELRT---------------------HNLTDLSPHLRYRFQLQATTKEGPGEAIVREGGTMALS
GISDFGNISATA--GENYSVVSWVPK-----EGQCNFRFHILFKALGEEKGGASLSPQYVSYNQSS-------------------YTQWDLQPDTDYEIHLFKERMFRHQMAVKTNGTGRVRLPPA
INSR:
CENELLKFSYIR-TSFDKILLRWEPYW--PPDFRDLLGFMLFYKEAPYQNVTEFDGQDACGSNSWTVVDIDPPLRSNDPKSQNHPGWLMRGLKPWTQYAIFVKTLVTFSDERRTYGAKSDIIYVQTDA
NPSVPLDPISVSN--SSSQIILKWKPPS---DPNGNITHYLVFWERQAEDSELFELDYCLΛNKES----------------------
LVISGLRHFTGYRIELQACNQDTPEERCSVAAYVSARTMPEAKADD
IVGPVTHEIFEN----NVVHLMWQEPK---EPNGLIVLYEVSYRRYGDEELHLCVSRKHFALERG--------------------CRLRGLSPG-NYSVRIRATSLAGNGSWTEPTYFYVTDY
IGF1R:
PERKRRDVMQVANTTMSSRSRNTTAADTYNITDPEELETEYPFFESRVDNKER-----------------------------------TVISNLRPFTLYRIDIHSCNHEAEKLGCSASNFVFARTMPA
IP-GPVTWEP-RPENSIFLKWPEPE---NPNGLILMYEIKYGSQVEDQRECVSRQEYRKYGG----------------------AKLNRLNPG-NYTARIQATSLSGNGSWTDPVFFYVQAK
Fibronectin:
GPVEVFITETPSQPNSHPIQWNAPQ-----PSHISKYILRWRPKNSVGRWKEATIPGHLNS-----------------------YTIKGLKPGVVYEGQLISIQQYGHQEVTRFDFTTTST
TSESVTEIT---ASSFVVSWVSAS------DTVSGFRVEYELSEEGDEPQYLDLPSTATS-----------------------VNIPDLLPGRKYIVNVYQISEDGEQSLILSTSQTTAPD
APP--DTTVDQ-VDDTSIVVRWSRPQ------APITGYRIVYSPSVEGSSTELNLPETANS------------------------VTLSDLQPGVQYNITIYAVEENQESTPVVIQQETTGTPRSDT
VPSP-RDLQFVE-VTDVKVTIMWTPPE------SAVTGYRVDVIPVNLPGEHGQRLPISRNTF-----------------------AEVTGLSPGVTYYFKVFAVSHGRESKPLTAQQTTKLD
AP-TNLQFVN-ETDSTVLVRWTPPR------AQITGYRLTVGLTRRGQPRQYNVGPSVSK------------------------YPLRNLQPASEYTVSLVAIKGNQESPKATGVFTTLQPG
SSIPP-YNTEVTE----TTIVITWTPAP--------RIGFKLGVRPSQGGEAPREVTSDSGS-------------------------IVVSGLTPGVEYVYTIQVLRDGQERDAPIVNKVVTPLS
PP-TNLHLEANPDTGVLTVSWERST-----TPDITGYRITTTPTNGQQGNSLEEVVHADQSS----------------------CTFDNLSPGLEYNVSVYTVKDDKESVPISDTIIPEVPQL
TDLSFVD-ITDSSIGLRWTPLN-----SSTIIGYRITVVAAGEGIPIFEDFVDSSVGY----------------------YTVTGLEPGIDYDISVITLINGGESAPTTLTQQTAV
PPP-TDLRFTN-IGPDTMRVTWAPPP-----SIDLTNFLVRYSPVKNEEDVAELSISPSDNA-----------------------VVLTNLLPGTEYVVSVSSVYEQHESTPLRGRQKTGLD
SP-TGIDFSD-ITANSFTVHWIAPR------ATITGYRIRHHPEHFSGRPREDRVPHSRNS-----------------------ITLTNLTPGTEYVVSIVALNGREESPLLIGQQSTVSD
VP-RDLEVVA-ATPTSLLISWDAPA------VTVRYYRITYGETGGNSPVQEFTVPGSKST----------------------ATISGLKPGVDYTITVYAVTGRGDSPASSKPISINYRTEID
KP-SQMQVTD-VQDNSISVKWLPSS------SPVTGYRVTTTPKNGPGPTKTKTAGPDQTE-----------------------MTIEGLQPTVEYVVSVYAQNPSGESQPLVQTAVTNID
RP-KGLAFTD-VDVDSIKIAWESPQ------GQVSRYRVTYSSPEDGIHELFPAPDGEEDT-----------------------AELQGLRPGSEYTVSVVALHDDMESQPLIGTQSTAIP
AP-TDLKFTQ-VTPTSLSAQWTPPN------VQLTGYRVRVTPKEKTGPMKEINLAPDSSS-----------------------VVVSGLMVATKYEVSVYALKDTLTSRPAQGVVTTLENVS
PP-RRARVTD-ATETTITISWRTKT------ETITGFQVDAVPANGQTPIQRTIKPDVRS------------------------YTITGLQPGTDYKIYLYTLNDNARSSPVVIDASTAID
AP-SNLRFLA-TTPNSLLVSWQPPR------ARITGYIIKYEKPGSPPREVVPRPRPGVTE----------------------ATITGLEPGTEYTIYVIALKNNQKSEPLIGRKKTDELP
GPGLNP--NASTGQ-EALSQTTISWAPFQ-------DTSEYIISCHPVGTDEEPLQFRVPGTSTS----------------------ATLTGLTRGATYNVIVEALKDQQRHKVREEVVTVGNSVNEG
DCC:
APRDVVPVLV--SSRFVRLSWRPPA---EAKGNIQTFTVFFSREGDNRERALNTTQPGSLQ----------------------LTVGNLKPEAMYTFRVVAYNEWGPGESSQPIKVATQPE
LQVPGPVENLQAVS-TSPTSILITWEPPA---YANGPVQGYRLFCTEVSTGKEQNIEVDGLS--------------------------YKLEGLKKFTEYSLRFLAYNRYGPGVSTDDITVVTLSD
VPSAPP-QNVSLEV-VNSRSIKVSWLPPP-----SGTQNGFITGYKIRHRKTTRRGEMETLEPNNLW----------------------YLFTGLEKGSQYSFQVSAMTVNGTGPPSNWYTAETPEN
DLDESQVPDQP-SSLHVRP--QTNCIIMSWTPPL---NPNIVVRGYIIGYGVGSPYAETVRVDSKQRY-------------------------YSIERLESSSHYVISLKAFNNAGEGVPLYESATTRSITDPT
```

```
FPTSVPDLSTPMLPP-VGVQAVA-LTHDAVRVSWADNSVPKNQKTSEVRLYTVRWRTSFSASAKYKSEDTTSLS-----------------------YTATGLKPNTMYEFSVMVTKNRRSSTWSMTAHATTYEA
         APTSAPKDLTVITREGKPRAVIVSWQPPL---EANGKITAYILFYTLDKNIPIDDWIMETISGDRLT---------------------HQIMDLNLDTMYYFRIQARNSKGVGPLSDPILFRTLKV
```

**Leptin receptor:**

```
                PP-LGLHMEI-TDDGNLKISWSSPP----LVPFPLQYQVKYSENSTTVIREADKIVSATS------------------------LLVDSILPGSSYEVQVRGKRLDGPGIWSDWSTPRVFTTQDV
                PP-SSVKAEITINIGLLKISWEKPV----FPENNLQFQIRYGLSGKEVQWKMYEVYDAKSKS----------------------VSLPVPDLCAVYAVQVRCKRLDGLGYWSNWSNPAYTVVM
       DIKVPMRGPEFWRIINGDTMKKEKNVTLLWKPLM-KNDSLCSVQRYVINHHTSCNGTWSEDVGNH--------------------------TKFTFLWTEQAHTVTVLAINSIGASVANFNLTFS
          WPMSKVNIVQSLSAYP-LNSSCVIVSWILSP----SDYKLMYFIIEWKNLNEDGEIKWLRISSSVKKY----------------------YIHDHFIPIEKYQFSLYPIFMEGVGKPKIINSFTQDDI
```

**Prolactin receptor:**

```
             PPGKP-EIFKCRS-PNKETFTCWWRPGT----DGGLPTNYSLTYHREGETLMHECPDYITGGPNSCH--------------------FGKQYTSMWRTYIMMVNATNQMGSSFSDELYVDVTYIVQPD
             PPLELAVEVKQPE-DRKPYLWIKWSPPT---LIDLKTGWFTLLYEIRLKPEKAAEWEIHFAGQQTE---------------------FKILSLHPGQKYLVQVRCKPDHGYWSAWSPATFIQIPSD
```

**CRLF1:**

```
             PPEKP-VNISCWS-KNMKDLTCRWTPGA--HGETFLHTNYSLKYKLRWYGQDNTCEEYHTVGPHSC--------------------HIPKDLALFTPYEIWVEATNRLGSARSDVLTLDILDVVTTDP
                PP-DVHVSRVGGLEDQLSVRWVSPP-ALKDFLFQAKYQIRYRVEDSVDWKVVDDVSNQTS-----------------------CRLAGLKPGTVYFVQVRCNPFGIYGSKKAGIWSEWSHPTAA
```

**Integrin beta4:**

```
             APQNP--NAKAAG---SRKIHFNWLPPS------GKPMGYRVKYWIQGDSESEAHLLDSKVPS------------------------VELTNLYPYCDYEMKVCAYGAQGEGPYSSLVSCRTHQE
             VPSEP-GRLAFNV-VSSTVTQLSWAEPA---ETNGEITAYEVCYGLVNDDNRPIGPMKKVLVDNPKNRM-----------------LLIENLRESQPYRYTVKARNGAGWGPEREAIINLATQPK
                        FSALGPTSLRVSWQEPR----CERPLQGYSVEYQLLNGGELHRLNIPNPAQTS----------------------VVVEDLLPNHSYVFRVRAQSQEGWGREREGVITIESQVHPQ
    LPGSAFTLSTPSAP-GPLVFTA-LSPDSLQLSWERPR---RPNGDIVGYLVTCEMAQGGGPATAFRVDGDSPESR--------------------
LTVPGLSENVPYKFKVQARTTEGFGPEREGIITIESQDGGPFP
```

**ROBO3:**

```
    PDPPTEPSSPPGAP-SQPVVTE-ITKNSITLTWKPNP---QTGAAVTSYVIEAFSPAAGNTWRTVADGVQLET---------------------HTVSGLQPNTIYLFLVRAVGAWGLSEPSPVSEPVRTQDS
          VAVRLQEP---IVLGP----RTLQVSWTVDG----PVQLVQGFRVSWRVAGPEGGSWTMLDLQSPSQQS--------------------TVLRGLPPGTQIQIKVQAQGQEGLGAESLSVTRSIPE
                PPQGVAVALGGDGNSSITVSWEPPL-PSQQNGVITEYQIWCLGNESRFHLNRSAAGWARS-----------------------AMLRGLVPGLLYRTLVAAATSAGVGVPSAPVLVQLPSPPDL
```

**ROBO4:**

```
             TLLNP-DPAEGPK--PRPAVWLSWKVSG----PAAPAQSYTALFRTQTAPGGQGAPWAEELLAGWQS-------------------AELGGLHWGQDYEFKVRPSSGRARGPDSNVLLLRLPEK
            VPSAPP-QEVTLKP--GNGTVFVSWVPPP-AENHNGIIRGYQVWSLGNTSLPPANWTVVGEQT----------------------QLEIATHMPGSYCVQVAAVTGAGAGEPSRPVCLLLEQAMERA
```

**Tenascin:**

```
                PP-KDLVVTE-VTEETVNLAWDNEM-------RVTEYLVVYTPTHEGGLEMQFRVPGDQTS---------------------TIIQELEPGVEYFIRVFAILENKKSIPVSARVATYLPAP
                   EGLKFKS-IKETSVEVEWDPLD------IAFETWEIIFRNMNKEDEGEITKSLRRPETS----------------------YRQTGLAPGQEYEISLHIVKNNTRGPGLKRVTTTRLD
                AP-SQIEVKD-VTDTTALITWFKPL------AEIDGIELTYGIKDVPGDRTTIDLTEDENQ---------------------YSIGNLKPDTEYEVSLISRRGDMSSNPAKETFTTGLD
                AP-RNLRRVS-QTDNSITLEWRNGK------AAIDSYRIKYAPISGGDHAEVDVPKSQQATTK-------------------TTLTGLRPGTEYGIGVSAVKEDKESNPATINAATELDTPKD
                     LQVSE-TAETSLTLLWKTPL------AKFDRYRLNYSLPTGQWVGVQLPRNTTS----------------------YVLRGLEPGQEYNVLLTAEKGRHKSKPARVKASTEQAP
                 ELENLTVTE-VGWDGLRLNWTAAD------QAYEHFIIQVQEANKVEAARNLTVPGSLRA---------------------VDIPGLKAATPYTVSIYGVIQGYRTPVLSAEASTGE
               TPNLGEVVVAE-VGWDALKLNWTAPE------GAYEYFFIQVQEADTVEAAQNLTVPGGLRS---------------------TDLPGLKAATHYTITIRGVTQDFSTTPLSVEVLTEE
               VPDMGNLTVTE-VSWDALRLNWTTPD------GTYDQFTIQVQEADQVEEAHNLTVPGSLRS---------------------MEIPGLRAGTPYTVTLHGEVRGHSTRPLAVEVVTEDLPQL
                   GDLAVSE-VGWDGLRLNWTAAD------NAYEHFVIQVQEVNKVEAAQNLTLPGSLRA---------------------VDIPGLEAATPYRVSIYGVIRGYRTPVLSAEASTAK
               EPEIGNLNVSD-ITPESFNLSWMATD------GIFETFTIEIIDSNRLLETVEYNISGAERT---------------------AHISGLPPSTDFIVYLSGLAPSIRTKTISATATTEA
               LPLLENLTISD-INPYGFTVSWMASE------NAFDSFLVTVVDSGKLLDPQEFTLSGTQRK---------------------LELRGLITGIGYEVMVSGFTQGHQTKPLRAEIVTEA
               EPEVDNLLVSD-ATPDGFRLSWTADE------GVFDNFVLKIRDTKKQSEPLEITLLAPERT---------------------RDITGLREATEYEIELYGISKGRRSQTVSAIATTAMG
                SP-KEVIFSD-ITENSATVSWRAPT------GAQVESFRITYVPITGGTPSMVTVDGTKTQ---------------------TRLVKLIPGVEYLVSIIAMKGFEESEPVSGSFTTALD
                        N-ITDSEALARWQPAI------GATVDSYVISYTGEKVPEITRTVSGNTVE---------------------YALTDLEPATEYTLRIFAEKGPQKSSTITAKFTTDLD
                        E-VQSETALLTWRPPR------GASVTGYLLVYESVDGTVKEVIVGPDTTS---------------------YSLADLSPSTHYTAKIQALNGPLRSNMIQTIFTTIGLL
```

**TenascinX:**

```
             RPQELGELRVLGRDETGRLRVVWTAQP------GDTFAYFQLRMRVPEGPGAHEEVLPGDVRQ---------------------ALVPPPPPGTPYELSLHGVPPGGKPSDPIIYQGIMDKDEEKP
                 RLGELTVTD-RTSDSLLLRWTVPE------GEFDSFVIQYKDRDGQPQVVPVEGPQRS----------------------AVITSLDPGRKYKFVLYGFVGKKRHGPLVAEAKILPQSD
           PSPGTPPHLGNLWVTD-PTPDSLHLSWTVPE------GQFDTFMVQYRDRDGRPQVVPVEGPERS----------------------FVVSSLDPDHKYRFTLFGIANKKRYGPLTADGTTAPERKE
          RPEFLEQPLLGELTVTG-VTPDSLRLSWTVAQ------GPFDSFMVQYKDAQGQPQAVPVAGDENE----------------------VTVPGLDPDRKYKMNLYGLRGRQRVGPESVVAKTAPQEDVDE
```

```
GTEAPESPEEPLLGELTVTG-SSPDSLSLFWTVPQ------GSFDSFTVQYKDRDGRPRAVRVGGKESE-------------------------VTVGGLEPGHKYKMHLYGLHEGQRVGPVSAVGVTAPQQEET
TESPLEPRLGELTVTD-VTPNSVGLSWTVPE------GQFDSFIVQYKDKDGQPQVVPVAADQRE-------------------------VTVYNLEPERKYKMNMYGLHDGQRMGPLSVVIVTAPLPPAPA
KPPLEPRLGELTVTD-ITPDSVGLSWTVPE------GEFDSFVVQYKDRDGQPQVVPVAADQRE-------------------------VTIPDLEPSRKYKFLLFGIQDGKRRSPVSVEAKTVARGDA
SPGAPPRLGELWVTD-PTPDSLRLSWTVPE------GQFDSFVVQFKDKDGPQVVPVEGHERS--------------------------VTVTPLDAGRKYRFLLYGLLGKKRHGPLTADGTTEARSAMDD
KRPPKPRLGEELQVTT-VTQNSVGLSWTVPE------GQFDSFVVQYKDRDGQPQVVPVEGSLRE-------------------------VSVPGLDPAHRYKLLLYGLHHGKRVGPISAVAITAGREETET
APTPPAPEPHLGELTVEE-ATSHTLHLSWMVTE------GEFDSFEIQYTDRDGQLQMVRIGGDRND-------------------------ITLSGLESDHRYLVTLYGFSDGKHVGPVHVEALTVPEEEKPS
ATPEPPIKPRLGELTVTD-ATPDSLSLSWTVPE------GQFDHFLVQYRNGDGQPKAVRVPGHEEG-------------------------VTISGLEPDHKYKMNLYGFHGGQRMGPVSVVGVTAAEEETPS
APEPAEEPLLGELTVTG-SSPDSLSLSWTVPQ------GRFDSFTVQYKDRDGRPQVVRVGGEESE-------------------------VTVGGLEPGRKYKMHLYGLHEGRRVGPVSAVGVTAPEE
ESPDAPLAKLRLGQMTVRD-ITSDSLSLSWTVPE------GQFDHFLVQFKNGDGQPKAVRVPGHEDG-------------------------VTISGLEPDHKYKMNLYGFHGGQRVGPVSAVGLTAPGKDEEM
EPPTPEPPIKPRLEELTVTD-ATPDSLSLSWTVPE------GQFDHFLVQYKNGDGQPKATRVPGHEDR-------------------------VTISGLEPDNKYKMNLYGFHGGQRVGPVSAIGVTAAEEETPS
APEPPEEPLLGELTVTG-SSPDSLSLSWTVPQ------GRFDSFTVQYKDRDGRPQVVRVGGEESE-------------------------VTVGGLEPGRKYKMHLYGLHEGRRVGPVSTVGVTAPQEDVDE
APGPPEEPLLGELTVTG-SSPDSLSLSWTVPQ------GRFDSFTVQYKDRDGRPQAVRVGGQESK-------------------------VTVRGLEPGRKYKMHLYGLHEGRRLGPVSAVGVTEDEAETTQ
VPTMTPEPPIKPRLGELTMTD-ATPDSLSLSWTVPE------GQFDHFLVQYRNGDGQPKAVRVPGHEDG-------------------------VTISGLEPDHKYKMNLYGFHGGQRVGPISVIGVTAAEEETPS
APEPPEEPLLGELTVTG-SSPDSLSLSWTIPQ------GHFDSFTVQYKDRDGRPQVMRVRGEESE-------------------------VTVGGLEPGRKYKMHLYGLHEGRRVGPVSTVGVTAPEDEAET
PPNKPRLGELTVTD-ATPDSLSLSWMVPE------GQFDHFLVQYRNGDGQPKVVRVPGHEDG-------------------------VTISGLEPDHKYKMNLYGFHGGQRVGPISVIGVTAAEEETPA
APEPPEEPLLGELTVTG-SSPDSLSLSWTIPQ------GRFDSFTVQYKDRDGRPQVVRVRGEESE-------------------------VTVGGLEPGCKYKMHLYGLHEGQRVGPVSAVGVTAPKDEAE
AVPTMTPEPPIKPRLGELTVTD-ATPDSLSLSWMVPE------GQFDHFLVQYRNGDGQPKAVRVPGHEDG-------------------------VTISGLEPDHKYKMNLYGFHGGQRVGPVSAIGVTEEETPSPT
APEAPEEPLLGELTVTG-SSPDSLSLSWTVPQ------GRFDSFTVQYKDRDGQPQVVRVRGEESE-------------------------VTVGGLEPGRKYKMHLYGLHEGQRVGPVSTVGITAPLPTPLP
VEPRLGELAVAA-VTSDSVGLSWTVAQ------GPFDSFLVQYRDAQGQPQAVPVSGDLRA-------------------------VAVSGLDPARKYKFLLFGLQNGKRHGPVPVEARTAPDTKPS
PRLGELTVTD-ATPDSVGLSWTVPE------GEFDSFVVQYKDKDGRLQVVPVAANQRE-------------------------VTVQGLEPSRKYRFLLYGLSGRKRLGPISADSTTAPLEK
ELPPHLGELTVAE-ETSSSLRLSWTVAQ------GPFDSFVVQYRDTDGQPRAVPVAADQRT-------------------------VTVEDLEPGKKYKFLLYGLLGGKRLGPVSALGMTAPEEDTPA
APEPPEEPRLGVLTVTD-TTPDSMRLSWSVAQ------GPFDSFVVQYEDTNGQPQALLVDGDQSK-------------------------ILISGLEPSTPYRFLLYGLHEGKRLGPLSAEGTTGLAPAGQT
SEESRPRLSQLSVTD-VTTSSLRLNWEAPP------GAFDSFLRFGVPSPSTLEPHPRPLLQRELMVPGTRHS----------------AVLRDLRSGTLYSLTLYGLRGPHKADSIQGTARTLSPVLE
SP-RDLQFSE-IRETSAKVNWMPPP------SRADSFKVSYQLADGGEPQSVQVDGQART------------------------QKLQGLIPGARYEVTVVSVRGFEESEPLTGFLTTVPD
GP-TQLRALN-LTEGFAVLHWKPPQ------NPVDTYDVQVTAPGAPPLQAETPGSAVD-------------------------YPLHDLVLHTNYTATVRGLRGPNLTSPASITFTTGLE
AP-RDLEAKE-VTPRTALLTWTEPP------VRPAGYLLSFHTPGGQNQEILLPGGITS-------------------------HQLLGLFPSTSYNARLQAMWGQSLLPPVSTSFTTGGLRIP
```

**SPEG:**

```
KLAPP----EVPQ-TYQDTALVLWKPGD-----SRAPCTYTLERRVDGESVWHPVSSGIPDCYYN----------------------VTHLPVGVTVRFRVAC--ANRAGQGPFSNSSEKVFVRGTQDS
```

**COL6A3:**

```
MSREVQVFE-ITENSAKLHWERAE-----PPGPYFYDLTVTSAHDQSLVLKQNLTVTD-------------------------RVIGGLLAGQTYHVAVVCYLRSQVRATYHGSFSTKKSQPFPP
```

**IL2RG:**

```
IPWAP-ENLTLHK-LSESQLELNWNNRF-----LNHCLEHLVQYRTDWDHSWTEQSVDYRHK-------------------------FSLPSVDGQKRYTFRVRSRFNPLCGSAQHWSEWSHPIHWGSN
```

**CDON:**

```
PDAP-IILSPPQTHTPDTYNLVWRAGK---DGGLPINAYFVKYRKLDDGVGMLGSWHTVRVPGSENE-------------------
LHLAELEPSSLYEVLMVARSAAGEGQPAMLTFRTSKEKTASSKNTQASSPPV
PEAP-DRPTIST-ASETSVYVTWIPRA---NGGSPITAFKVEYKRMRTSNWLVAAEDIPPSKLS----------------------VEVRSLEPGSTYKFRVIAINHYGESFRSSASRPYQVVGFPNR
FSSRPITGP-HIAYTEA-VSDTQIMLKWTYIP-SSNNNTPIQGFYIYYRPTDSDNDSDYKRDVVEGSKQW---------------------
HMIGHLQPETSYDIKMQCFNEGGESEFSNVMICETKVKRVPGASEYPVKDLS
```

**COL12A1:**

```
PP-SDLNFKI-IDENTVHMSWAKPV------DPIVGYRITVDPTTDGPTKEFTLSASTTE------------------------TLLSELVPETEYVVTITSYDEVEESVPVIGQLTIQTGS
PP-SNLIAME-VSSKYVKLNWNPSP------SPVTGYKVILTPMTAGSRQHALSVGPQTTT-----------------------LSVRDLSADTEYQISVSAMKGMTSSEPISIMEKTQPMK
PP-KDLSFSE-VTSYGFKTNWSPAG------ENVFSYHITYKEAAGDDEVTVVEPASSTS------------------------VVLSSLKPETLYLVNVTAEYEDGFSIPLAGEETTEEVKG
AP-RNLKVTD-ETTDSFKITWTQAP------GRVLRYRIIYRPVAGGESREVTTPPNQRR------------------------RTLENLIPDTKYEVSVIPEYFSGPGTPLTGNAATEEVRG
NP-RDLRVSD-PTTSTMKLSWSGAP------GKVKQYLVTYTPVAGGETQEVTVRGDTTN------------------------TVLQGLKEGTQYALSVTALYASGAGDALFGEGTTLEER
GSP-QDLVTKD-ITDTSIGAYWTSAP------GMVRGYRVSWKSLYDDVDTGEKNLPEDAIH-----------------------TMIENLQPETKYRISVFATYSSGEGEPLTGDATTELSQD
SKTLKVDE-ETENTMRVTWKPAP------GKVVNYRVVYRPHGRGKQMVAKVPPTVTS------------------------TVLKRLQPQTTYDITVLPIYKMGEGKLRQGSGTTASRFK
SP-RNLKTSD-PTMSSFRVTWEPAP------GEVKGYKVTFHPTGDDRRLGELVVGPYDNT-----------------------VVLEELRAGTTYKVNVFGMFDGGESSPLVGQEMTTLSD
```

```
AP-SNLVISE-RTHRSFRVSWTPPS------DSVDRYKVEYYPVSGGKRQEFYVSRMETS------------------------TVLKDLKPETEYVVNVYSVVEDEYSEPLKGTEKTLPVP
  VVSLNIYD-VGPTTMHVQWQPVG-------GATGYILSYKPVKDTEPTRPKEVRLGPTVND---------------------MQLTDLVPNTEYAVTVQAVLHDLTSEPVTVREVTLPLP
RP-QDLKLRD-VTHSTMNVFWEPVP------GKVRKYIVRYKTPEEDVKEVEVDRSETS------------------------TSLKDLFSQTLYTVSVSAVHDEGESPPVTAQETTRPVPAP
   TNLKITE-VTSEGFRGTWDHGA------SDVSLYRITWAPFGSSDKMETILNGDENT------------------------LVFENLNPNTIYEVSITAIYPDESESDDLIGSERTLPILTTQ
GP-RNLQVYN-ATSNSLTVKWDPAS------GRVQKYRITYQPSTGEGNEQTTTIGGRQNS----------------------VVLQKLKPDTPYTITVSSLYPDGEGGRMTGRGKTKPLNTVRN
      LRVYD-PSTSTLNVRWDHAE------GNPRQYKLFYAPAAGGPEELVPIPGNTNY-----------------------AILRNLQPDTSYTVTVVPVYTEGDGGRTSDTGRTLMRG
 LARNVQVYN-PTPNSLDVRWDPAP------GPVLQYRVVYSPVDGTRPSESIVVPGNTRM----------------------VHLERLIPDTLYSVNLVALYSDGEGNPSPAQGRTLPRS
GP-RNLRVFG-ETTNSLSVAWDHAD------GPVQQYRIIYSPTVGDPIDEYTTVPGRRNN----------------------VILQPLQPDTPYKITVIAVYEDGDGGHLTGNGRTVGLL
PP-QNIHISD-EWYTRFRVSWDPSP------SPVLGYKIVYKPVGSNEPMEAFVGEMTS------------------------YTLHNLNPSTTYDVNVYAQYDSGLSVPLTDQGTTLYLN
 VTDLKTYQI--GWDTFCVKWSPHR-------AATSYRLKLSPADGTRGQEITVRGSETS------------------------HCFTGLSPDTDYGVTVFVQTPNLEGPGVSVKEHTTVKPTEAP
```

**EPHB4:**

```
PPSAP--RSVVSR-LNGSSLHLEWSAPL--ESGGREDLTYALRCRECRPGGSCAPCGGDLTFDPGPRDLVEPW-------------VVVRGLRPDFTYTFEVTALNGVSSLATGPVPFEPVNVTTDRE
 VPPAVSDIRVTR-SSPSSLSLAWAVPR---APSGAVLDYEVKYHEKGAEGPSSVRFLKTSENR----------------------AELRGLKRGASYLVQVRARSEAGYGPFGQEHHSQTLDE
```

**IL31RA:**

```
   KP-ENISCVY-YYRKNLTCTWSPGK-----ETSYTQYTVKRTYAFGEKHDNCTTNSSTSENRASCS-----------------FFLPRITIPDNYTIEVEAENGDGVIKSHMTYWRLENIAKT
  EPPKIFRVKPVLGIKRMIQIEWIKPE--LAPVSSDLKYTLRFRTVNSTSWMEVNFAKNRKDKNQT-------------------YNLTGLQPFTEYVIALRCAVKESKFWSDWSQEKMGMTEEE
APCGLELWRVLKPAEA-DGRRPVRLLWKKAR-GAPVLEKTLGYNIWYYPESNTNLTETMNTTNQ------------------------QLELHLGGESFWVSMISYNSLGKSPVATL
RIPAIQEKSFQCIEVMQACVAEDQLVVKWQSSA------LDVNTWMIEWFPDVDSEPTTLSWESVSQATNWT-------------------IQQDKLKPFWCYNISVYPMLHDKVGEPYSIQAYA
 KEGVPSEGP--ETKVEN-IGVKTVTITWKEIP-KSERKGIICNYTIFYQAEGGKGFSKTVNSSILQ-----------------------YGLESLKRKTSYIVQVMASTSAGGTNGTSINFKTLSF
```

**IL21R:**

```
   PDLVCYTDYLQTVICILEMWNLHP-----STLTLTWQDQYEELKDEATSCSLHRSAHNATHATYT-----------------CHMDVFHFMADDIFSVNITDQSGNYSQECGSFLLAESI
KPAPP-FNVTVTF---SGQYNISWRSDY------EDPAFYMLKGKLQYELQYRNRGDPWAVSPRRKLISVDSRSVS-----------LLPLEFRKDSSYELQVRAGPMPGSSYQGTWSEWSDPVIFQTQ
```

**IL11RA:**

```
PPARP--VVSCQA-ADYENFSCTWSPSQ----ISGLPTRYLTSYRKKTVLGADSQRRSPSTGPWPCPQDPLGAAR------------CVVHGAEFWSQYRINVTEVNPLGASTRLLDVSLQSILRPD
 PPQGLRVESVPGYPRRLRASWTYPA----SWPCQPHFLLKFRLQYRPAQHPAWSTVEPAGLE----------------------EVITDAVAGLPHAVRVSARDFLDAGTWSTWSPEAWGTPST
```

**IL12RB:**

```
  GP-RDLRCYR-ISSDRYECSWQYEG----PTAGVSHFLRCCLSSGRCCYFAAGSATRL-------------------------QFSDQAGVSVLYTVTLWVESWARNQTEKSPEVTLQLYNS
VKYEPPLGDIKVSK--LAGQLRMEWETPD---NQVGAEVQFRHRTPSSPWKLGDCGPQDDDTESC----------------------LCPLEMNVAQEFQLRRRQLGSQGSSWSKWSSPVCVPPE
NPPQPQVRFSVEQLGQDGRRRLTLKEQPTQLELPEGCQGLAPGTEVTYRLQLHMLSCPCKAKATRTLHL-----------------GKMPYLSGA-AYNVAVISSNQFGPGLNQTWHIPAD
THTEP-VALNISV--GTNGTTMYWPARA-------QSMTYCIEWQPVGQDGGLATCSLTAPQDPDPAGMATYSWS------------RESGAMGQEKCYYITIFASAHPEKLTLWSTVLSTYHFGGNAS
AAGTP-HHVSVKN-HSLDSVSVDWAPSL-LSTCPGVLKEYVVRCRDEDSKQVSEHPVQPTETQ------------------------VTLSGLRAGVAYTVQVRADTAWLRGVWSQPQRFSIEVQ
```

**Interferon gamma receptor2:**

```
 PAPQHPKIRLYN----AEQVLSWEPVA--LSNSTRPVVYQVQFKYTDSKWFTADIMSIGVNCTQITATECDFT-------------AASPSAGFPMDFNVTLRLRAELGALHSAWVTMPWFQHYRNV
TVGPP-ENIEVTP--GEGSLIIRFSSPFDIADTSTAFFCYYVHYWEKGGIQQVKGPFRSNS-------------------------ISLDNLKPSRVYCLQVQAQLLWNKSNIFRVGHLSNISCYETM
```

**Nephrin:**

```
   PP-SGLKVVS-LTPHSVGLEWKPGF----DGGLPQRFCIRYEALGTPGFHYVDVVPPQATT----------------------FTLTGLQPSTRYRVWLLASNALGDSGLADKGTQLPITTPGL
```

**IL7R:**

```
   AP-FDLSVVYREGANDFVVTFNTSH-------LQKKYVKVLMHDVAYRQEKDENKWTHVNLSSTKLT-----------------LLQRKLQPAAMYEIKVRSIPDHYFKGFWSEWSPSYYFRTPEI
```

**Anosmin-1:**

```
 PLKPRKELRFTE-LQSGQLEVKWSSKF---NISIEPVIYVVQRRWNYGIHPSEDDATHWQTVAQTTDER-----------------VQLTDIRPSRWYQFRVAAVNVHGTRGFTAPSKHFRSSKDPS
APPAPANLRLANSTVNSDGSVTVTIVWDLPE---EPDIPVHHYKVFWSWMVSSKSLVPTKKKRRKTTDGFQNS-----------------VILEKLQPDCDYVVELQAITYWGQTRLKSAKVSLHFTSTHAT
  RPTRPLEVGAPFYQ-DGQLQVKVYWKKTE-----DPTVNRYHVRWFPEACAHNRTTGSEASSGMTHENY-------------------IILQDLSFSCKYKVTVQPIRPKSHSKAEAVFFTTPPCS
   KPENLSASFIVQDVNITGHFSWKMAK--ANLYQPMTGFQVTWAEVTTESRQNSLPNSIISQSQILPSDHYV--------------LTVPNLRPSTLYRLEVQVLTPGGEGPATIKTFRTPELPPS
```

**MYBPC3:**

```
 IDVPDAPAAPKISNVGEDSCTVQWEPPA--YDGGQPILGYILERKKKKSYRWMRLNFDLIQELS----------------------HEARRMIEGVVYEMRVYAVNAIGMSRPSPASQPFMPIGPP
  SEP--THLAVED-VSDTTVSLKWRPPE--RVGAGGLDGYSVEYCPEGCSEWVAALQGLTEHTS----------------------ILVKDLPTGARLLFRVRAHNMAGPGAPVTTTEPVTVQEI
KPSPP-QDLRVTD-AWGLNVALEWKPPQ--DVGNTELWGYTVQKADKKTMEWFTVLEHYRRTH------------------------CVVPELIFGNGYYFRVFSQNMVGFSDRAATTKEPVFIPRPG
```

```
SPEG:
            PDGAP-QVVAVTG----RMVTLTWNPPR---SLDMAIDPDSLTYTVQHQVLGSDQWTALVTGLREPG--------------------WAATGLRKGVQHIFRVLSTTVKSSSKPSPPSEPVQLLEH
            KLAPP--EVPQTY---QDTALVLWKPGD-----SRAPCTYTLERRVDGESVWHPVSSGIPDCY------------------------YNVTHLPVGVTVRFRVACANRAGQGPFSNSSEKVFVRGTQDS
Thrombopoietin receptor:
               GPRDPKNSTG---PTVIQLIATETC-CPALQRPHSASALDQSPCAQPTMPWQDGPKQTSPSREASALTAEGGS-----------CLISGLQPGNSYWLQLRSEPDGISLGGSWGSWSLPVTVDLPG
                PTPNLHWRE-ISSGHLELEWQHPS---SWAAQETCYQLRYTGEGHQDWKVLEPPLGAR-------------------------GGTLELRPRSRYRLQLRARLNGPTYQGPWSSWSDPTRVETAT
OSMR:
               NP-FSVNFEN-VNATNAIMTWKVHS-----IRNNFTYLCQIELHGEGKMMQYNVSIKVNGE-----------------------YFLSELEPATEYMARVRCADASHFWKWSEWSGQNFTTLEA
         APSEAPDVWRIVSLEP--GNHTVTLFWKPLS-KLHANGKILFYNVVVENLDKPSSSELHSIPAPA-------------------------NSTKLILDRCSYQICVIANNSVGASPASVIVISADPEN
               KEVEEERIAG--TEGGFSLSWKPQP------GDVIGYVVDWCDHTQDVLGDFQWKNVGPNTTSTV-------------------ISTDAFRPGVRYDFRIYGLSTKRIACLLEKKTGYSQEL
           APSDNP--HVLVDT-LTSHSFTLSWKDYS-----TESQPGFIQGYHVYLKSKARQCHPRFEKAVLSDGSECCKYKIDNPEEKA-----LIVDNLKPESFYEFFITPFTSAGEGPSATFTKVTTPDE
GHR:
      NPGLKTNSSKEP-KFTKCRS-PERETFSCHWTDEV--HHGTKNLGPIQLFYTRRNTQEWTQEWKECPDYVSAGENSCY---------------FNSSFTSIWIPYCIKLTSNGGTVDEKCFSVDEIVQPD
         PPIALNWTLLNVSLTG--IHADIQVRWEAPRNADIQKGWMVLEYELQYKEVNETKWKMMDPILTTS------------------------VPVYSLKVDKEYEVRVRSKQRNSGNYGEFSEVLYVTLPQM
CSF3R:
               IP-HNLSCLMNLTTSSLICQWEPGP----ETHLPTSFTLKSFKSRGNCQTQGDSILDCVPKDGQSHCC----------------IPRKHLLLYQNMGIWVQAENALGTSMSPQLCLDPMDVVKLEP
          MLRTMDP-SPEAAPP--QAGCLQLCWEPWQ---PGLHINQKCELRHKPQRGEASWALVGPLPLEALQ----------------------YELCGLLPATAYTLQIRCIRWPLPGHWSDWSPSLELRTTER
            APTVRLDTWWRQRQLDPRTVQLFWKPVP-LEEDSGRIQGYVVSWRPSGQAGAILPLCNTTE-------------------------LSCTFHLPSEAQEVALVAYNSAGTSRPTPVVFSESRGP
                ALTRLHAMA-RDPHSLWVGWEPPN------PWPQGYVIEWGLGPPSASNSNKTWRMEQNGRATGF------------------LLKENIRPFQLYEIIVTPLYQDTMGPSQHVYAYSQEM
           APSHAP--ELHLKH-IGKTWAQLEWVPEP-PELGKSPLTHYTIFWTNAQNQSFSAILNASSRG-------------------------FVLHGLEPASLYHIHLMAASQAGATNSTVLTLMTLTPEGS
CSF2R:
        YPNSGREGTAAQNFSCFI-YNADLMNCTWARGP----TAPRDVQYFLYIRNSKRRREIRCPYYIQDSGTHVGCHL-----------------DNLSGLTSR-NYFLVNGTSREIGIQFFDSLLDTKKIERFN
               PP-SNVTVRC--NTTHCLVRWKQPRTYQKLSYLDFQYQLDVHRKNTQPGTENLLINVSGDLENR--------------------YNFPSSEPRAKHSVKIRAADVRILNWSSWSEAIEFGSDDG
Tie2/TEK:
           KPLNAP-NVIDTGHN--FAVINISSEPYF----GDGPIKSKKLLYKPVNHYEAWQHIQVTNEI-------------------------VTLNYLEPRTEYELCVQLVRRGEGGEGHPGPVRRFTTASI
            GLPPP-RGLNLLPK-SQTTLNLTWQPIF-----PSSEDDFYVEVERRSVQKSDQQNIKVPGNLTS----------------------VLLNNLHPREQYVVRARVNTKAQGEWSEDLTAWTLSD
           ILPPQP-ENIKISNI-THSSAVISWTILD-----GYSISSITIRYKVQGKNEDQHVDVKIKNATITQ---------------------YQLKGLEPETAYQVDIFAENNIGSSNPAFSHELVTLPES
```

#### 5.e Identity of mapped 3Fn variants: disease proteins and disease variants

Summary of included proteins containing naturally occurring variants associated with diseases in domains shared with SORLA.

| Gene | Protein | Uniprot | #dom | Associated Diseases/Syndromes (CODE) | Variants mapped onto big alignments (from Uniprot entries except where a specific ref is provided) | #variants | #positions | Citations for variants discussed in main text |
|---|---|---|---|---|---|---|---|---|
| USH2A | Usherin | O75445 | 34 | Usher syndrome 2A (USH2A)<br>Retinitis pigmentosa 39 (RP39) | **USH2A**: P1059L, P1212L, A1953G, K2080N, H2116R, C2128Y/C2128F, S2196T, E2238A, A2249D, S2260P, R2292H, R2354H, V2562K, S2639P, D2738N, **W2744C**, G2752R, F2786S, A2795S, R3124G, E3448K, T3462I, W3479C, P3504T, D3515G, **W3521R**, G3529S, G3546R, T3571M, Y3747C, **I3844M**, N3894D, G3895E, R3904K, T3976M, S4054I, R4115C, S4174R, P4232R, P4269R, T4337M, I4386F, T4425M, V4433L, TR4439I, Y4487C, R4570H, Q4592H, Q4662E, G4692R, G4763R, **L4795R**, C4808R, G4817R, P4818L, T4918M<br>**RP39**: F1442S, P1978S, D2237Y, R2460H, R2573H, N2930K, L3606P, G3618S, S3669R, R3719H, N4094K, R4115C, R4192H, H4248N, T4425M, M4447V, R4674G, L4840P, T4844M | 74 | 73 | **W2744C** [167]<br>**W3521R** [168]<br>**I3844M** [193] |
| FN1 | Fibronectin | P02751 | 17 | Glomerulopathy with fibronectin deposits 2 (GFND2) | **GFNDS2**: **Y973C**, **W1925R**, **L1974R** | 3 | 3 | **Y973C** causal of GFND2 [170]. Segregate with disease in four unrelated families<br>**L1974R** causal [170]<br>**W1925R** causal [170] |
| TNXB | Tenascin-X | P22105 | 30 | Ehlers-Danlos Syndrome, classic-like (EDSCLL)<br>Vesicourecteral reflux 8 (VUR8) | **EDSCLL**: V1108M<br>**VUR8**: T1244R, V3212I | 3 | 3 | |
| TNC | Tenascin | P24821 | 15 | Deafness, autosomal dominant, 56 (DFNA56) | **DFNA56**: V1773M, T1796S | 2 | 2 | |
| L1CAM | Neural cell adhesion molecule L1 | P32004 | 5 | Hydrocephalus due to stenosis of the aqueduct of Sylvius (HSAS)<br>Mental retardation, aphasia, shuffling gait, and adducted thumbs syndrome (MASA) | **HSAS**: K655E, A691T, G698R, M741T, V752M, V768F, **Y784C**, L935P, P941L, **W1036L**, Y1070C<br>**MASA**: R632P, S674C, A691D, G698R, V952M, D770N, P941L | 16 | 15 | **Y784C** [178]<br>**W1036L** defective protein [169] |
| TEK | Angiopoietin-1 receptor | Q02763 | 2 | Glaucoma 3, primary congenital, E (GLC3E) | **GLC3E**: **Y611C** | 1 | 1 | **Y611C** in a pedigree and mutant protein can't undergo ligand induced phosphorylation [175] |
| INSR | Insulin receptor | P06213 | 3 | Rabson-Mendenhall syndrome (RMS)<br>Leprechaunism (LEPRCH)<br>Diabetes mellitus, non-insulin-dependent (NIDDM) | **RMS**: S635L, S835I, A842V, P874L, N878S<br>LEPRCH: V657F, W659R, **Y818C**, **I925T**, **R926W**, T937M<br>**NIDDM**: T858A | 12 | 12 | **Y818C** abolishes post-translational processing [176,177] |

| | | | | | | | | |
|---|---|---|---|---|---|---|---|---|
| | | | | | | | | **I925T** abolish post-translational processing; abolish insulin binding [177,180] **R926W** markedly impaired isulin binding, impaired post-translational processing [177,180] |
| IGF1R | Insulin-like growth factor 1 receptor | P08069 | 4 | Insulin-like growth factor 1 resistance (IGF1RES) | **IGF1RES**: R739Q, Y865C | 2 | 2 | |
| DCC | Netrin receptor DCC | P43146 | 6 | Mirror movements 1 (MRMV1)<br>Gaze palsy, familial horizontal, with progressive scoliosis, 2, with impaired intellectual development (HGPPS2) | **MRMV1**: R597P, M743L, V754G, V793G, G805E, A893T<br>**HPPPS2**: Q691K | 7 | 7 | |
| ROBO3 | Roundabout homolog 3 | Q96MS0 | 3 | Gaze palsy, familial horizontal, with progressive scoliosis, 1 (HGPPS1) | **HGPPS1**: R703P, S705P | 2 | 2 | |
| ROBO4 | Roundabout homolog 4 | Q8WZ75 | 2 | Aortic valve disease 3 (AOVD3) | **AOVD3**: **Y280S**, H411Q | 2 | 2 | |
| LEPR | Leptin receptor | P48357 | 4 | Leptin receptor deficiency (LEPRD) | **LEPRD**: C604G, L786P | 2 | 2 | |
| PRLR | Prolactin receptor | P16471 | 2 | Multiple fibroadenomas of the breast (MFAB)<br>Hyperprolactinemia (HPRL) | **MFAB**: I170L<br>**HPRL**: H212R | 2 | 2 | |
| SPEG | Striated muscle preferentially expressed protein kinase | Q15772 | 2 | Myopathy, centronuclear, 5 (CNM5) | **CNM5**: G2757V | 1 | 1 | |
| ITGB4 | Integrin beta4 | P16144 | 4 | Epidermolysis bullosa letalis, with pyloric atresia (EB-PA) | **EB-PA**: R1225H, R1281W | 2 | 2 | |
| CRLF1 | Cytokine receptor-like factor 1 | O75462 | 2 | Crisponi/Cold-induced sweating syndrome 1 (CISS1) | **CISS1**: P138L, S145P, R216C, F268S, W284C, **R312P**, R340C | 7 | 7 | **R312P** [181] |
| COL6A3 | Collagen alpha-3(VI) chain | P12111 | 1 | Dystonia 27 (DYT27) | **DYT27**: R3043H, P3082R | 2 | 2 | |
| IL2RG | Cytokine receptor common subunit gamma | P31785 | 1 | Severe combined immunodeficiency X-linked T-cell-negative/B-cell-positive/NK-cell-negative (XSCID)<br>X-linked combined immunodeficiency (XCID) | **XSCID**: A156V, L162H, L172P/L172Q, C182R, L183S, **R224W**, R226C/R226H, F227C, L230P, C231Y, G232R, W240C, S241I<br>**XCID**: R222C | 16 | 14 | **R224W** [182] |
| CDON | Cell adhesion molecule-related/down-regulated by oncogenes | Q4KMG0 | 3 | Holoprosencephaly 11 (HPE11) | **HPE11**: T684S, P689A, V691M, V780E, T790A, S940R | 6 | 6 | |
| COL12A1 | Collagen alpha-1(XII) chain | Q99715 | 18 | Bethlem myopathy 2 (BTHLM2) | **BTHLM2**: R1965C | 1 | 1 | |
| EPHB4 | Ephrin type-B receptor 4 | P54760 | 2 | Capillary malformation-arteriovenous malformation 2 (CMAVM2) | **CMAVM2**: V469G, G516R | 2 | 2 | |
| IL31RA | Interleukin-31 receptor subunit alpha | Q8NI17 | 5 | Amyloidosis, primary localized cutaneous, 2 (PLCA2) | **PLCA2**: S489F | 1 | 1 | |
| IL21R | Interleukin-21 receptor | Q9HBE5 | 2 | Immunodeficiency 56 (IMD56) | **IMD56**: **R201L** | 1 | 1 | **R201L** defective trafficking, misfolding and impaired processing [160,161] |

| | | | | | | | | |
|---|---|---|---|---|---|---|---|---|
| IL11RA | Interleukin-11 receptor subunit alpha | Q14626 | 2 | Craniosynostosis and dental anomalies (CRSDA) | **CRSDA**: P221R, S245C, R296W | 3 | 3 | |
| IL12RB1 | Interleukin-12 Receptor subunit beta | P42701 | 5 | Immunodeficiency 30 (IMD30) | **IMD30**: **R213W** | 1 | 1 | **R213W** [183] |
| IFNGR2 | Interferon gamma receptor 2 | P38484 | 2 | Immunodeficiency 28 (IMD28) | **IMD28**: **R114C**, S124F, G141R, T168N, G227R | 5 | 5 | **R114C** misfolding and abnormal glycosylation, mistrafficking, reduced response to INFG [184,185] |
| NPHS1 | Nephrin | O60500 | 1 | Nephrotic syndrome 1 (NPHS1) | **NPHS1**: R976S, S1016N, G1020V | 3 | 3 | |
| IL7R | Interleukin-7 receptor subunit alpha | P16871 | 1 | Severe combined immunodeficiency autosomal recessive T-cell-negative/B-cell-positive/NK-cell-positive (T(-)B(+)NK(+) (SCID) | **SCID**: P132S | 1 | 1 | |
| ANOS1 | Anosmin-1 | P23352 | 4 | Hypogonadotropic hypogonadism 1 with or without anosmia (HH1) | **HH1**: R262P, N267K, N304S, S396L, E514K, F517L, **W571R**, V587L | 8 | 8 | **W571R** in patients with Kallmann Syndrome/HH1 [171] |
| MYBPC3 | Myosin-binding protein C, cardiac-type | Q14896 | 3 | Cardiomyopathy, familial hypertrophic 4 (CMH4) | **CMH4**: D770N, V771M, **W792R**, R810H, K811R, R820Q, A833T/A833V, R834W, P873H, N948T, T957S, T958I, I1131T | 14 | 13 | **W792R** in patients with CMH4 [172] |
| MPL | Thrombopoietin receptor | P40238 | 2 | Congenital amegakaryocytic thrombocytopenia (CAMT) | **CAMT**: **R257L**, P257T, W435C | 3 | 3 | **R257L** [186] |
| OSMR | Oncostatin-M-specific receptor subunit beta | Q99650 | 4 | Amyloidosis, primary localized cutaneous, 1 (PLCA1) | **PLCA1**: G618A, D647V, I691T, P694L, K697T | 5 | 5 | |
| GHR | Growth hormone receptor | P10912 | (2) | Laron syndrome (LARS)<br>Growth hormone insensitivity, partial (GHIP) | **LARS**: C56S, S58L, **W68R**, R89K, F114S, V143A, P149Q, V162D, D170H, I171T, Q172P, V173G, **Y226C**, R229G, S244I<br>**GHIP**: E62K, R179C | 17 | 17 | **Y226C** is causal of severe growth retardation [179]<br>**W68R** in LARS patient [173] |
| CSF3R | Granulocyte colony-stimulating factor receptor | Q99062 | 5 | Neutropenia, severe congenital 7, autosomal recessive (SCN7) | **SCN7**: **R308C** | 1 | 1 | **R308C** Decreased localization at cell surface and less signaling. Impaired glycosylation [187] |
| CSF2RA | Granulocyte-macrophage colony-stimulating factor receptor subunit alpha | P15509 | 2 | Pulmonary surfactant metabolism dysfunction 4 (SMDP4) | **SMDP4**: G196R | 1 | 1 | |
| | | | | | | **229** | 224 | |

#### 5.f 3Fn disease variants listed according to domain positions

Disease-mutations domain-mapping analysis with identification of pathogenic variants in other proteins with 3Fn-domains (as listed in **Supplemental Information 5e**). Here variants are mapped onto domain positions following alignment of internally repeated sequences in the SORLA domain sequences. The number of hits for each position depicted in the bar diagram of **Supplemental Figure 5e**.

| **3Fn-domain positions** | Number of hits | Identified variants | | Priority |
|---|---|---|---|---|
| 1 | | | | |
| 2 | | | | |
| 3 | | | | |
| 4 | | | | |
| 5 | | | | |
| 6; P | 6 | p.P1059L(USH2A), p.N2930K(USH2A), p.P3504T(USH2A), p.R739Q(IGF1R), p.A156V(IL2RG), p.P873H(MYBPC3) | Loss of Pro *R739 of IGF1R part of furin site* | high |
| 7; P | 4 | p.P4269R(USH2A), p.P138L(CRLF1), p.P221R(IL11RA), p.P132S(IL7R) | Loss of Pro | high |
| 8 | 0 | | | |
| 9 | 1 | p.T858A(INSR) | | |
| 10 | 1 | p.R1255H(ITGB4) | | |
| 11; **ø** | 2 | p.A2249D(USH2A), p.V162D(GHR) | Loss of hydrophobic | moderate |
| 12 | 0 | | | |
| 13; **ø** | 2 | p.L162H(IL2RG), p.C56S(GHR) | Loss of hydrophobic | moderate |
| 14 | 2 | p.T3976M(USH2A), p.L4840P(USH2A) | | |
| 15 | 3 | p.S145P(CRLF1), p.N304S(ANOS1), p.S58L(GHR) | | |
| 16 | 0 | | | |
| 17 | 0 | | | |
| 18 | 1 | p.S635L(INSR) | | |
| 19 | 4 | p.D2738N(USH2A), p.R3124G(USH2A), p.D3515G(USH2A), p.E62K(GHR) | | |
| 20 | 3 | p.S2639P(USH2A), p.T4844M(USH2A), p.D170H(GHR) | | |
| 21; **ø** | 2 | p.L3606P(USH2A), p.I171T(GHR) | Loss of hydrophobic | moderate |
| 22 | 3 | p.R632P(L1CAM), p.L935P(L1CAM), p.Q172P(GHR) | | |
| 23; **ø** | 4 | p.M743L(DCC), p.L172P(IL2RG), p.L172Q(IL2RG), p.V173G(GHR) | Loss of hydrophobic | moderate |
| 24 | 2 | p.S2260P(USH2A), p.S4174R(USH2A) | | |
| 25; **W** | 7 | p.W2744C(USH2A), p.W3521R(USH2A), p.W1036L(L1CAM), p.W1925R(FN1), p.W571R(ANOS1), p.W729R(MYBPC3), p.W68R(GHR) | Loss of Trp | high |
| 26 | 0 | | | |
| 27; p | 1 | p.D647V(OSMR) | | |
| 28; p | 3 | p.P1978S(USH2A), p.P941L(L1CAM), p.P874L(INSR) | Loss of Pro | high |
| 29 | 1 | p.R179C(GHR) | | |
| 30 | 0 | | | |
| 31 | 1 | p.R2460H(USH2A) | | |
| 32 | 1 | p.M741T(L1CAM) | | |
| 33 | 3 | p.K2080N(USH2A), p.R2354H(USH2A), p.Q4662E(USH2A) | | |
| 34 | 2 | p.S245C(IL11RA), p.T168N(INGR2) | | |
| 35 | 4 | p.N4094K(USH2A), p.N3894D(USH2A), p.N878S(INSR), p.F268S(CRLF1) | | |
| 36; G | 5 | p.G2752R(USH2A), p.G3529S(USH2A), p.G3618S(USH2A), p.G3895E(USH2A), p.G4763R(USH2A) | Loss of Gly | high |
| 37 | 2 | p.V2562A(USH2A), p.V754G(DCC) | | |
| 38 | 2 | p.I4386F(USH2A), p.C182R(IL2RG) | | |
| 39 | 1 | p.L183S(IL2RG) | | |

| | | | | |
|---|---|---|---|---|
| 40 | 0 | | | |
| 41; **Y** | 3 | p.Y865C(IGF1R), p.Y280S(ROBO4), p.Y217D(ANOS1)[194] | Loss of Tyr | high |
| 42 | 3 | p.R751P(L1CAM), p.R703P(ROBO3), p.R1965C(COL12A1) | | |
| 43; **ø** | 4 | p.V752M(L1CAM), p.V657F(INSR), p.V469G(EPHB4), p.V587L(ANOS1) | Loss of hydrophobic | moderate |
| 44 | 2 | p.S705P(ROBO3), p.R976S(NPHS1) | | |
| 45; **Y** | 6 | p.R3719H(USH2A), p.R3904K(USH2A), p.R4192H(USH2A), p.R4674G(USH2A), p.W659R(INSR), p.R810H(MYBPC3) | | moderate |
| 46 | 2 | p.R4570H(USH2A), p.K811R(MYBPC3) | | |
| 47 | 1 | p.I170L(PRLR) | | |
| 48 | 4 | p.E3448K(USH2A), p.R2573H(USH2A), p.K655E(L1CAM), p.R89K(GHR) | | |
| 49 | 0 | | | |
| 50 | | *Alignment not precise enough* | | |
| 51 | | *Alignment not precise enough* | | |
| 52 | | *Alignment not precise enough* | | |
| 53 | | *Alignment not precise enough* | | |
| 54 | | *Alignment not precise enough* | | |
| 55 | | *Alignment not precise enough* | | |
| 56 | | *Alignment not precise enough* | | |
| 57 | | *Alignment not precise enough* | | |
| 58 | | *Alignment not precise enough* | | |
| 59 | | *Alignment not precise enough* | | |
| 60 | | *Alignment not precise enough* | | |
| 61 | | *Alignment not precise enough* | | |
| 62 | | *Alignment not precise enough* | | |
| 63 | | *Alignment not precise enough* | | |
| 64 | | *Alignment not precise enough* | | |
| 65 | | *Alignment not precise enough* | | |
| 66 | | *Alignment not precise enough* | | |
| 67 | | *Alignment not precise enough* | | |
| 68 | | *Alignment not precise enough* | | |
| 69 | | *Alignment not precise enough* | | |
| 70 | | *Alignment not precise enough* | | |
| 71 | | *Alignment not precise enough* | | |
| 72; **ø** | 4 | p.S674C(L1CAM), p.R3043H(COL6A3), p.V780E(CDON), p.F114S(GHR) | | moderate |
| 73 | 1 | p.T3462I(USH2A) | | |
| 74; **ø** | 2 | p.A833T(MYBPC3), p.A833V(MYBPC3) | | moderate |
| 75 | 1 | p.R834W(MYBPC3) | | |
| 76 | 0 | | | |
| 77; **L** | 1 | p.L4795R(USH2A) | | high |
| 78 | 1 | p.H411Q(ROBO4) | | |
| 79; P | 3 | p.P1212L(USH2A), p.P4232R(USH2A), p.I1131T(MYBPC3) | Loss of Pro | high |
| 80 | 2 | p.F2786S(USH2A), p.C604G(LEPR) | | |
| 81 | 1 | p.V1773M(TNC) | | |
| 82 | 3 | p.Q691K(DCC), p.T790A(CDON), p.S489F(IL31RA) | | |
| 83; **Y** | 6 | p.H2116R(USH2A), p.Y784C(L1CAM), p.Y818C(INSR), p.Y973C(FN1), p.Y226C(GHR), p.Y611C(TEK) | Loss of Tyr | high |
| 84 | 1 | p.V793G(DCC) | | |
| 85; **ø** | 2 | p.F1442S(USH2A), p.Y3747C(USH2A) | | moderate |
| 86 | 4 | p.R597P(DCC), p.R222C(IL2RG), p.R262P(ANOS1), p.R229G(GHR) | Loss of Arg | |
| 87; **ø** | 3 | p.I3844M(USH2A), p.I925T(INSR), p.L1974R(FN1) | Loss of hydrophobic | moderate |
| 88 | 8 | p.R926W(INSR), p.R312P(CRLF1), p.R224W(IL2RG), p.R201L(IL21R), p.R213W(IL12RB), p.R114C(INGR2), p.R257L(MPL), p.R308C(CSF3R) | Loss of Arg | high |
| 89; **a** | 2 | p.A2795S(USH2A), p.A691T(L1CAM) | Loss of Ala (or hyd.) | moderate |
| 90 | 6 | p.W3479C(USH2A), p.C4808R(USH2A), p.R226C(IL2RG), p.R226H(IL2RG), p.R296W(IL11RA), p.S1016N(NPHS1) | | |
| 91 | 6 | p.T3571M(USH2A), p.T4337M(USH2A), p.T4425M(USH2A), p.F227C(IL2RG), p.N276K(ANOS1), p.N948T(MYBPC3) | | |
| 92 | 1 | p.R216C(CRLF1) | | |
| 93 | 1 | p.H212R(PRLR) | | |

| | | | | |
|---|---|---|---|---|
| 94; g | 2 | p.L230P(IL2RG), p.G1020V(NPHS1) | Loss of Gly | moderate |
| 95 | 4 | p.H4248N(USH2A), p.C2128Y(USH2A), p.C2128F(USH2A), p.C231Y(IL2RG) | | |
| 96; g | 7 | p.T4918M(USH2A), p.G698R(L1CAM), p.G805E(DCC), p.G232R(IL2RG), p.G516R(EPHB4), p.G1020V(NPHS1), p.G2757V(SPEG) | Loss of Gly | high |
| 97 | 0 | | | |
| 98; s | 3 | p.S3669R(USH2A), p.S4054I(USH2A), p.S124F(INGR2) | Loss of Ser? | |
| 99 | | *Alignment not precise enough* | | |
| 100 | | *Alignment not precise enough* | | |
| 101 | | *Alignment not precise enough* | | |
| 102 | | *Alignment not precise enough* | | |
| 103 | | *Alignment not precise enough* | | |
| 104 | | *Alignment not precise enough* | | |
| 105 | | *Alignment not precise enough* | | |
| 106 | | *Alignment not precise enough* | | |
| 107 | | *Alignment not precise enough* | | |
| 108 | | *Alignment not precise enough* | | |
| 109 | | *Alignment not precise enough* | | |
| 110 | | *Alignment not precise enough* | | |
| 111 | | *Alignment not precise enough* | | |
| | **173*** | | | |

* A number of identified variants (listed in **Supplemental Information 5e**) map within 3Fn-domain sequences where the alignment does not allow unambiguous domain position identification

### 6 Transmembrane and cytoplasmic domains (residues 2122-2214)

#### 6.a Sequence details

*Membrane-anchoring domain*

Immediately following the 3Fn-domains is a 16 amino acid long stalk region (residues 2122-2137) suggested to lift the ectodomain a short distance from the plasma membrane, enabling TACE-dependent cleavage, leading to ectodomain shedding when mature SORLA is at the cell surface [162,195]. After the stalk region, SORLA contains a 23-amino acid single-pass transmembrane (TM) domain (residues 2138-2160), presumably alpha-helical, and a cytoplasmic domain (CD; often referred to as the 'tail') including 54 amino acids (residues 2161-2214) (**Fig. 3**).

The TM-domains of mammalian SORLA proteins are highly conserved during evolution, with human and insects sharing >95% sequence identity [196] (**Supplemental Figure S6**). The number of amino acids in the TM-domain may influence the localization of a protein within the cell [197]. With this reasoning, the number of residues in the SORLA TM-domain might underlie its manifestation in the membranes of the trans-Golgi and trafficking network. Similar to most membrane-anchored proteins, the amino acid composition of the SORLA TM-domain is mainly characterized by hydrophobic residues that form non-polar interactions that stabilize the helix structure within the phospholipid bilayer. Therefore, the preservation of the hydrophobicity of TM-domains is key for proper insertion of the protein into the membrane: substitutions of hydrophobic residues with polar or charged residues in the LDLR TM-domain prevented membrane insertion, which was causative for familial hypercholesterolemia [198,199]. Thus far, it is unknown whether mutations in the SORLA-TM domain will have similar effects.

*The intracellular cytoplasmic domain*

Similar to most members of the LDLR and VPS10p receptor families, a short (~50 amino acid) cytoplasmic tail constitutes the intracellular part of SORLA. This tail determines the complex intracellular sorting itinerary that SORLA follows (**Supplemental Figure S6**). Binding to different intracellular adaptor proteins via short motifs in the tail determines key sorting steps of SORLA and its cargo, such as endocytosis and intracellular trafficking between Golgi/TGN and endo-lysosomal compartments (see below) (**Supplemental Figure S6**). Deletion of the entire tail leads to the direct vesicular transportation of the receptor from the TGN to the plasma membrane and, upon cleavage by sheddase (i.e. TACE), its release into extracellular space [67,200].

*GGA*

The very C-terminal part of SORLA contains a DXXLL-like motif ($^{2208}$DVPMV$^{2212}$), which is a target for the Golgi-localized proteins GGA1, GGA2 and GGA3. These adaptors regulate trafficking of a set of membrane-spanning proteins from Golgi to endosomal compartments and – at least in the case of GGA3 – from endosome towards lysosome [201,202 203]. This suggests that SORLA can sort directly from Golgi to the endosome after its synthesis. The minimal motif required for binding of GGA adaptors to the SORLA cytoplasmic domain is characterized by two acidic residues $^{2207}$DD$^{2208}$ preceding methionine M$^{2211}$ of the hydrophobic cluster at the very C-terminal end [204]. In 2009, the structure of part of the SORLA tail was determined in complex with the VHS domain of GGA1 [205] confirming that the GGA adaptor directly binds these residues. This highly conserved motif is crucial for SORLA function: a mutation in this motif yields a receptor with compromised trafficking to the cell surface [200,206,207]. The overlapping $^{2212}$VIA$^{2214}$ motif was recently shown to bind PICK1, suggesting that also this protein is capable of regulating SORLA's intracellular itinerary [208].

*FANSHY*

The SORLA tail also includes the $^{2172}$FANSHY$^{2177}$ motif, which is strictly conserved from human to insects suggesting an indispensable physiological function (**Supplemental Figures S6**). This motif is similar to the (F/Y)XNPXY motif as identified in the cytoplasmic domain of many proteins including APP, LDLR, and LRP, in which it serves as an endocytosis signal [209]. However, in SORLA, the Pro residue is absent, and here, the motif is instead required for binding the retromer core complex (VPS26, VPS29 and VPS35), which is involved in the recycling of various transmembrane receptors between the endosomal network to the cell surface or in the retrograde trafficking to the Golgi/TGN [210]. The Phe-2172 of the $^{2172}$FANSHY motif is essential for association of the SORLA tail with the VPS26 subunit of the retromer complex [211].

The tail-motif of several receptors is the simple NXXY motif, i.e. also without a proline residue as well as the preceding aromatic F/Y residue, and these receptors can be bound to endosomal recycling proteins such as SNX17, SNX27, and SNX31 [212,213], raising the possibility that the FA**N**SH**Y** site in the SORLA tail may interact not also interact with such sorting complexes for endosomal sorting, which may assist SORLA cycling from endosomes to the cell surface [214]. Interestingly, a fragment of the intracellular domain spanning the FANSHY motif folds into an amphiphatic $\alpha$-helical structure with the potential for sensing membrane curvature as yet another determinant of SORLA intracellular transport [215].

*Acidic*

An acidic cluster in the tail of SORLA (corresponding to residues $^{2190}$DDLGEDDED$^{2198}$) is reported to bind cytoplasmic adaptor proteins including PACS1, depending on cellular localization and cell type [200,216-218]. In clathrin-vesicles, SORLA-binding goes via the AP1 and AP2 adaptor-binding proteins, which bind the EXXXLL-like motif ($^{2197}$EDAPMI$^{2202}$). Interactions with the AP1, AP2 and PACS1 adaptor proteins regulate mainly retrograde sorting pathways from cell surface to endosome (AP2) or endosomes to TGN (PACS1). However, AP1 is also involved in directing cargo in the anterograde direction from TGN towards endosomes, and described to do so partly in concert with GGAs [219,220]. Receptors carrying substitutions within this acidic motif have a strong defect in endocytosis, due to lack of AP2 binding [200,216]. It has also been reported that the relatively unknown HSPA12A cytosolic protein binds to the acidic motif within the tail of SORLA [221].

*Phosphorylation*

Twelve of the 54 amino acids of the SORLA tail are potential targets of phosphorylation (Ser/Thr/Tyr) and several of these residues occur in consensus target sequences for specific kinases [222], suggesting phosphorylation might play important functions in the sorting of SORLA (**Fig. 3**). For example, the ROCK2 kinase can phosphorylate Ser ($S^{2206}$), which modifies the cytoplasmic domain of SORLA such that ectodomain shedding of SORLA is increased [223]. Particularly, the presence of both a tyrosine and a serine in the FANSHY motif may provide possible phosphorylation-dependent regulatory mechanisms of SORLA endosomal sorting. The significance of such modifications is currently under investigation.

Andersen
Supplemental Figure S6 - Tail-domain

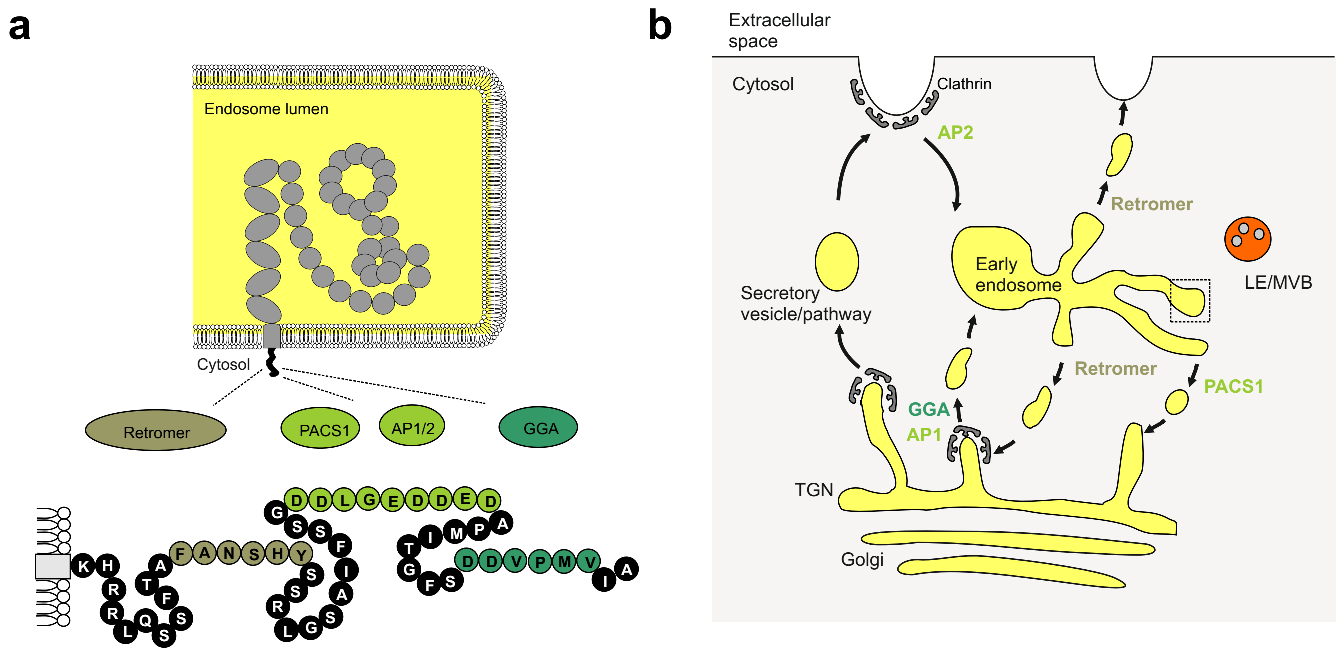

c

| | | Stalk | Transmembrane | Cytoplasmic tail |
|---|---|---|---|---|
| | | | | 1 10 20 30 40 50 |
| Human | 2122-2214 | GADASATQAARSTDVA | AVVVPILFLILLSLGVGFAILYT | KHRRLQSSFTA**FANSHY**SSRLGSAIFSSG**DDLGEDDED**APMITGFS**DDVPMVIA** |
| Monkey | | GADASAMQAARSTDVA | AVVVPILFLILLSLGVGFAILYT | KHRRLQSSFTAFANSHYSSRLGSAIFSSGDDLGEDDEDAPMITGFSDDVPMVIA |
| Pig | | GRDVSAIQATRSTDVA | AVVVPILFLILLSLGIGFAILYT | KHRRLQNSFTAFANSHYSSRLGSAIFSSGDDLGEDDEDAPMITGFSDDVPMVIA |
| Rat | | GGDATVVQTARSTDVA | AVVVPILFLILLSLGVGFAVLYT | KHRRLQSSFTAFANSHYSSRLGSAIFSSGDDLGEDDEDAPMITGFSDDVPMVIA |
| Mouse | | GADAAVIQAARSTDVA | AVVVPILFLILLSLGVGFAILYT | KHRRLQSSFSAFANSHYSSRLGSAIFSSGDDLGEDDEDAPMITGFSDDVPMVIA |
| Dog | | GGGASAFQAARSTDVA | AVVVPILFLILLSLGVGFAILYT | KHRRLQSSFTAFANSHYSSRLGSAIFSSGDDLGEDDEDAPMITGFSDDVPMVIA |
| Cat | | GAGASSARAARSTDVA | AVVVPILFLILLSLAVGFAVLYT | KHRRLRSSFTAFANSHYSSRLGSAIFSSGDDLGEDDEDAPMITGFSDDVPMVIA |
| Horse | | GGDASAFQAARSTDVA | AVVVPILFLILLSLGVGFAILYT | KHRRLQSSFTAFANSHYSSRLGSAIFSSGDDLGEDDEDAPMITGFSDDVPMVIA |
| Cow | | GGDASTIQAARSTDVA | AVVVPILFLILLSLGVGFAILYT | KHRRLQNSFTAFANSHYSSRLGSAIFSSGDDLGEDDEDAPMITGFSDDVPMVIA |
| Frog | | --DPLYSKNVQSTDVA | AIVVPILFLLLVATGFGFVILYT | RHRRLQNSFTAFANSHYSSRLGAAIFSSGDDIG-DDDDAPMITGFSDDVPMVIA |
| Zebrafish | | GQNDAASQSGKSEDMA | AIVVPVLFLLLVGVCGGLVVLYL | RHRRLQHSFTAFANSHYNSRLGSAIFSSGDELGDDDEDAPMISGFSDDVPMVIA |
| Parasitoid | | SSLAPWPATINSTNML | SIAIPICLLI-VALGVGLAYFFV | RHRRLSNSFTQFANSHYDTRRGQATFPGTTDAL-DEEDSPVIRGFSDDEPLVIA |
| Honey bee | | AEVSSLPVTINTSNIL | SFAIPICLLII-ALGSALAYFVV | RHRRLTNSFTQFANSHYDTRRGQATFPGTIDGL-EEEDSPVIRGFSDDEPLVIA |
| Termite | | MPVGSWKAVLSPRNIV | SVVVPVGLVMIALCGA-LAFFVL | RHRRLQRSFSSFANSHYDTRSGAATFSGGDGL--DEEDSPVIRGFSDDEPLVIA |
| Plant bug | | VPLGAWASVMTPTRVM | GIFVPICLVLIIVAGA-FVIFIV | RHRRLQNNFVTFANSHYSTRSGAATFTT-DGL--DDEDSPVIRGFSDDEPLVVA |
| | | | | **FANSHY** **DDLGEDDED** **DDVPMVIA** |

#### 6.b Supplemental Figure S6. Transmembrane and cytoplasmic domains

**a.** The cytosolic tail of SORLA contains different motifs that recognize molecular adaptors responsible of the intracellular localization of the receptor, incl sites for binding to Retromer (motif: FANSHY), PACS1 and AP1/AP2 (acidic motif: DDLGEDDED), and GGA (motif: DDVPMVIA).

**b.** Schematic of the intracellular trafficking pathways in which different adaptor proteins determine if SORLA goes into the secretory, endocytic, endosome delivery, or endosome recycling pathways.

**c.** Alignment of the human SORLA residues 2122-2214 with the corresponding regions of SORLA from different species including mammals, fish and insects. Residues 2138-2160 correspond to the transmembrane segment (yellow box) forming an $\alpha$-helical structure (lower panel). The three motifs in the cytoplasmic tail known to interact with adaptor proteins are shown below the alignment in bold letters.

#### 6.c *SORL1* variants in transmembrane and tail-domains

The 16 residues that make up the stalk and transmembrane, represent a small fraction of the entire SORLA protein. The ADES-ADSP dataset includes a variant that maps to the stalk region: the relatively prevalent p.T2134M variant which was observed in 22 cases and 13 controls [40], associating with a 1.7-fold increased AD risk (OR = 1.72 95% CI: 0.87 - 3.42 p=0.08), however, the much larger GWAS study does not support a pathogenic role of this variant (OR=1.14; p=0.4904) [224]. Functional testing indicated that mutated receptor was unable to protect APP from cleavage into Aβ by γ-secretase when compared to non-mutated SORLA, likely because the formation of a complex with APP was perturbed [225]. Also, lower levels of SORLA located to the cell surface of transfected cells, suggesting impaired trafficking properties of mutant SORLA.

Little is known about how the 23-residue TM region affects SORLA function, making it difficult to assess whether a variant will provide AD risk. But as pathogenic variants in the TM of LDLR have been reported to associate with FH [198], some variants in the SORLA TM region could also be speculated to be pathogenic.

Position 3 in the cytoplasmic tail, very close to the TM region, is part of a highly conserved sequence rich in positively charged amino acids ($^{2161}$KHRR$^{2164}$). These four residues are suggested to constitute a nuclear localization motif that is responsible for the translocation of the cytoplasmic tail of SORLA when liberated from the membrane after γ-secretase cleavage and with a role in signaling processes [226]. The ADES-ADSP dataset includes two variants at this position: p.R2163W observed in one case and p.R2163Q observed in one control. The dataset further includes several variants within the conserved $^{2172}$FANSHY motif, which may lead to a perturbed interaction with retromer and perturbed engagement in endosomal recycling. The ADES-ADSP dataset includes variants p.A2173T (conservation: 40/40; "*likely pathogenic*") observed in an AD case, p.S2175R (resulting from two independent genetic changes with different allele frequencies; conservation 40/40) that was observed in 11 cases and 10 controls, such that pathogenicity of this substitution is questionable. The dataset further includes the p.H2176R substitution, observed in one AD case. Together, the assessment of pathogenicity-levels for substitutions in the FANSHY motif require additional testing of variant effects. Finally, the ADES-ADSP dataset includes 3 variants that substitute key residues of the acidic/DXXXLL and gga/DXXVI: p.D2190N observed in an AD case and in a control, p.E2194K in two AD cases and a control and p.D2207G in an AD case. While such very conserved variants are likely linked with SORLA dysfunction, we observed them both in cases and controls such that more evidence is necessary to estimate their effects.

### 7 Summary of domain position prioritization for AD risk

Prioritization of positions likely to harbor pathogenic mutations based on either domain sequence conservation or because the disease-mutation domain-mapping analysis show enrichment for disease mutations for domains in other proteins.

| | | | | |
|---|---|---|---|---|
| **SORLA motifs** | **dom pos** | **SORLA residue/aa no.** | **description** | **priority** |
| RGD (in ProP) | | $R^{63}$, $G^{64}$, $D^{65}$ | | moderate |
| RRKR (in ProP) | | $R^{78}$, $R^{79}$, $K^{80}$, $R^{81}$ | | moderate |
| **VPS10p** | | | | |
| Hydrophobic; A-strand | 5, 6 | $V^{106}$, $V^{107}$,<br>$F^{167}$, $Y^{168}$,<br>$L^{209}$, $L^{210}$,<br>NA, $F^{251}$,<br>$F^{300}$, NA,<br>$Y^{349}$, $Y^{350}$,<br>$F^{414}$, NA,<br>$L^{495}$, NA,<br>$Y^{539}$, $Y^{540}$,<br>$V^{583}$, $Y^{584}$ | Conservative substitutions likely tolerated | moderate |
| Hydrophobic; B-strand | 19, 20, 21 | $I^{117}$, $V^{118}$, $A^{119}$,<br>$Y^{177}$, $I^{178}$, $F^{179}$,<br>$L^{218}$, $L^{219}$, $L^{220}$,<br>$I^{264}$, $Y^{265}$, $I^{266}$,<br>$Y^{306}$, $M^{307}$, $F^{308}$,<br>$V^{359}$, $F^{360}$, $V^{361}$,<br>$Y^{424}$, $I^{425}$, $A^{426}$,<br>$I^{504}$, $I^{505}$, $A^{506}$,<br>$I^{548}$, $I^{549}$, NA,<br>$V^{596}$, $F^{597}$, NA | Conservative substitutions likely tolerated | moderate |
| Hydrophobic; C-strand | 39, 40, 41 | $V^{135}$, $Y^{136}$, $V^{137}$,<br>$L^{187}$, $W^{188}$, $I^{189}$,<br>$L^{231}$, $W^{232}$, NA,<br>$V^{277}$, $F^{278}$, NA,<br>$L^{326}$, $W^{327}$, $V^{328}$,<br>$L^{372}$, $Y^{373}$, $I^{374}$,<br>NA, $V^{441}$, $I^{442}$,<br>$V^{520}$, $Y^{521}$, $I^{522}$,<br>$L^{561}$, NA, $Y^{563}$,<br>$L^{613}$, NA, $V^{615}$ | Conservative substitutions likely tolerated | moderate |
| Asp-box | 42, 44, 46, 48 49 | $S^{138}$, $D^{140}$, $G^{142}$, $S^{144}$, $F^{145}$,<br>$T^{190}$, $D^{192}$, NA, $T^{196}$, NA<br>$S^{234}$, $D^{236}$, $G^{238}$, $T^{240}$, $W^{241}$,<br>$S^{280}$, $D^{282}$, NA, $S^{286}$, NA,<br>$S^{329}$, NA, NA, NA, NA,<br>$S^{375}$, NA, $G^{379}$, NA, $F^{382}$,<br>$T^{443}$, $D^{445}$, $G^{447}$, $T^{449}$, $W^{450}$,<br>$S^{523}$, NA, $G^{527}$, NA, $W^{530}$,<br>$S^{564}$, $N^{566}$, $G^{568}$, $T^{570}$, $W^{571}$ | Positions are likely most pathogenic when occupied by a 'conserved' residue | high |
| L2 cysteines | | $C^{467}$, $C^{473}$ | Likely disulfide | high |

| | | | | |
|---|---|---|---|---|
| L1 & L2 cys gained | L1 & L2 protrusions | 391 - 411<br>457 - 493 | Disturb disulfide | high |
| **10CC** | | | | |
| ONC (9C; Cys loss) | | $C^{625}$, $C^{643}$, $C^{660}$, $C^{675}$, $C^{677}$, $C^{684}$, $C^{699}$, $C^{716}$, $C^{736}$, $C^{752}$ | Conserved disulfides | high |
| ONC (11C; Cys gained) | random | | Disturb disulfides | high |
| **YWTD** | | | | |
| YWTD | 16, 17, 18, 19 | $Y^{803}$, $W^{804}$, $S^{805}$, $D^{806}$,<br>$Y^{847}$, $W^{848}$, NA, $D^{850}$,<br>$F^{891}$, $W^{892}$, $T^{893}$, $D^{894}$,<br>$Y^{934}$, $W^{935}$, $T^{936}$, $D^{937}$,<br>$Y^{974}$, $W^{975}$, NA, $D^{977}$ | No substitution tolerated | high |
| hydrophobic | 6, 8, 15 | $L^{793}$, $F^{795}$, $L^{802}$,<br>$L^{837}$, $F^{839}$, $L^{846}$,<br>$L^{881}$, $L^{883}$, $M^{890}$,<br>$I^{926}$, $V^{928}$, $I^{933}$,<br>$I^{966}$, $V^{968}$, $I^{973}$,<br>$M^{1007}$, $I^{1009}$, NA | Conservative substitutions likely tolerated | moderate |
| hydrophobic | 41, 42 | NA, $L^{782}$,<br>$V^{825}$, $I^{826}$,<br>$I^{869}$, $V^{870}$,<br>$L^{915}$, $V^{916}$,<br>$I^{956}$, $L^{957}$,<br>$I^{996}$, $L^{997}$ | Conservative substitutions likeluy tolerated | moderate |
| Partly conserved Pro | 3 | $P^{878}$, $P^{923}$, $P^{963}$ | Only when P is lost | high |
| Partly conserved Asp | 9 | $D^{794}$, $D^{929}$ | Only when D is lost | high |
| Highly conserved Ile | 27 | $I^{769}$, $I^{812}$, $I^{856}$, $I^{902}$, $I^{943}$, $I^{983}$ | Only when I is lost | high |
| Highly conserved Arg | 29 | $R^{771}$, $R^{814}$, NA, $R^{904}$, $R^{945}$, $R^{985}$ | Only when R is lost | high |
| Highly conserved G | 35 | $G^{777}$, $G^{819}$, $G^{863}$, $G^{909}$, $G^{950}$, $G^{991}$ | Only when G is lost | high |
| Highly conserved L | 47 | $L^{787}$, $L^{831}$, $L^{875}$, $V^{920}$, $L^{960}$, $L^{1001}$ | Only when L is lost | moderate |
| R | 38 | $R^{866}$, $R^{953}$ | Partly conserved R - but DMDM only when R is changed | high |
| SBIN | 4 and 20 in β4/β5/β6 blades | $R^{879}$, $W^{895}$, $N^{924}$, $Y^{964}$, $W^{978}$ | (NXI) Ligand binding? | moderate |
| Disulfide β2 | | $C^{801}$, $C^{816}$ | Likely disulfide | high |
| **EGF** | | | | |
| ONC (7C; Cys loss) | | $C^{1021}$, $C^{1026}$, $C^{1030}$, $C^{1040}$, $C^{1042}$, $C^{1058}$, $C^{1060}$, $C^{1071}$ | Conserved disulfides | high |
| ONC (9C; Cys gain) | random | | Disturb disulfides | high |
| **CR** | | | | |
| CaCa (D,D,D,E) | 37, 41, 47, 48 | $D^{1098}$, $D^{1102}$, $D^{1108}$, $E^{1109}$,<br>$D^{1139}$, $D^{1143}$, $D^{1149}$, $E^{1150}$,<br>$D^{1178}$, $D^{1182}$, $D^{1188}$, $E^{1189}$, | Calcium cage | high |

| | | | | |
|---|---|---|---|---|
| | | $D^{1219}$, $D^{1223}$, $D^{1229}$, $E^{1230}$,<br>$D^{1257}$, $D^{1261}$, $D^{1267}$, $E^{1268}$,<br>$D^{1297}$, $Q^{1301}$, $D^{1307}$, $E^{1308}$,<br>$D^{1345}$, $D^{1349}$, $D^{1355}$, $E^{1356}$,<br>$D^{1389}$, $D^{1393}$, $D^{1399}$, $E^{1400}$,<br>$D^{1439}$, $D^{1443}$, $D^{1449}$, $E^{1450}$,<br>$D^{1492}$, $D^{1496}$, $D^{1502}$, $E^{1503}$,<br>$D^{1535}$, $D^{1539}$, $D^{1545}$, $E^{1546}$ | | |
| ONC (5C; Cys loss) | 15, 23, 29, 36, 42, 55 | $C^{1078}$, $C^{1085}$, $C^{1090}$, $C^{1097}$, $C^{1103}$, $C^{1112}$,<br>$C^{1117}$, $C^{1125}$, $C^{1131}$, $C^{1138}$, $C^{1144}$, $C^{1153}$,<br>$C^{1158}$, $C^{1165}$, $C^{1170}$, $C^{1177}$, $C^{1183}$, $C^{1192}$,<br>$C^{1199}$, $C^{1206}$, $C^{1211}$, $C^{1218}$, $C^{1224}$, $C^{1235}$,<br>$C^{1239}$, $C^{1244}$, $C^{1249}$, $C^{1256}$, $C^{1262}$, $C^{1271}$,<br>$C^{1275}$, $C^{1283}$, $C^{1289}$, $C^{1296}$, $C^{1302}$, $C^{1315}$,<br>$C^{1325}$, $C^{1332}$, $C^{1337}$, $C^{1344}$, $C^{1350}$, $C^{1359}$,<br>$C^{1368}$, $C^{1376}$, $C^{1381}$, $C^{1388}$, $C^{1394}$, $C^{1403}$,<br>$C^{1419}$, $C^{1426}$, $C^{1431}$, $C^{1438}$, $C^{1444}$, $C^{1453}$,<br>$C^{1471}$, $C^{1478}$, $C^{1484}$, $C^{1491}$, $C^{1497}$, $C^{1506}$,<br>$C^{1514}$, $C^{1521}$, $C^{1527}$, $C^{1534}$, $C^{1540}$, $C^{1549}$ | Conserved disulfides | high |
| ONC (7C; Cys gain) | random | | Disturb disulfides | high |
| Asx-turn (D) | 44 | $D^{1105}$, $D^{1146}$, $D^{1185}$, $D^{1226}$, $D^{1264}$, $D^{1304}$,<br>$D^{1352}$, $D^{1396}$, $D^{1446}$, $D^{1499}$, $D^{1542}$ | No substitutions tolerated | high |
| Asx-turn (S) | 46 | $S^{1107}$, $S^{1148}$, $S^{1187}$, $S^{1228}$, $S^{1266}$, $S^{1306}$,<br>$S^{1354}$, $S^{1398}$, $S^{1448}$, NA, $S^{1544}$ | Only when S is lost | moderate |
| Fingerprints | 34, 39 | $W^{1095}$, $D^{1100}$,<br>$Y^{1136}$, $E^{1141}$,<br>$W^{1175}$, $D^{1180}$,<br>$W^{1216}$, $D^{1221}$,<br>$K^{1254}$, $L^{1259}$,<br>$M^{1294}$, $I^{1299}$,<br>$W^{1342}$, $M^{1347}$,<br>$W^{1386}$, $E^{1391}$,<br>$W^{1436}$, $Y^{1441}$,<br>$K^{1489}$, $H^{1494}$,<br>$E^{1532}$, $F^{1537}$ | | moderate |
| Phe-Ile hyd core | 21, 30 | $Y^{1083}$, $I^{1091}$,<br>$F^{1123}$, $I^{1132}$,<br>$Y^{1163}$, $I^{1171}$,<br>$F^{1204}$, $I^{1212}$,<br>$F^{1242}$, $I^{1250}$,<br>$F^{1281}$, $L^{1290}$,<br>$F^{1330}$, $I^{1338}$,<br>$F^{1374}$, $I^{1382}$,<br>$Y^{1424}$, $V^{1432}$,<br>$F^{1476}$, $I^{1485}$,<br>$F^{1519}$, $I^{1528}$ | Strong conservation, but only few variants in the DMDM analysis | moderate |
| Partly conserved glycines | 27, 38 | $G^{1088}$, NA,<br>$G^{1129}$, NA,<br>$G^{1168}$, $G^{1179}$,<br>$G^{1209}$, $G^{1220}$,<br>$G^{1247}$, $G^{1258}$,<br>NA, $G^{1298}$,<br>$G^{1335}$, $G^{1346}$,<br>$G^{1379}$, NA,<br>$G^{1429}$, $G^{1440}$,<br>NA, $G^{1493}$,<br>NA, $G^{1536}$ | Only partial conservation.<br>Few variants in DMDM analysis | moderate |
| **3Fn** | | | | |

| | | | | |
|---|---|---|---|---|
| Partly conserved prolines | 6, 7 | NA, NA, $P^{1654}$, NA, $P^{1749}$, $P^{1750}$, $P^{1843}$, $P^{1844}$, $P^{1934}$, $P^{1935}$, NA, $P^{2027}$ | Only when P is lost | high |
| Hydrophobic | 11, 13 | $L^{1561}$, $W^{1563}$,<br>$L^{1659}$, $L^{1661}$,<br>$I^{1753}$, $I^{1755}$,<br>$L^{1848}$, $A^{1850}$,<br>$L^{1938}$, $V^{1940}$,<br>$L^{2030}$, $I^{2032}$ | Conservative substitutions likely tolerated | moderate |
| W; B-strand | 25 | $W^{1575}$, $W^{1673}$, ($L^{1767}$), $W^{1862}$, $W^{1952}$, $W^{2043}$ | No substitutions tolerated | high |
| Hydrophobics in B-strand | 21, 23 | $V^{1571}$, $L^{1573}$, $I^{1669}$, NA, $L^{1763}$, $F^{1765}$, $V^{1858}$, $C^{1860}$, $V^{1948}$, $I^{1950}$, $V^{2039}$, $L^{2041}$ | Conservative substitutions likely tolerated | moderate |
| Partly conserved prolines | 28 | $P^{1578}$, $P^{1676}$, NA, $P^{1865}$, $P^{1955}$, NA | Only when P is lost | high |
| Y; C-strand | 41 | $Y^{1588}$, $Y^{1686}$, $Y^{1778}$, $Y^{1870}$, $Y^{1965}$, $Y^{2059}$ | No substitutions tolerated | high |
| Hydrophobics in C-strand | 43, 45 | $V^{1590}$, $Y^{1592}$, $V^{1688}$, $Y^{1690}$, $V^{1780}$, $L^{1782}$, $I^{1872}$, $Y^{1874}$, $V^{1967}$, $V^{1969}$, $I^{2061}$, $M^{2063}$ | Conservative (hydrophobic) substitutions tolerated | moderate |
| L; EF-loop (tyrosine corner) | 77 | $L^{1617}$, $L^{1713}$, $L^{1810}$, NA, $L^{1996}$, $L^{2087}$ | No substitutions tolerated | high |
| Partly conserved glycines | 36, 96 | $G^{1681}$, $G^{1732}$, $G^{1917}$ | Only when G is lost | high |
| Partly conserved glycines | 94 | $G^{1730}$, $G^{1827}$, $G^{2104}$ | Only when G is lost | moderate |
| Y; F-strand (tyrosine-corner) | 83 | $Y^{1623}$, $Y^{1719}$, $Y^{1816}$, $Y^{1905}$, $Y^{2002}$, $Y^{2093}$ | No substitutions tolerated | high |
| Hydrophobics in F-strand | 85, 87, 89 | $V^{1625}$, $V^{1627}$, $V^{1629}$, $V^{1721}$, $V^{1723}$, $A^{1725}$, $I^{1818}$, $A^{1820}$, $A^{1822}$, $F^{1907}$, $V^{1909}$, $V^{1911}$, $I^{2004}$, $V^{2006}$, $L^{2008}$, $F^{2095}$, $V^{2097}$, $A^{2099}$ | Conservative (hydrophobic) substitutions tolerated | moderate |
| Hydrophobics in E-strand | 72, 74 | NA, $L^{1614}$,<br>NA, $I^{1710}$,<br>NA, $V^{1807}$,<br>$V^{1897}$, $V^{1899}$,<br>$Y^{1991}$, $L^{1993}$,<br>$F^{2082}$, $I^{2084}$ | | moderate |
| Partly conserved prolines | 79 | $P^{1619}$, $P^{1998}$ | Only when P is lost | high |
| Trp-ladders | | $R^{1593}$, $W^{1600}$, $K^{1626}$, $H^{1636}$, $E^{1690}$, $W^{1698}$, $R^{1722}$, $W^{1734}$ | | moderate |
| Arg hotspot | 88 | $R^{1910}$ | Based on Arg substitutions in DMDM pathogenic | high |
| Disulfide, 1st | | $C^{1586}$, $C^{1631}$ | Disulfide | high |
| Disulfide, 6th | | $C^{2101}$, $C^{2108}$ | Disulfide | high |
| **Tail** | | | | |
| FANSHY | | $F^{2172}$, $A^{2173}$, $N^{2174}$, $S^{2175}$, $H^{2176}$, $Y^{2177}$ | | moderate |
| Acidic | | $D^{2190}$, $D^{2191}$, $L^{2192}$, $G^{2193}$, $E^{2194}$, $D^{2195}$, $D^{2196}$, $E^{2197}$, $D^{2198}$ | | moderate |

| | | | | |
|---|---|---|---|---|
| GGA | | $D^{2207}$, $D^{2208}$, $V^{2209}$, $P^{2210}$, $M^{2211}$, $V^{2212}$, $I^{2213}$, $A^{2214}$ | | moderate |

### 8 SORL1 sequences for alignment: species conservation

A total of 40 different SORLA protein sequences are included in the species alignment (**Supplemental Information 9**).

This alignment allows look-up if genetic variants that make substitutions in the human protein leads to inclusion of amino acids present in SORLA from other species – in which case such a variant is more likely to be tolerated than if the position is strictly conserved across all species.

| | |
|---|---|
| Homo sapiens (human): | NP_003096.2/Q92673-1 |
| Macaca mulatta (rhesus macaque): | XP_014971461.2/H9ZCQ1-1 |
| Pan troglodytes (chimpanzee): | XP_016777658.1/H2Q4Z6-1 |
| Sus scrofa (pig): | XP_020918668.1/I3L8K1-1 |
| Capra hircus (goat): | XP_017915127.1/A0A452FGN5-1 |
| Ovis aries (sheep): | XP_027835138.1/W5QA68-1 |
| Equus caballus (horse): | XP_023500779.1/F7CG82-1 |
| Bos taurus (cow): | NP_001179686.1/E1BPZ1-1 |
| Loxodonta africana (elephant): | XP_003418317.2/G3T328-1 |
| Canis lupus familiaris (dog): | XP_536545.2/ E2R5F5-1 |
| Canis lupus dingo (wolf): | XP_025320979.1 |
| Vulpes vulpes (fox): | XP_025854818.1/A0A3Q7SF13-1 |
| Ursus arctos horribilis (bear): | XP_026361326.1 |
| Felis catus (cat): | XP_023094970.1/M3WIG3-1 |
| Panthera pardus (leopard): | XP_019324393.1 |
| Oryctolagus cuniculus (rabbit): | NP_001076133.1/Q95209-1 |
| Rattus norvegicus (rat): | NP_445971.1/P0DSP1-1 |
| Mus musculus (mouse): | NP_035566.2/O88307-1 |
| Ornithorhynchus anatinus (platypus): | XP_028930988.1/F7D9P5-1 |
| Gallus gallus (chicken): | NP_001292089.1/E1BUD4-1 |
| Anas platyrhynchos (duck): | XP_027299930.1 |
| Columba livia (pigeon): | XP_021152551.1/A0A2I0LS76-1 |
| Haliaeetus leucocephalus (eagle): | XP_010579730.1 |
| Falco cherrug (falcon): | XP_027662563.1 |
| Aptennodytes forsteri (penguin): | XP_009279556.1 |
| Danio rerio (zebra fish): | XP_005157607.1/X1WHE3-1 |
| Salmo salar (salmon): | XP_014017958.1/A0A1S3NRA8-1 |
| Amphiprion ocellaris (clown fish): | XP_023117170.1 |
| Pygocentrus nattereri (piranha): | XP_017568534.1 |
| Esox lucius (pike): | XP_010891973.1 |
| Hippocampus comes (seahorse): | XP_019716899.1 |
| Rhincodon typus (whale shark): | XP_020382540.1 |
| Xenopus tropicalis (frog): | XP_031762503.1/A0A1L8FLB9-1 |
| Pelodiscus sinensis (turtle): | XP_025045904.1/ K7F2T1-1 |
| Alligator mississippiensis (alligator): | XP_014449241.1/ A0A151NQK7-1 |
| Crocodylus porosus (crocodile): | XP_019402601.1 |
| Gekko japonicus (gecko): | XP_015277064.1 |
| Python bivittatus (python): | XP_007431813.2 |
| Anolis carolinensis (anole): | XP_016850209.1/ H9G6H6-1 |
| Podarcis muralis (lizard): | XP_028563124.1 |

### 9 SORL1 sequence alignment

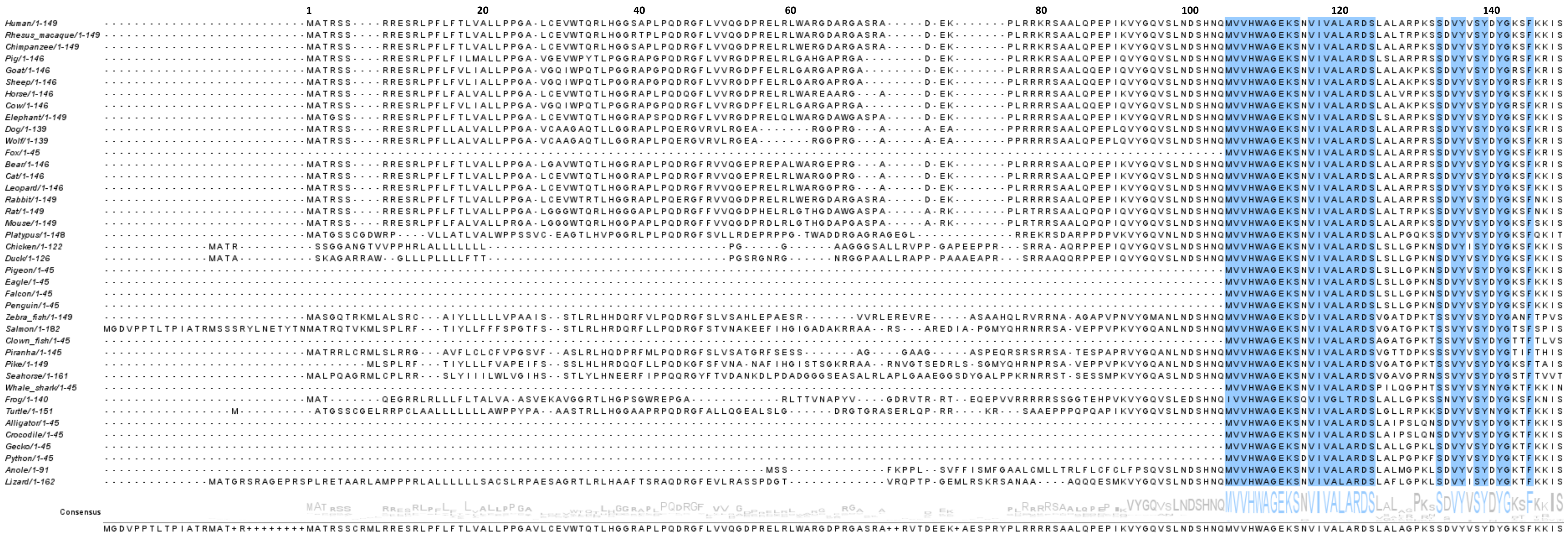
1
20
40
60
80
100
120
140
Human/1-149
Rhesus_macaque/1-149
Chimpanzee/1-149
Pig/1-146
Goat/1-146
Sheep/1-146
Horse/1-146
Cow/1-146
Elephant/1-149
Dog/1-139
Wolf/1-139
Fox/1-45
Bear/1-146
Cat/1-146
Leopard/1-146
Rabbit/1-149
Rat/1-149
Mouse/1-149
Platypus/1-148
Chicken/1-122
Duck/1-126
Pigeon/1-45
Eagle/1-45
Falcon/1-45
Penguin/1-45
Zebra_fish/1-149
Salmon/1-182
Clown_fish/1-45
Piranha/1-145
Pike/1-149
Seahorse/1-161
Whale_shark/1-45
Frog/1-140
Turtle/1-151
Alligator/1-45
Crocodile/1-45
Gecko/1-45
Python/1-45
Anole/1-91
Lizard/1-162
Consensus

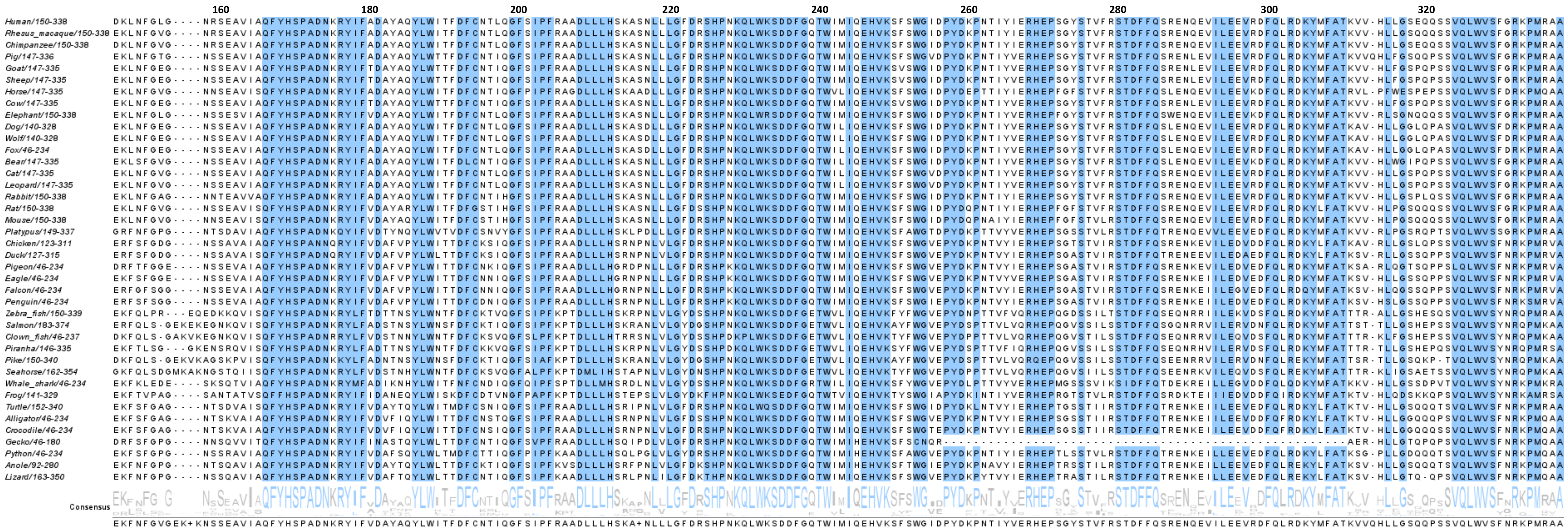
160
180
200
220
240
260
280
300
320
Human/150-338
Rhesus_macaque/150-338
Chimpanzee/150-338
Pig/147-336
Goat/147-335
Sheep/147-335
Horse/147-335
Cow/147-335
Elephant/150-338
Dog/140-328
Wolf/140-328
Fox/46-234
Bear/147-335
Cat/147-335
Leopard/147-335
Rabbit/150-338
Rat/150-338
Mouse/150-338
Platypus/149-337
Chicken/123-311
Duck/127-315
Pigeon/46-234
Eagle/46-234
Falcon/46-234
Penguin/46-234
Zebra_fish/150-339
Salmon/183-374
Clown_fish/46-237
Piranha/146-335
Pike/150-340
Seahorse/162-354
Whale_shark/46-234
Frog/141-329
Turtle/152-340
Alligator/46-234
Crocodile/46-234
Gecko/46-180
Python/46-234
Anole/92-280
Lizard/163-350
Consensus

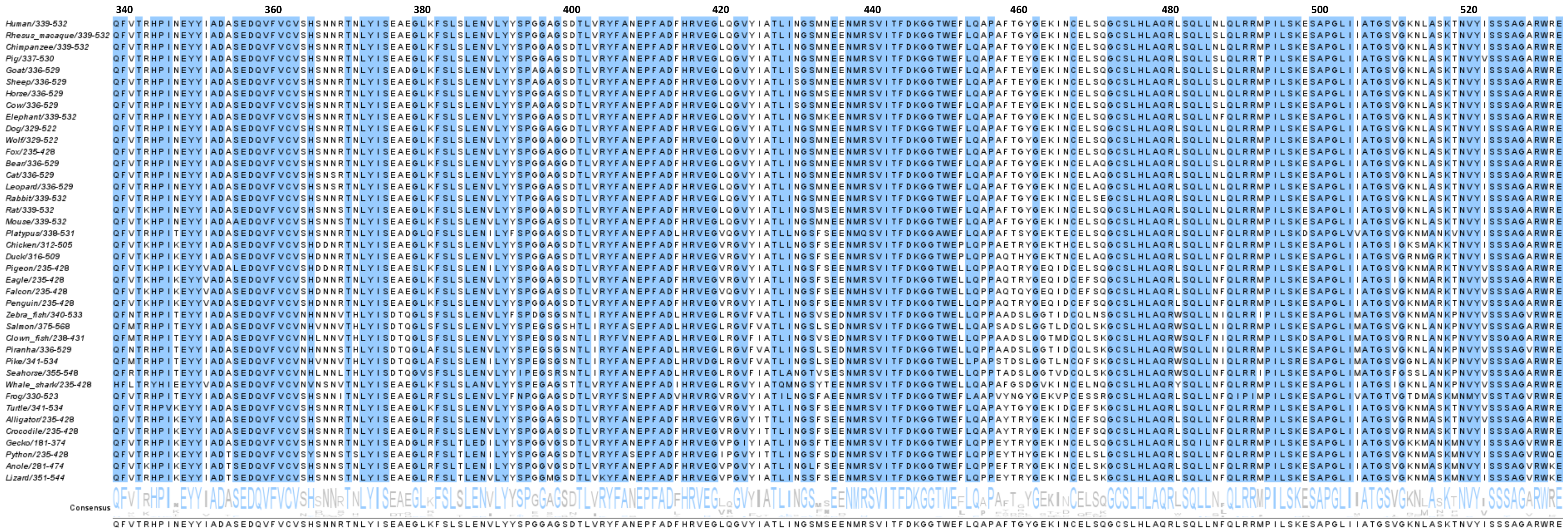
340
360
380
400
420
440
460
480
500
520
Human/339-532
Rhesus_macaque/339-532
Chimpanzee/339-532
Pig/337-530
Goat/336-529
Sheep/336-529
Horse/336-529
Cow/336-529
Elephant/339-532
Dog/329-522
Wolf/329-522
Fox/235-428
Bear/336-529
Cat/336-529
Leopard/336-529
Rabbit/339-532
Rat/339-532
Mouse/339-532
Platypus/338-531
Chicken/312-505
Duck/316-509
Pigeon/235-428
Eagle/235-428
Falcon/235-428
Penguin/235-428
Zebra_fish/340-533
Salmon/375-568
Clown_fish/238-431
Piranha/336-529
Pike/341-534
Seahorse/355-548
Whale_shark/235-428
Frog/330-523
Turtle/341-534
Alligator/235-428
Crocodile/235-428
Gecko/181-374
Python/235-428
Anole/281-474
Lizard/351-544
Consensus

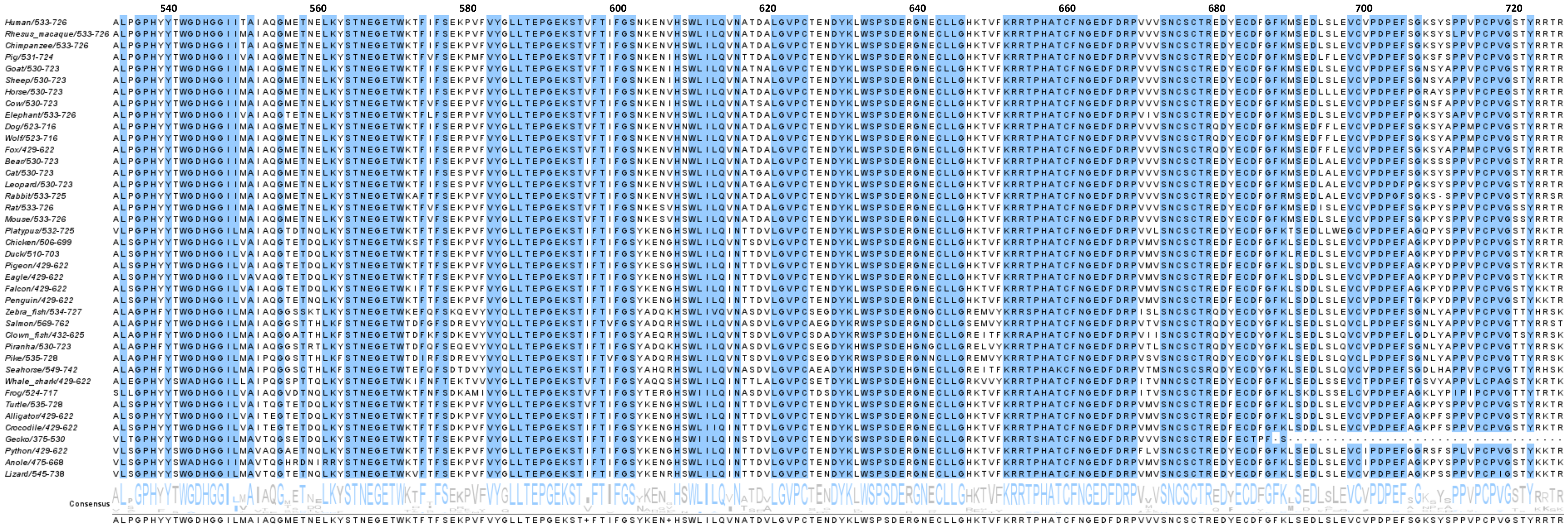

540
560
580
600
620
640
660
680
700
720
Human/533-726
Rhesus_macaque/533-726
Chimpanzee/533-726
Pig/531-724
Goat/530-723
Sheep/530-723
Horse/530-723
Cow/530-723
Elephant/533-726
Dog/523-716
Wolf/523-716
Fox/429-622
Bear/530-723
Cat/530-723
Leopard/530-723
Rabbit/533-725
Rat/533-726
Mouse/533-726
Platypus/532-725
Chicken/506-699
Duck/510-703
Pigeon/429-622
Eagle/429-622
Falcon/429-622
Penguin/429-622
Zebra_fish/534-727
Salmon/569-762
Clown_fish/432-625
Piranha/530-723
Pike/535-728
Seahorse/549-742
Whale_shark/429-622
Frog/524-717
Turtle/535-728
Alligator/429-622
Crocodile/429-622
Gecko/375-530
Python/429-622
Anole/475-668
Lizard/545-738
Consensus
ALPGPHYYTWGDHGGILMAIAQGMETNELKYSTNEGETWKTFTFSEKPVFVYGLLTEPGEKST+FTIFGSYKEN+HSWLILQVNATDVLGVPCTENDYKLWSPSDERGNECLLGHKTVFKRRTPHATCFNGEDFDRPVVVSNCSCTREDYECDFGFKLSEDLSLEVCVPDPEFSGKSYSPPVPCPVGSTYRRTR

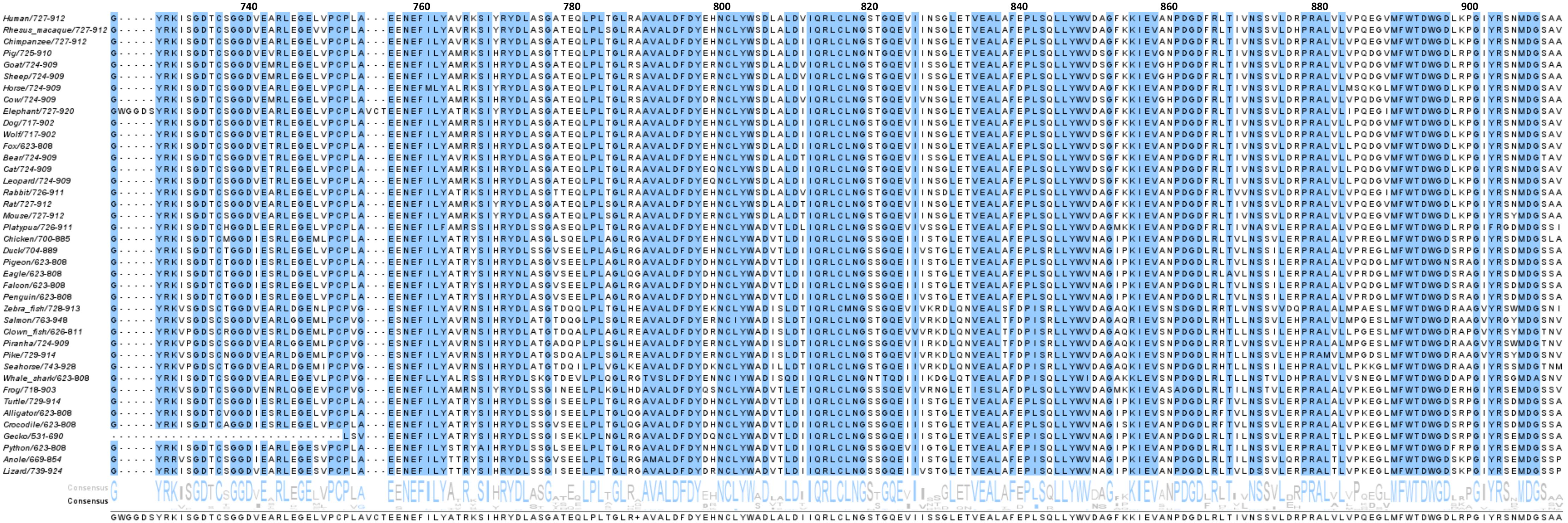
740
760
780
800
820
840
860
880
900
Human/727-912
Rhesus_macaque/727-912
Chimpanzee/727-912
Pig/725-910
Goat/724-909
Sheep/724-909
Horse/724-909
Cow/724-909
Elephant/727-920
Dog/717-902
Wolf/717-902
Fox/623-808
Bear/724-909
Cat/724-909
Leopard/724-909
Rabbit/726-911
Rat/727-912
Mouse/727-912
Platypus/726-911
Chicken/700-885
Duck/704-889
Pigeon/623-808
Eagle/623-808
Falcon/623-808
Penguin/623-808
Zebra_fish/728-913
Salmon/763-948
Clown_fish/626-811
Piranha/724-909
Pike/729-914
Seahorse/743-928
Whale_shark/623-808
Frog/718-903
Turtle/729-914
Alligator/623-808
Crocodile/623-808
Gecko/531-690
Python/623-808
Anole/669-854
Lizard/739-924
Consensus
Consensus

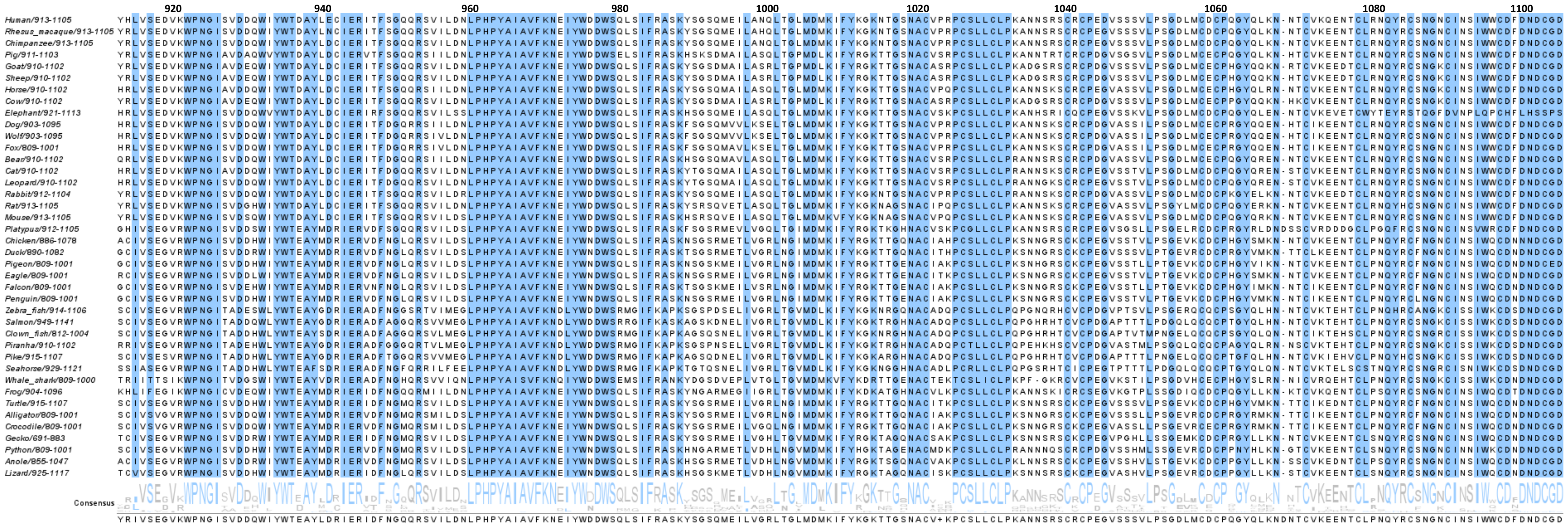
920
940
960
980
1000
1020
1040
1060
1080
1100
Human/913-1105
Rhesus_macaque/913-1105
Chimpanzee/913-1105
Pig/911-1103
Goat/910-1102
Sheep/910-1102
Horse/910-1102
Cow/910-1102
Elephant/921-1113
Dog/903-1095
Wolf/903-1095
Fox/809-1001
Bear/910-1102
Cat/910-1102
Leopard/910-1102
Rabbit/912-1104
Rat/913-1105
Mouse/913-1105
Platypus/912-1105
Chicken/886-1078
Duck/890-1082
Pigeon/809-1001
Eagle/809-1001
Falcon/809-1001
Penguin/809-1001
Zebra_fish/914-1106
Salmon/949-1141
Clown_fish/812-1004
Piranha/910-1102
Pike/915-1107
Seahorse/929-1121
Whale_shark/809-1000
Frog/904-1096
Turtle/915-1107
Alligator/809-1001
Crocodile/809-1001
Gecko/691-883
Python/809-1001
Anole/855-1047
Lizard/925-1117
Consensus

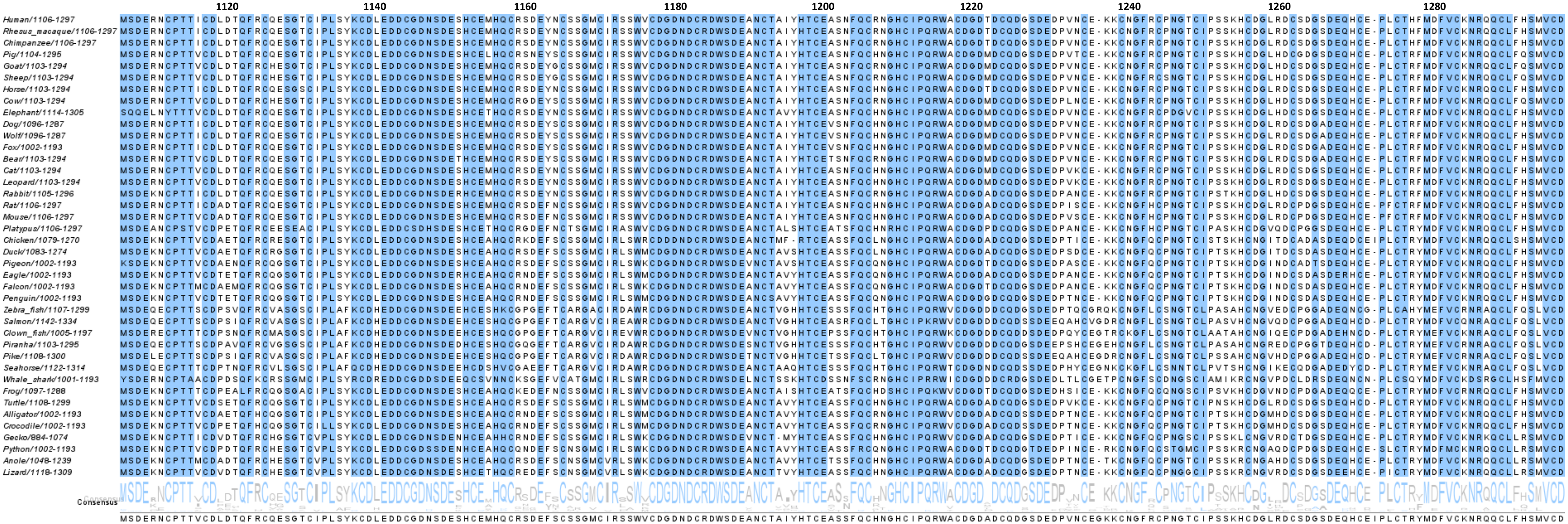
1120
1140
1160
1180
1200
1220
1240
1260
1280
Human/1106-1297
Rhesus_macaque/1106-1297
Chimpanzee/1106-1297
Pig/1104-1295
Goat/1103-1294
Sheep/1103-1294
Horse/1103-1294
Cow/1103-1294
Elephant/1114-1305
Dog/1096-1287
Wolf/1096-1287
Fox/1002-1193
Bear/1103-1294
Cat/1103-1294
Leopard/1103-1294
Rabbit/1105-1296
Rat/1106-1297
Mouse/1106-1297
Platypus/1106-1297
Chicken/1079-1270
Duck/1083-1274
Pigeon/1002-1193
Eagle/1002-1193
Falcon/1002-1193
Penguin/1002-1193
Zebra_fish/1107-1299
Salmon/1142-1334
Clown_fish/1005-1197
Piranha/1103-1295
Pike/1108-1300
Seahorse/1122-1314
Whale_shark/1001-1193
Frog/1097-1288
Turtle/1108-1299
Alligator/1002-1193
Crocodile/1002-1193
Gecko/884-1074
Python/1002-1193
Anole/1048-1239
Lizard/1118-1309
Consensus
MSDERNCPTTVCDLDTQFRCQESGTCIPLSYKCDLEDDCGDNSDESHCEMHQCRSDEFSCSSGMCIRSSWVCDGDNDCRDWSDEANCTAIYHTCEASSFQCHNGHCIPQRWACDGDADCQDGSDEDPVNCEGKKCNGFRCPNGTCIPSSKHCDGLRDCSDGSDEQHCEIPLCTRYMDFVCKNRQQCLFHSMVCD

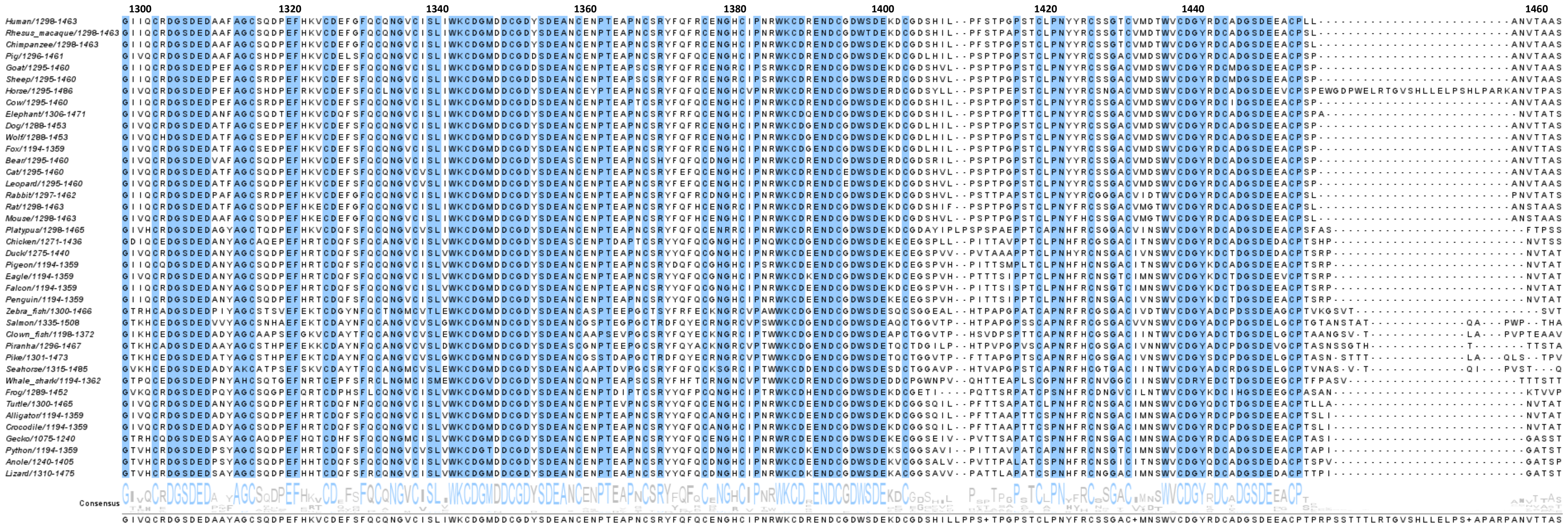
1300
1320
1340
1360
1380
1400
1420
1440
1460
Human/1298-1463
Rhesus_macaque/1298-1463
Chimpanzee/1298-1463
Pig/1296-1461
Goat/1295-1460
Sheep/1295-1460
Horse/1295-1486
Cow/1295-1460
Elephant/1306-1471
Dog/1288-1453
Wolf/1288-1453
Fox/1194-1359
Bear/1295-1460
Cat/1295-1460
Leopard/1295-1460
Rabbit/1297-1462
Rat/1298-1463
Mouse/1298-1463
Platypus/1298-1465
Chicken/1271-1436
Duck/1275-1440
Pigeon/1194-1359
Eagle/1194-1359
Falcon/1194-1359
Penguin/1194-1359
Zebra_fish/1300-1466
Salmon/1335-1508
Clown_fish/1198-1372
Piranha/1296-1467
Pike/1301-1473
Seahorse/1315-1485
Whale_shark/1194-1362
Frog/1289-1452
Turtle/1300-1465
Alligator/1194-1359
Crocodile/1194-1359
Gecko/1075-1240
Python/1194-1359
Anole/1240-1405
Lizard/1310-1475
Consensus

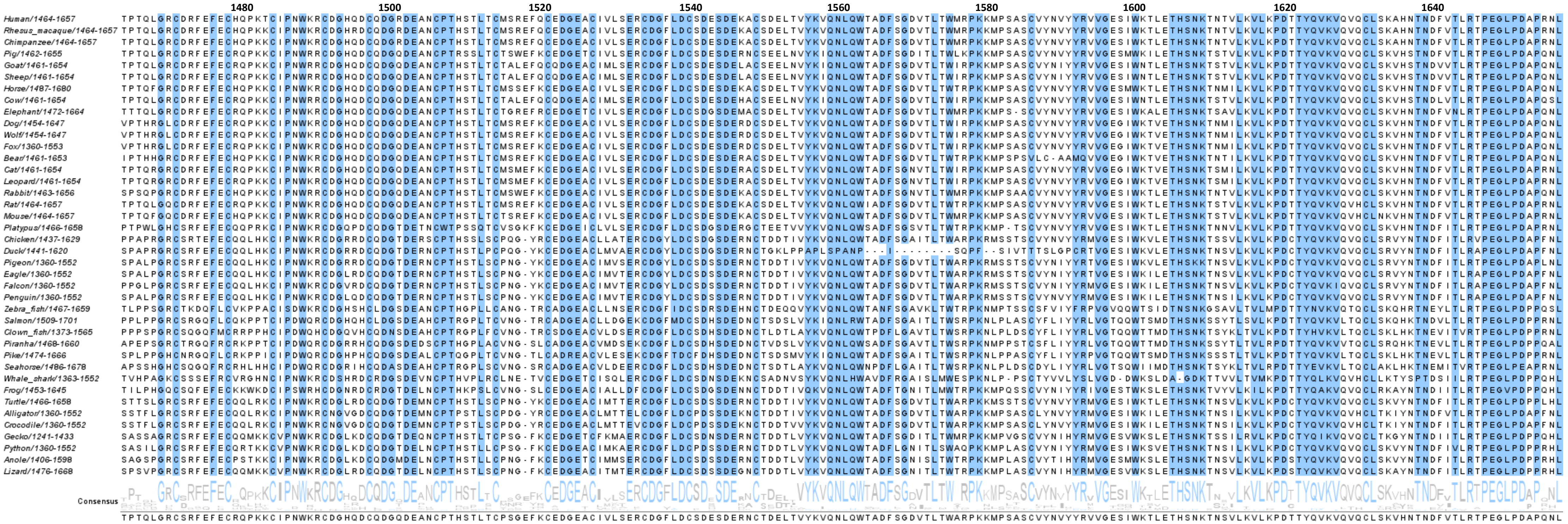
1480
1500
1520
1540
1560
1580
1600
1620
1640
Human/1464-1657
Rhesus_macaque/1464-1657
Chimpanzee/1464-1657
Pig/1462-1655
Goat/1461-1654
Sheep/1461-1654
Horse/1487-1680
Cow/1461-1654
Elephant/1472-1664
Dog/1454-1647
Wolf/1454-1647
Fox/1360-1553
Bear/1461-1653
Cat/1461-1654
Leopard/1461-1654
Rabbit/1463-1656
Rat/1464-1657
Mouse/1464-1657
Platypus/1466-1658
Chicken/1437-1629
Duck/1441-1620
Pigeon/1360-1552
Eagle/1360-1552
Falcon/1360-1552
Penguin/1360-1552
Zebra_fish/1467-1659
Salmon/1509-1701
Clown_fish/1373-1565
Piranha/1468-1660
Pike/1474-1666
Seahorse/1486-1678
Whale_shark/1363-1552
Frog/1453-1645
Turtle/1466-1658
Alligator/1360-1552
Crocodile/1360-1552
Gecko/1241-1433
Python/1360-1552
Anole/1406-1598
Lizard/1476-1668
Consensus

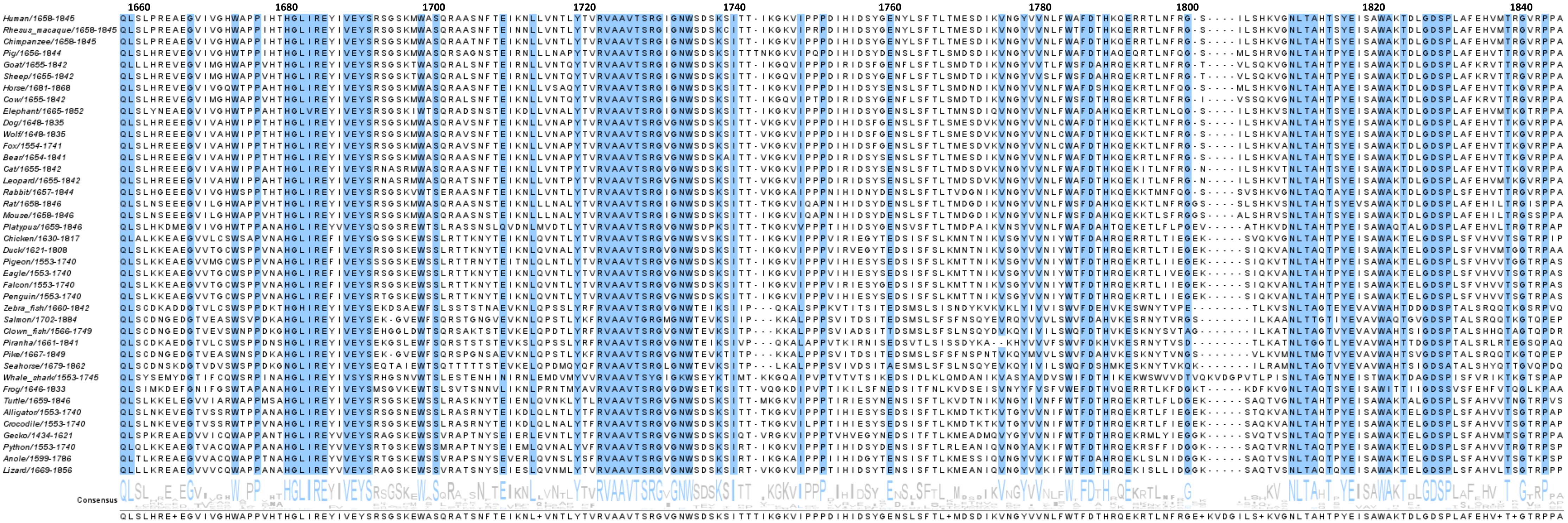
1660
1680
1700
1720
1740
1760
1780
1800
1820
1840
Human/1658-1845
Rhesus_macaque/1658-1845
Chimpanzee/1658-1845
Pig/1656-1844
Goat/1655-1842
Sheep/1655-1842
Horse/1681-1868
Cow/1655-1842
Elephant/1665-1852
Dog/1648-1835
Wolf/1648-1835
Fox/1554-1741
Bear/1654-1841
Cat/1655-1842
Leopard/1655-1842
Rabbit/1657-1844
Rat/1658-1846
Mouse/1658-1846
Platypus/1659-1846
Chicken/1630-1817
Duck/1621-1808
Pigeon/1553-1740
Eagle/1553-1740
Falcon/1553-1740
Penguin/1553-1740
Zebra_fish/1660-1842
Salmon/1702-1884
Clown_fish/1566-1749
Piranha/1661-1841
Pike/1667-1849
Seahorse/1679-1862
Whale_shark/1553-1745
Frog/1646-1833
Turtle/1659-1846
Alligator/1553-1740
Crocodile/1553-1740
Gecko/1434-1621
Python/1553-1740
Anole/1599-1786
Lizard/1669-1856
Consensus

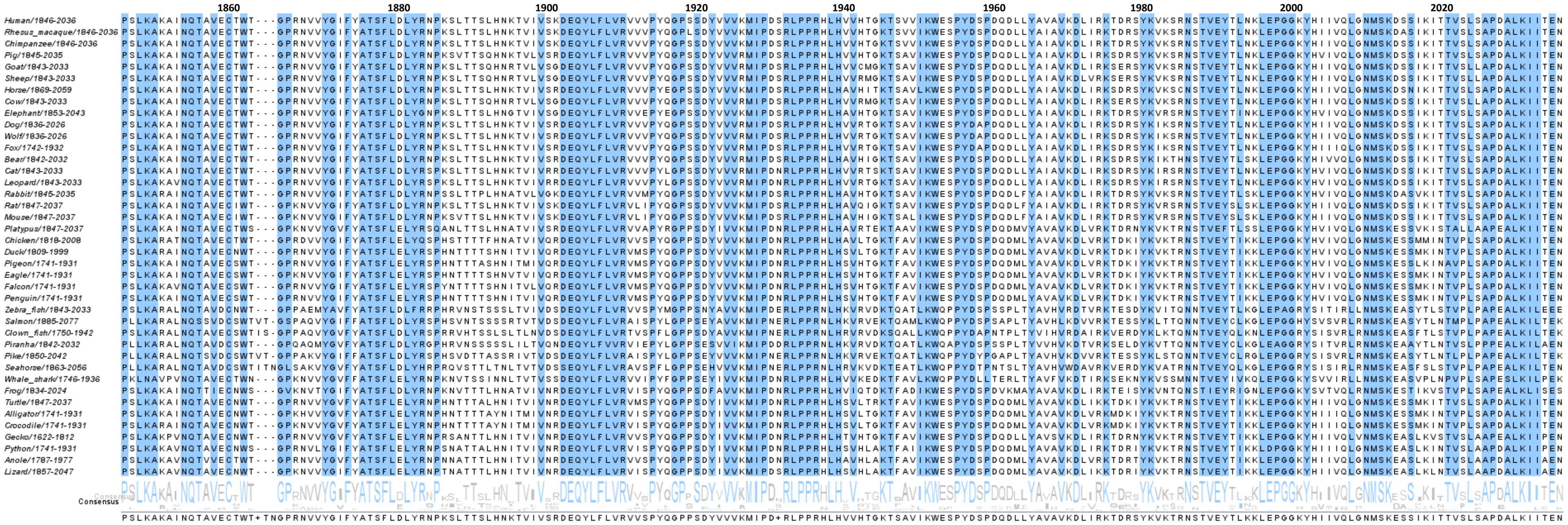
1860
1880
1900
1920
1940
1960
1980
2000
2020
Human/1846-2036
Rhesus_macaque/1846-2036
Chimpanzee/1846-2036
Pig/1845-2035
Goat/1843-2033
Sheep/1843-2033
Horse/1869-2059
Cow/1843-2033
Elephant/1853-2043
Dog/1836-2026
Wolf/1836-2026
Fox/1742-1932
Bear/1842-2032
Cat/1843-2033
Leopard/1843-2033
Rabbit/1845-2035
Rat/1847-2037
Mouse/1847-2037
Platypus/1847-2037
Chicken/1818-2008
Duck/1809-1999
Pigeon/1741-1931
Eagle/1741-1931
Falcon/1741-1931
Penguin/1741-1931
Zebra_fish/1843-2033
Salmon/1885-2077
Clown_fish/1750-1942
Piranha/1842-2032
Pike/1850-2042
Seahorse/1863-2056
Whale_shark/1746-1936
Frog/1834-2024
Turtle/1847-2037
Alligator/1741-1931
Crocodile/1741-1931
Gecko/1622-1812
Python/1741-1931
Anole/1787-1977
Lizard/1857-2047
Consensus
PSLKAKAINQTAVECTWT+TNGPRNVVYGIFYATSFLDLYRNPKSLTTSLHNKTVIVSRDEQYLFLVRVVSPYQGPPSDYVVVKMIPD+RLPPRHLHVVHTGKTSAVIKWESPYDSPDQDLLYAVAVKDLIRKTDRSYKVKTRNSTVEYTLKKLEPGGKYHIIVQLGNMSKESSIKITTVSLSAPDALKIITEN

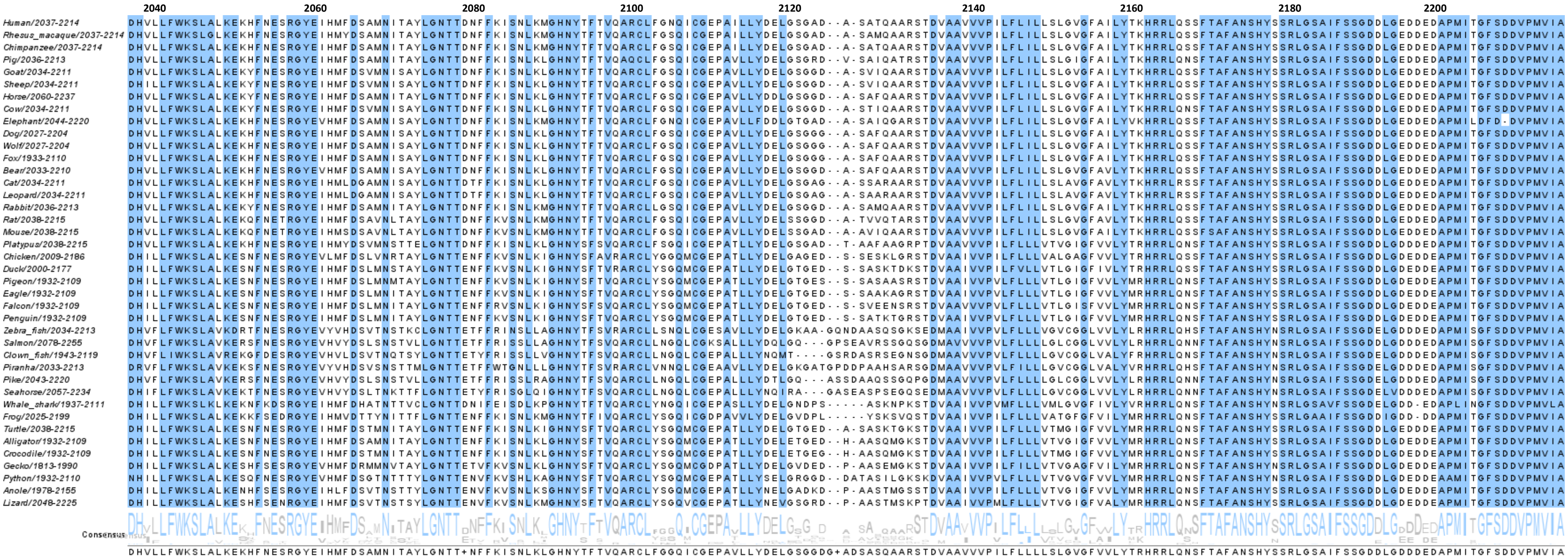
2040
2060
2080
2100
2120
2140
2160
2180
2200
Human/2037-2214
Rhesus_macaque/2037-2214
Chimpanzee/2037-2214
Pig/2036-2213
Goat/2034-2211
Sheep/2034-2211
Horse/2060-2237
Cow/2034-2211
Elephant/2044-2220
Dog/2027-2204
Wolf/2027-2204
Fox/1933-2110
Bear/2033-2210
Cat/2034-2211
Leopard/2034-2211
Rabbit/2036-2213
Rat/2038-2215
Mouse/2038-2215
Platypus/2038-2215
Chicken/2009-2186
Duck/2000-2177
Pigeon/1932-2109
Eagle/1932-2109
Falcon/1932-2109
Penguin/1932-2109
Zebra_fish/2034-2213
Salmon/2078-2255
Clown_fish/1943-2119
Piranha/2033-2213
Pike/2043-2220
Seahorse/2057-2234
Whale_shark/1937-2111
Frog/2025-2199
Turtle/2038-2215
Alligator/1932-2109
Crocodile/1932-2109
Gecko/1813-1990
Python/1932-2110
Anole/1978-2155
Lizard/2048-2225
Consensus

### 10 Phylogenetic tree for SORLA

Andersen
Supplemental Figure S7 - Phylogenetics

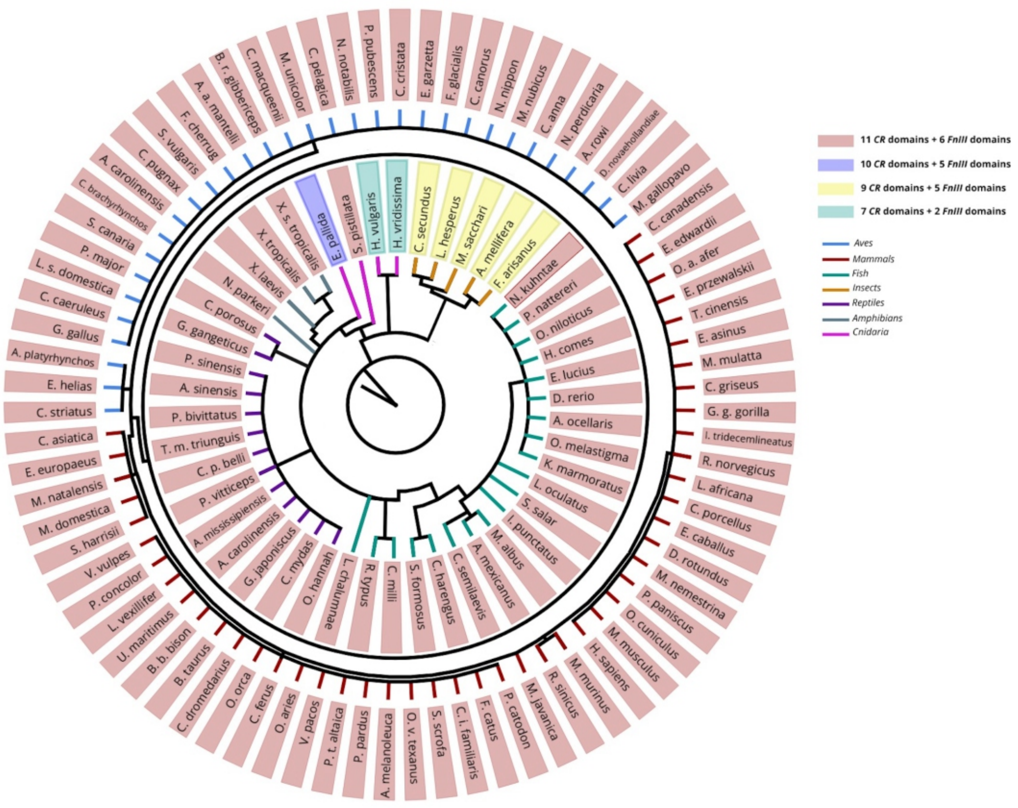

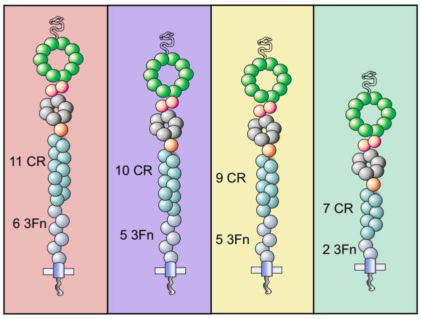

#### 10.a Supplemental Figure S7. Phylogenetic tree

Phylogenetic tree of 126 analyzed SORLA sequences available from the NCBI website, showing how most receptors contain 11 CR-domains and 6 3Fn-domains (pink), but also isoforms with only 10 CR-domains and 5 3Fn-domains (blue), 9 CR-domains and 5 3Fn-domains (yellow), or 7 CR-domains and 2 3Fn-domains exist. Domain compositions are summarized by schematics at the bottom following same color scheme as above.

### 11 Supplemental methods

**Phylogenetic tree generation:**

To create a comprehensive phylogenetic tree, the SORLA amino acid sequences from 126 of the >300 different species with known SORLA protein sequences were obtained from the NCBI website. Sequences were selected based on (1) having a sequence length of >1,000 aa, (2) obtaining a maximally diverse selection of animals within each species subclass (mammals, birds, fish, reptiles, insects, amphibians and cnidaria). Firstly, a multiple sequence alignment was created using the ClustalIW algorithm [227,228] and the phylogenetic tree was constructed based on this alignment with the Maximum Likelihood method using the MEGA7 software [229] to create a rooted tree. Subsequently, the phylogenetic tree was prepared and edited using GravitDesigner (Corel Corporation, Alludo).

**Conservation analysis: alignment of 40 representative SORLA sequences**

To determine semi-quantitative conservation of any residue within the human SORLA sequence across species, we aligned 40 representative sequences (**Supplemental Information 9**). The amino acid sequences were obtained from the NCBI website and an alignment was created using the ClustalIW algorithm [227,228]. Subsequently, the alignment was edited using Jalview software [230] to visualize consensus sequence and to highlight amino acids with 100% conservation (blue). This alignment reveals whether an introduced amino acid by novel SORL1 genetic variants is present in SORLA from other species.

### 12 Putting the pieces together - the conformational space

*Elongated or compact - Flexible or rigid structures?*

Thus far, all attempts to solve the structure of the full SORLA extracellular region have failed. Therefore, it is currently unknown how all the presented domains of the full-length SORLA protein fold relative to each other. In accordance with the different SORLA functions, the protein structure may also adopt different conformations. SORLA engages in the binding and sorting of cargo through cellular compartments, each with distinct pH levels and capacity to modify post-translationally attached *N*-glycosylations and perhaps phosphorylations [162].

Schematic illustrations of the SORLA molecule often show each domain lined up one after the other, extending to a predicted ~700 Å (=70 nm) (**Supplemental Figure S8a**). Such a linear conformation may be relevant at the plasma membrane if SORLA uses its VPS10p-domain to scavenge extracellular ligands. Or, since the receptor can locate to the postsynapse, the elongated SORLA-conformation could reach out across the synaptic cleft to binding partners at the presynapse. However, when SORLA is located in tubular extensions of the endosome (described to have a diameter of only 20-50 nm [231]), it is highly unlikely that SORLA will adopt this conformation [211]. Here, SORLA may adopt one or more compact conformations, similar to what has been found for LDLR and integrins [232-234].

The 3Fn-domain region may behave as a single compact structure, connected with the two β-propellers by the flexible CR-cluster. The region of the 3Fn-domains may be described *as a single and rather solid unit* [235]. In many receptor proteins the consecutive 3Fn-domains have a "*rod-like*" structure, with an angle between two neighboring 3Fn-domains close to 180 degrees: if all six 3Fn-domains of SORLA adopt this rod-like structure, the total structure would span ~210 Å. Two neighboring 3Fn-domains can also "*tilt*" with an angle typically around 120 degrees [154]: if all six SORLA 3Fn-domains adopt this "*wiggling*" structure they span only ~140 Å (**Supplemental Figure S8b**).

It is still unclear whether the two β-propeller domains in SORLA have a fixed or flexible orientation relative to one another. The two adjacent β-propeller domains may behave as a combined rigid structural block, which would be in accordance with the rigid connection between tandem β-propellers observed in LRP5 and LRP6. In this scenario the 5 amino acids ($^{753}$PLAEE$^{757}$) around the 10CC-YWTD linker that separates the two propellers, would not function as a hinge. Nevertheless, the 10CC-domain position relative to the VPS10p-domain may change with pH as described [5], arguing that this model may be over-simplified.

In contrast, the sequence directly following the EGF-domain of SORLA may act as a first hinge region (**H1, Supplemental Figure S8b**), paralleling the EGF-domain of integrin (also of the 8 Cys-type) that act as a hinge region between larger rigid units [63]. The site between the end of the CR-cluster and beginning of the 3Fn-domains of SORLA may serve as a second hinge region (**H2**), similar to the hinge region in LDLR that is found after its CR-cluster [102,232,236]. Interestingly, this second hinge would then locate next to the extremely long linkers in the SORLA CR-cluster and allow for substantial structural flexibility for this region. These two hinge regions might bring the receptor from a very "*elongated*" fold toward a more "*compact*" form (**Supplemental Figure S8b; Fig. 1**). With such a flexible CR-cluster it is unclear whether the YWTD- or the VPS10p-domain propeller is closest to the cell membrane (**Supplemental Figure S8**). However, most VPS10p-domain receptors have the VPS10p-domain close to the membrane (**Fig. 2**), suggesting that this a preferred position that may relate to shared functionalities.

*Monomer, dimer – or polymer*

The ability of SORLA to bind, traffic, and release cargo may depend on its flexibility but also on possible dimer formation. SORCS1 and SORCS2 exist mainly as a dimer; when the PKD-domains C-terminal to their VPS10p-domains interact, the top-face of their β-propellers are available for ligand binding, important for cell surface signaling [237,238].

When sortilin is internalized in endosomes it dimerizes upon the pH-drop, leading to ligand-release [239-241]. With the drop in pH, the side chains of a stretch of His residues on the top face of the VPS10p-domain β-propeller become protonated, leading to a conformational change of the VPS10p-domain loops, allowing dimerization at the top faces between two propeller-domains, enabling ligand displacement. It is unlikely that identical dimerization mechanisms occur for the SORLA VPS10p-domain because the SORLA sequence does not include any PKD-domains that drive dimer formation of SORCS1 and SORCS2. Neither does the SORLA VPS10p-domain sequence hold the His residues responsible for pH-dependent rearrangements in sortilin, nor the residues in the loops important for the direct contacts between the two VPS10p-domains [240,242]

However, nearly all known receptor-like molecules containing 3Fn-domains have the capacity to form dimers [148], and also SORLA dimerization can be mediated by its 3Fn-domains [24] (**Supplemental Figure S8**). Similar to SORLA, the extracellular region of growth-hormone-, prolactin-, and insulin receptors have 3Fn-domains that are located close to the plasma membrane and their activity is crucially associated with dimer conformation in the presence of bound ligands. The dimerization by 3Fn-domains of Tie1 and Tie2 receptors, from the tyrosine kinase family, is crucial for its activation [243]. While more studies are necessary to explore the physiological role of 3Fn-domain-mediated SORLA-dimerization, it is interesting to notice that several key ligands of SORLA (e.g. BDNF, APP, TrkB, HER2/3, GLUA1) exist in equilibrium between mono- and dimeric structures [244-246]. In contrast to the other sortilins, that only contain a single site of dimerization, the ability of a SORLA protein to bind another SORLA protein by either it's VPS10p-domain or its 3Fn-domains, suggest that also larger polymers of SORLA receptors may form [24] (**Supplemental Figure S8).**

While members of the LDLR family are mainly monomers, they also release ligands upon endosome internalization [102,233]. Here, the pH drop in endosomes leads to protonation of several histidine residues of the LDLR YWTD β-propeller. At the cell surface, LDLR has an elongated conformation and ligands bind to the CR-domains. Upon entering the endosome, the receptor adopts a more compact form: the β-propeller becomes protonated, and serves an alternative binding partner for the CR-domains which leads to displacement of the internalized ligand [232,247]. The released ligand is free for transport to lysosomal degradation while the receptor —in its closed conformation— is recycled through the acidic environments back to the cell surface, where it may once again adopt its open conformation, allowing CR-domains to interact with new ligands [248]. In contrast with LDLR, several cargo ligands for SORLA (incl. APP and HER2) have the highest affinity for SORLA-binding at *low* pH [70,249], allowing SORLA to escort ligands *out* of the endosome to either the TGN or to the cell surface [246,249] This requires efficient ligand binding *within* the endosome. Notably, the SORLA YWTD β-propeller sequence does *not* contain the His residues necessary for the pH-triggered release of ligands from the CR-domains [23]

*Support of "hidden" disulfides*

Based on the presented domain boundaries as defined by sequence alignments and expected domain folds, we present a 'compact' schematic diagram of SORLA, including all its 2,214 amino acids while taking into account the suggested presence of disulfides and β-strand secondary structures (**Fig. 1**). This diagram provides support for the presence of three disulfides that were not yet identified: (1) a disulfide between $C^{801}$ and $C^{816}$ at the β2-blade of the YWTD-domain, (2) a disulfide between $C^{1586}$ and $C^{1631}$ in the first 3Fn-domain, and (3) a disulfide between $C^{2101}$ and $C^{2108}$ in the sixth 3Fn-domain. (**Fig. 1**). In particular, the second suggested disulfide is separated by 45 amino acids in the primary structure, which can be appreciated only when taking the conserved topology of the 7-stranded 3Fn-domain scaffold into consideration. Both cysteine residues are predicted to point their side chain towards the hydrophobic interior of the 3Fn-domain β-blade sandwich as they occur "in-frame" with the alternating hydrophobic residues (see **Supplemental Figure S5**).

While many of these considerations about SORLA conformation await experimental validation, they may contribute to a better understanding of the nature of variant pathogenicity: SORLA folding, and thus activity, may not be compromised due to misfolding of single domains only, but also by damaging intramolecular interactions affecting bent and/or dimer/polymer conformations. We propose that next to variant effect prediction algorithms, the potential damagingness of genetic variants should be assessed based on features unique for SORLA: their position in the 3D alignments, and whether that position associates with disease in homologous proteins.

Andersen
Supplemental Figure S8 - conformational space

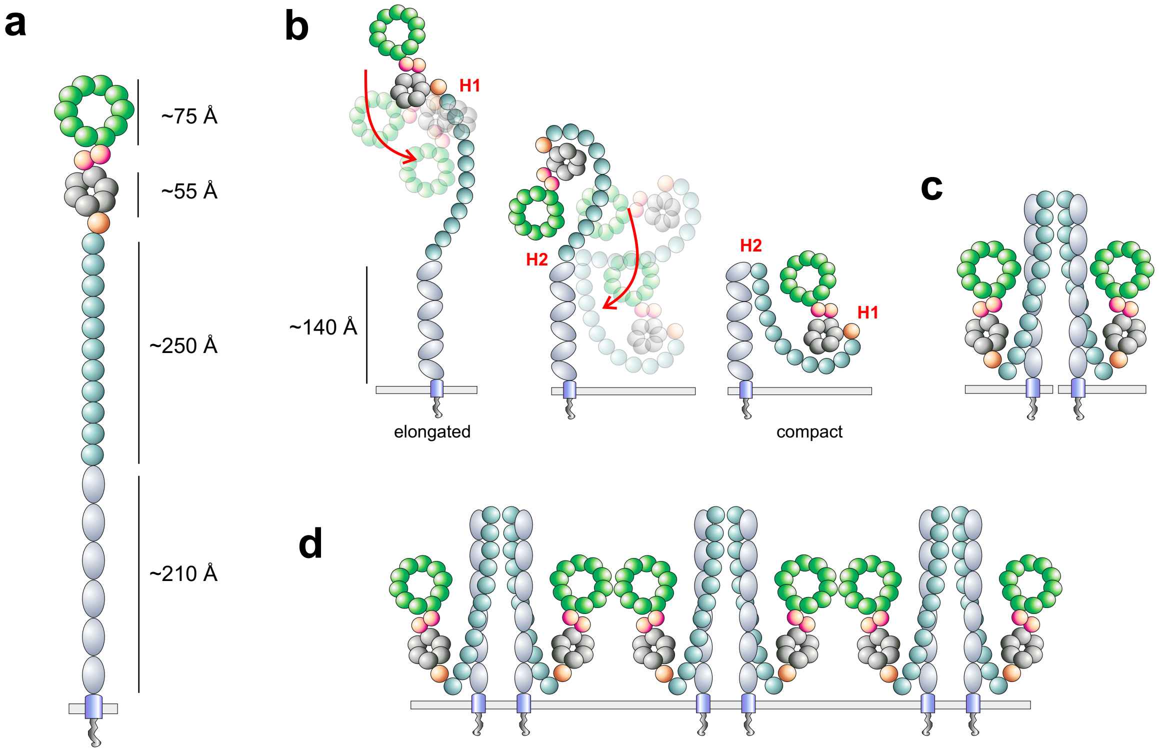

#### 12.a Supplemental Figure S8. The SORLA conformational space

**a**. Schematic of SORLA in an elongated conformation, spanning as much as ~700 Å if fully stretched, with indication of maximal expected size of the different receptor regions.

**b**. The speculated flexibility of the CR-cluster flanked by two potential hinge regions (H1 and H2) may contribute to several possible conformations, going from elongated to more compacts form. A tilt conformation between neighboring 3Fn-domains (angel set to 120) is indicated compared to the elongated possible conformation depicted in panel a.

**c**. Possible bend conformations with dimerization driven by the 3Fn-domains

**d**. Possible polymer formation of SORLA receptors due to the presence of sites in both the VPS10p- and 3Fn-domains that can facilitate dimerization and thus potential building of larger polymeric structures as previously suggested [24].
